# Putting the Asymmetric Response Concept to the test: modeling multiple stressor exposure and release in a stream food web

**DOI:** 10.1101/2024.07.02.601677

**Authors:** Annabel Kuppels, Helena S. Bayat, Svenja M. Gillmann, Ralf B. Schäfer, Matthijs Vos

**Affiliations:** Ruhr University Bochum, Faculty of Biology and Biotechnology, Theoretical and Applied Biodiversity Research, Bochum, Germany; Institute for Environmental Sciences, RPTU Kaiserslautern-Landau, Landau, Germany; Department of Aquatic Ecology, University of Duisburg-Essen, Essen, Germany; Research Center One Health Ruhr, University Alliance Ruhr & Faculty for Biology, University of Duisburg-Essen, Essen, Germany

**Keywords:** Degradation, Global warming, Heatwave, Multiple stressors, Recovery, Restoration, Tolerance

## Abstract

Communities in stream ecosystems often respond asymmetrically to increase and release of stressors, as indicated by slow and incomplete recovery. The Asymmetric Response Concept (ARC) posits that this is due to a shift in the relative importance of three mechanisms: tolerance, dispersal, and biotic interactions. In complex natural communities, these mechanisms may produce alternative outcomes through poorly understood indirect effects. To understand how the three mechanisms respond to different temporal stressor scenarios, we studied multiple scenarios using a stream food web model. We asked the following questions: Do groups of species decline as expected on the basis of individual tolerance rankings derived from laboratory experiments when they are embedded in a complex dynamic food web? Does the response of ecosystem function match that of communities? To address these questions, we aggregated data on individual tolerances at the level of functional groups and studied how single and multiple stressors affect food web dynamics and nutrient cycling. Multiple stressor scenarios involved different intensities of salt and temperature increase. Functional groups exhibited a different relative tolerance ranking between the laboratory and dynamic food web contexts. Salt as a single stressor had only minor and transient effects at low level but led to the loss of one or more functional groups at high level. In contrast, high temperature, alone or in combination with salt, caused the loss of functional groups at all tested levels. Patterns often differed between the response of communities and ecosystem function. We discuss our findings with respect to the ARC.

**Graphical Abstract:** 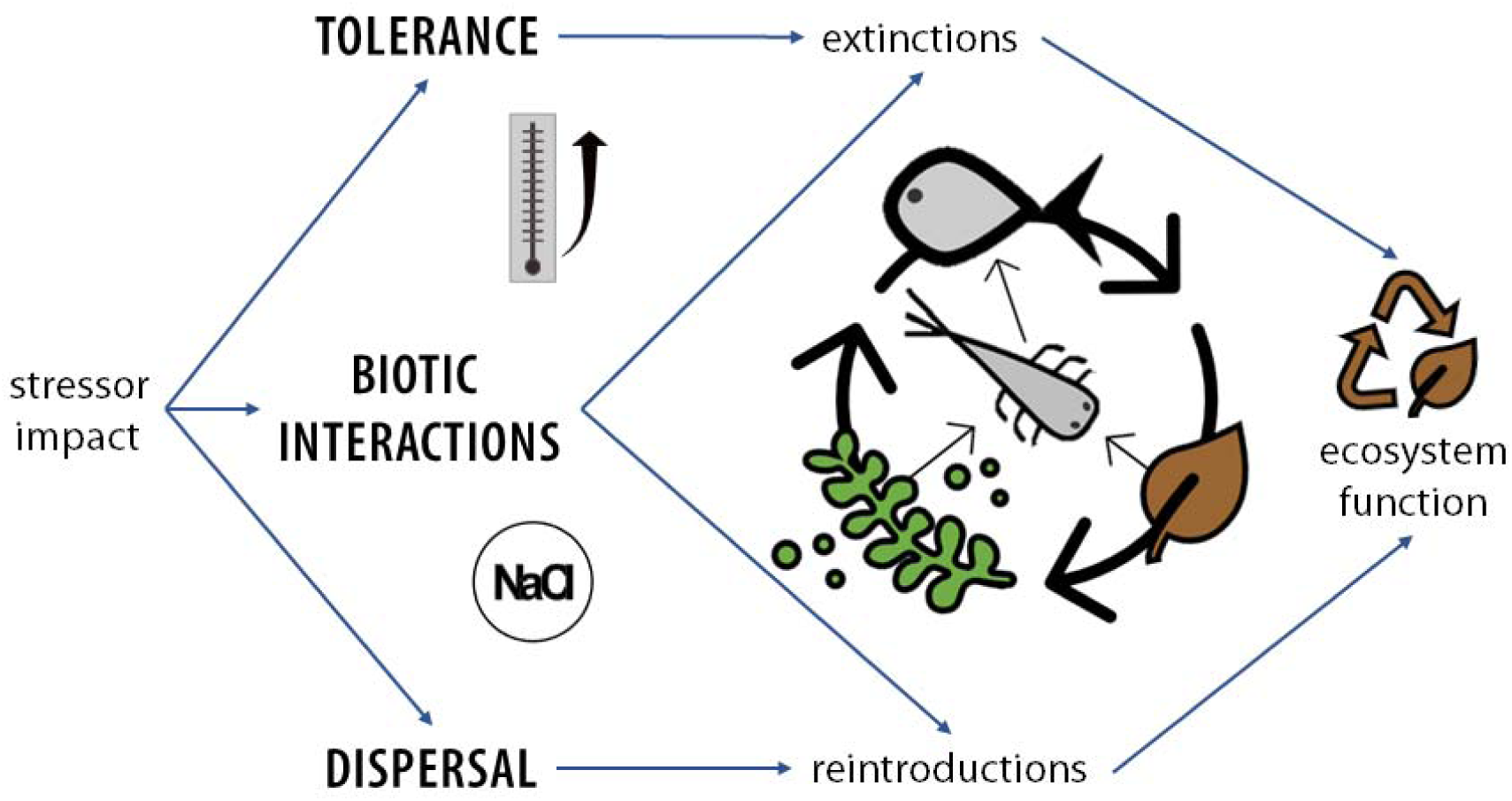

## 1. Introduction

### 1.1. General background

Multiple stressors pose threats to biodiversity and ecosystem function in aquatic and terrestrial ecosystems (Orr et al., 2020). Understanding multiple stressor effects is complicated by the diversity of stressors and the type, order and temporal range of intensities in which they may occur and co- occur (Jackson et al., 2021; Rillig et al., 2019; Speißer et al., 2022). Anthropogenic stressors are defined as any anthropogenic factor that causes a biological response to exceed its range of normal variation relative to reference conditions (Piggott et al., 2015; Straalen, 2003; Vos et al., 2023). Over the past decades, multiple stressor research has included nearly all taxonomic groups (Orr et al., 2020). However, we lack an understanding of how mechanisms operate across levels of biological organization, from individual responses up to the ecosystem level (Schäfer et al., 2023; Schneeweiss et al., 2023; Simmons et al., 2021).

Understanding the impacts of and recovery from stressors in the real world necessitates understanding the food webs that link members of ecological communities. In food webs, organism groups and functions are connected through direct and indirect interactions (Vanni and de Ruiter, 1996; Wootton, 1994). For example, responses of periphyton to stressor increase and release will not only depend on co-occurring responses in the grazer functional group but also on the levels of nutrients made available through nutrient cycling by bacteria, fungi, leaf litter shredders, and other detritivores. These are in turn differentially affected by invertebrate and fish predators. Biotic responses are therefore not only defined by competition and predation but are also affected by contributions to ecosystem function by some organism groups that other groups depend on (Vanni and De Ruiter 1996). This level of complexity tends to defy intuitive predictions of stressor effects and is difficult to study in the field or in the laboratory.

Models allow the investigation of mechanisms at this level of complexity. Theoretical ecology has a long history of using such models to answer general questions, such as whether increasing complexity increases or decreases the stability of food webs (Brose et al., 2003; de Ruiter et al., 1995; May, 1972; McCann et al., 1998; Neutel et al., 2007, 2002). To make full use of the power of models, they should be used as tools within a robust theoretical framework (De Laender, 2018; Vos et al., 2023).

The Asymmetric Response Concept (ARC) hereafter referred to as the ARC, outlines a theoretical framework for the ecological mechanisms governing response trajectories to stressors, before and after stressor release (Vos et al., 2023). According to the Asymmetric Response Concept, ecological trajectories of stressor effects are not necessarily mirror images of recovery trajectories. One key reason for these asymmetries is that different ecological processes dominate changes in community composition before and after stressors are released or reduced. Depending on how the processes of stress tolerance, dispersal and food web interactions play out over time, recovery may be fast or slow, complete or partial, result in novel communities and ecosystem states, or fail completely (Lake et al., 2007; Sarr, 2002; Vos et al., 2023). The canonical inference of the ARC dictates that tolerance best explains the extent of community change during stressor exposure, whereas dispersal and biotic interactions consecutively gain importance following stressor release (Vos et al., 2023).

Here we aim to contribute insight into the responses of stream food webs to multiple stressors using a stream food web model constructed on the theoretical basis set by the ARC. We focus on stream ecosystems because of their vulnerability to stressors and importance to many species, including humans, worldwide (Jackson et al., 2016; Lynch et al., 2023; Reid et al., 2019). Streams are subject to considerable restoration efforts that have only been partially successful (Jourdan et al., 2019; Muhar et al., 2016; Verdonschot et al., 2013; Vos et al., 2023). Real stream food webs can be divided into green, autotroph-based, and brown, detritus-based parts. These two are typically coupled by nutrient cycling and shared predators (Cordone et al., 2020; Evans-White and Halvorson, 2017; Mougi, 2020; Zelnik et al., 2022). The green part depends on nutrient remineralization. Nutrients from detritus and terrestrial leaf litter entering the system are made available to primary producers by bacteria, fungi, leaf shredders, and other detritivores (Moore et al., 2004; Wallace and Webster, 1996; Zou et al., 2016). Our model captures this general feature in its design and is based on classical equations for food webs with nutrient cycling (DeAngelis, 1992). The model captures essential, generalized features of a real stream food web, which may be revised according to local variability (cf. The River Continuum Concept in this context, see (Downs et al., 2002; Welcomme et al., 1989). In our analyses, the freely available nutrient quantity was used as a metric of ecosystem function. Nutrient quantity represents the efficacy of nutrient cycling by the brown part of the food web, whereby free nutrients are made available to the green part (Huang et al., 2018; O’Brien and Wehr, 2010). Re-colonization was investigated by introducing taxa into the system after the stressor phase. This allowed us to examine its impact on subsequent dynamics and recovery of function compared with controls that lacked such re-colonization. In real systems, re-colonization could occur by natural dispersal or by man-made species reintroductions (Dumeier et al., 2020). In our present work we study re-colonization as a particular aspect of dispersal by implementing reintroductions of functional groups and then evaluating whether these result in re-establishment.

We implemented temperature as a key stressor in our analysis because it may strongly affect stream communities (Leach et al., 2022) and is expected to significantly increase in the future, both gradually and in the form of summer heatwaves (Calvin et al., 2023; Hansen et al., 2012; IPCC, 2021). Salinization is similarly becoming a global problem (Kaushal et al., 2021). It is caused by sewage input and the effects of multiple land uses, including agriculture, mining, and deforestation (Cañedo- Argüelles et al., 2013). The use of road salt contributes to salinization (Kaushal et al., 2005). Prolonged summer droughts associated with climate change may additionally lead to salinization through groundwater incursions in coastal systems (IPCC, 2021; Venâncio et al., 2023).

To connect the functional groups in the food web model to stressor effects in a realistic way and simultaneously study the interplay between tolerance and biotic interactions in shaping effects, we aggregated individual tolerance data to the stressors temperature and salinity and integrated this data into various stressor scenarios. Because different stressors can impact streams alone or simultaneously, we implemented model scenarios in which stressors act singly or at the same time.

In this study, we address the core question of how ecological mechanisms, with a focus on tolerance, re-colonization and biotic interactions, shape food web responses to stressor exposure and release. We explore the fate of nutrient cycling following stressor-induced extinctions and study how re- colonization affects the recovery rate of this process. In line with the ARC framework, we hypothesize that 1) the least and most sensitive groups exhibit the weakest and strongest responses to stress, respectively, so that relative individual tolerance is related to the magnitude of effect during the stressor phase. In terms of food web simulations, this means that more sensitive groups should decline more strongly and reach extinction more rapidly under stressor increases. We further hypothesize that 2) a nearly symmetric (rapid and full) recovery of nutrient cycling occurs following successful reintroductions, but that asymmetric responses ensue (involving slow and/or incomplete recovery) when reintroductions fail or do not occur.

### 1.2. Theoretical framework

We address these hypotheses in view of our broader aim to refine the Asymmetric Response Concept and make it more applicable to real ecological communities. According to the ARC, there are at least 5 possible outcomes for ecological response to stressor exposure and release: 1) rubber band, 2) broken leg, 3) partial recovery, 4) no recovery, and 5) new state (**Figure 1**) (Vos et al., 2023). Here we operationalize these 5 response outcomes as follows for our model analysis: Outcome 1, ‘rubber band’, is a (practically) *symmetric* response before and after release from a stressor or stressor combination, both in terms of the initial and final state (recovery is complete) and in terms of trajectories (the time to depart from and return to an original state are identical). In our food web analysis, we refer to a response as *symmetric* (outcome 1) if full recovery occurs within 2 years of the stressor release. We selected this 2-year window to account for a possible re-set of the system by annually pulsed leaf litter input (see next section for details). Outcome 2 has been called the ‘broken leg’ model, as recovery is complete but takes longer than the time to depart from the original state (Figure 1). Outcome 2 combines symmetry in the initial and final states (complete recovery occurs) with *asymmetric* temporal trajectories. In our food web analysis, we refer to outcome 2 when recovery occurs between 2 and 8 years after stressor release. Outcome 3, ‘partial recovery’ and outcome 4, ‘no recovery’ are asymmetric in terms of both the initial and final states and the trajectories before and after release from stressors; recovery remains incomplete or does not occur at all. In our food web analysis, we refer to outcome 3 when the difference to the non-stressed system decreases over time until a plateau is reached (Figure 1). Outcome 4 implies that no recovery occurs after stressor release (Figure 1). Outcome 5, called ‘new state’, is also fully asymmetric. After stressor release, the system departs even further from its original state. In our food web analysis, outcome 5 implies that the difference to the non-stressed reference system increases after stressor release and then either stabilizes or continues to increase. We used the above classifications to link our simulation patterns for different stressor exposure and release scenarios directly to the ARC.

**Figure 1.**
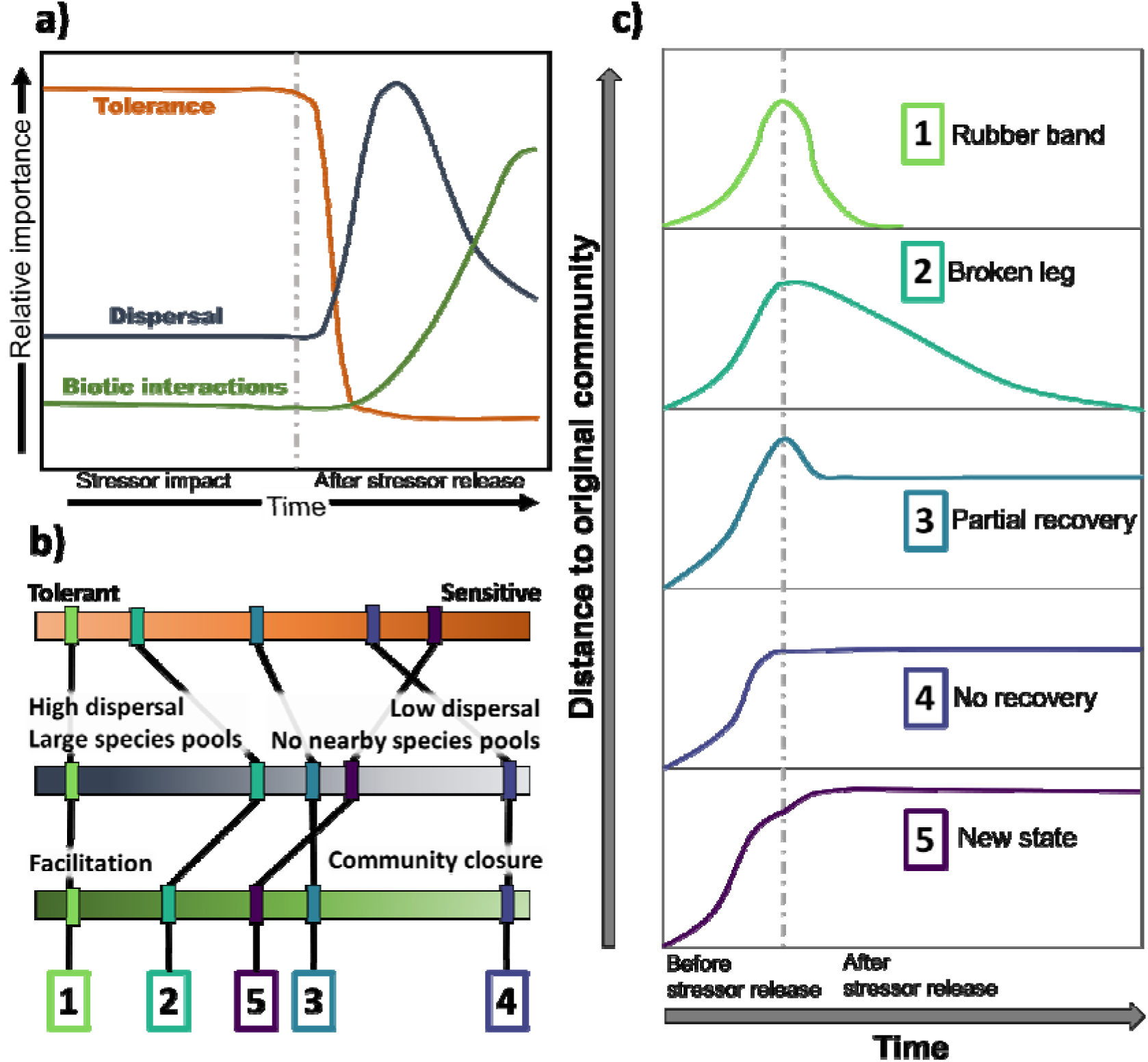
The three ecological mechanisms that change in relative importance after stressor release as postulated by the Asymmetric Response Concept (**a**). Variations in these mechanisms lead to one of five outcomes following stressor release (**b**, **c**). Example conditions for each ecological mechanism that can lead to alternative outcomes are shown in (**b**). Five possible outcomes linking community response before and after stressor release to the Asymmetric Response Concept (ARC): 1) rubber band, 2) broken leg, 3) partial recovery, 4) no recovery, and 5) new state (**c**).

## 2. Methods

### 2.1. Freshwater food web model

We developed a stream food web model consisting of functional groups connected by biomass flows through feeding, growth, and mortality described by ordinary differential equations to model food web dynamics and nutrient cycling (see **Appendix B** for equations). The modeling approach is based on a basic food web model (DeAngelis, 1992) that includes nutrient cycling. We extended the model with several functional groups to create a first approximation model that captures the essential, generalized features of a real stream food web (**Figure 2**). The model consists of a green food web fed by periphyton and a brown food web fed by detritus, connected by nutrient cycling and predators. The green part includes nutrient, periphyton, and grazer groups, whereas the brown part includes plant detritus, animal detritus, coarse particulate organic matter (CPOM) derived from terrestrial leaf litter, fine particulate organic matter (FPOM), (dissolved) organic matter (OM), a leaf shredder, and other types of detritivore, fungi, bacteria, dead fungi, and dead bacteria (**Figure 2**). The invertebrate and fish predator link the green and brown parts as they feed on both. We implemented trophic interactions between functional groups by functional responses that were adjustable so that they could range from Type 1 to Type 3.

**Figure 2.**
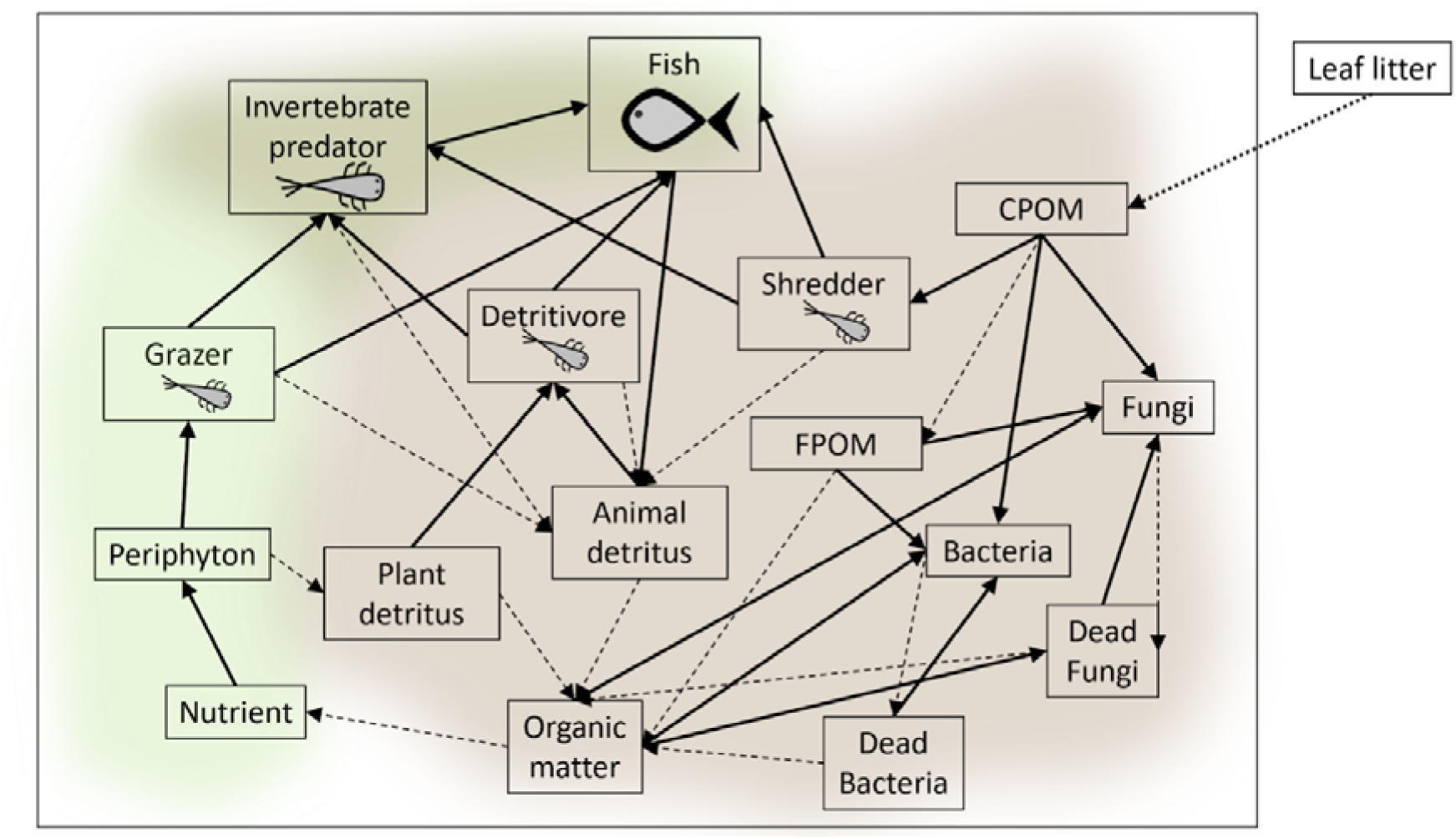
The food web model is composed of 16 functional groups divided into green and brown parts that are linked by predation (i.e. feeding by the invertebrate predator and fish predator). Leaf litter represents an external input variable to the food web and the annual terrestrial CPOM influx (dotted arrow). Solid arrows denote feeding relations, and dashed arrows indicate the flow of dead material and material breakdown.

All living functional groups have a natural mortality rate and may experience additional mortality when exposed to single or multiple stressors. Parameters were chosen to create a stable food web system in which all functional groups persist for more than 50 years. The simulations analyzed involve a period of 10 years. We used a Holling type III functional response as the basis for our simulations because it stabilizes consumer-resource systems and consumers tend to reduce foraging effort at very low resource densities (Kalinkat et al., 2023, 2013; Majumdar et al., 2022). Our choice of functional response additionally allowed the sub-webs and modules of this food web to be stable, enabling us to unravel mechanisms in relation to structure. All equations, parameters, and coefficients related to the functional groups can be found in the supplementary information (**Appendix B**).

In this study, the model simulations were run in daily time steps for 10 years. An annual leaf litter input, as a CPOM influx, marked the beginning of each simulation year. The extinction threshold for living compartments was set to 0.001 mg biomass per liter, representing a minimum viable density. When the density for a functional group fell below the threshold, the density was set to zero, resulting in extinction. Fungi and bacteria did not have a threshold because microbial populations can regrow from a single cell.

### 2.2. Individual tolerance

To implement tolerance to stressors in the food web, we retrieved data representing a typical central European stream in a highly urbanized region. We used a taxa list of freshwater macroinvertebrates resulting from field sampling conducted in 2021 in the Boye catchment (Western Germany) to query the literature and relevant databases for tolerance data at the species and genus level (Gillmann et al., 2023). Details of the catchment are described in Gillmann et al. (2023). A taxa list for fish species in this catchment was obtained from monitoring conducted by the Emschergenossenschaft Lippeverband (EGLV), a German water management organization (Jacobs, 2017). The macroinvertebrate taxa were first divided into feeding groups based on the classification of Moog 1995, obtained from freshwaterecology.info (Moog, 1995; Schmidt-Kloiber and Hering, 2015). These feeding groups were then adjusted to match the functional groups of the food web model. Two approaches were taken to translate tolerance data into death rates relevant to stressor scenarios run in the model.

For thermal tolerance, a linear increase in the death rate from optimum to maximum temperature was assumed based on the shape of the temperature performance curves (**Appendix A, Figure A1**). An optimum temperature of 15°C was assumed following Sundermann et al. (2022) and Tomczyk et al. (2022). Maximum values per group were determined as the average of all component taxa at the species and genus level. Data at the genus level were restricted to taxa from temperate climates. Excess mortality for each stressor scenario was derived in this way for each invertebrate group and the fish in the model. The equations and details of the calculations can be found in **Appendix A**.

For salinity tolerance, insufficient data were available at the species and genus level to justify calculating an average per functional group. Instead, excess mortality was determined for all invertebrate groups and fish separately. A species sensitivity distribution was fit for aquatic invertebrates and another for fish using data queried from the StandarTox database (Scharmüller et al., 2020). The fraction of affected organisms at the chosen salinity level was calculated for each stressor level to obtain an excess mortality rate for fish and each invertebrate group (**Appendix A, Figure A6, A7**).

### 2.3. Stressor scenarios

To implement stressor scenarios in the model, excess mortality at the selected stressor level was added to the natural death rate of the affected functional group. Excess mortality at a specified temperature or salinity level was derived from tolerance data as described above. For temperature, each fish and invertebrate group had a different excess mortality corresponding to differences in tolerance. For salinity, all four invertebrate groups (shredder, grazer, detritivore and invertebrate predator) had the same excess mortality at each salinity level, which differed from the excess mortality derived for fish. For multiple stressor scenarios, excess mortality rates were summed, assuming no stressor interaction at the functional group level. The effect strength for each scenario was calculated by summing excess mortalities for all functional groups and standardizing these to a value between 0 and 1 (**Table 1**).

**Table 1.** Description of the stressor scenarios run using the model.

| Scenario type | Number | Salinity level | Temperature level | Effect strength |
| --- | --- | --- | --- | --- |
| Single | 1 | 600 mg/L | NA | 0.05 |
| Single | 2 | 1500 mg/L | NA | 0.2 |
| Single | 3 | NA | 18.5°C | 0.21 |
| Single | 4 | NA | 25°C | 0.61 |
| Multiple | 5 | 600 mg/L | 18.5°C | 0.26 |
| Multiple | 6 | 1500 mg/L | 18.5°C | 0.41 |
| Multiple | 7 | 600 mg/L | 25°C | 0.66 |
| Multiple | 8 | 1500 mg/L | 25°C | 0.81 |
| Single | 9 | 200 mg/L | NA | 0.004 |
| Single | 10 | NA | 22°C | 0.42 |
| Single | 11 | 5300 mg/L | NA | 0.61 |

All stressor phases were implemented for 14 days in the middle of the third simulation year, from day 912 until day 925. We chose this time window because the interannual dynamics had been established by year three and the intra-annual dynamics of functional groups following leaf litter input had also stabilized. Stressors were tested at two levels, each alone and in combination, for a total of seven single stressor scenarios and four multiple stressor combinations. All 11 stressor scenarios tested are given and numbered in **Table 1**.

### 2.5. Re-colonization

Reintroduction scenarios were performed following stressor scenarios where extinctions occurred by adding a fixed density of a functional group into the system at one time step. Reintroductions were tested at different time steps, ranging from one day after the end of the stressor phase (day 927) until the next CPOM input 169 days later (day 1095). If an extinction occurred after the stressor phase then the reintroductions were tested from one day after the last extinction until the next CPOM input. Four reintroduction densities were chosen based on the density of the group before the start of the stressor phase: 1) the pre-stressor density, 2) + 10%, 3) -10% and 4) -90%. When more than one group was reintroduced, the reintroductions were conducted either simultaneously or sequentially. In these cases, the focus was on one group, which was reintroduced with the 4 densities mentioned above, whereas all other groups were reintroduced with their pre-stressor densities. If the groups were sequentially reintroduced, the focal group was either the first or the last to be reintroduced. All other groups were introduced simultaneously in this case.

### 2.6. Data analysis

Tolerance rankings of affected invertebrate groups and fish were determined based on experimental data from laboratory experiments as described in 2.2, hereafter called laboratory context, and for the dynamic food web context based on the relative changes of the groups during and after the stressor phase. Rankings within the food web context *during the stressor phase* were determined by comparing density decreases under stressor impact for each group during the first half of the stressor phase. Density decreases were calculated relative to densities occurring in the same period without a stressor phase (in a control simulation) because cyclical decreases in density are a feature of the dynamic model. Rankings *after the stressor phase* were based on the number of days to extinction within a scenario. Groups that took longer to become extinct were deemed more tolerant. Spearman’s correlation coefficients were determined between rankings using R version 4.2.1 (R Core Team, 2022). A detailed description of the calculations can be found in **Appendix A**. Data processing and the creation of figures were conducted in R using the packages tidyverse, ggplot2, viridis, and patchwork (Garnier et al., 2023; Pedersen, 2023; R Core Team, 2022; Wickham, 2016; Wickham et al., 2019).

## 3. Results

### 3.1 Baseline scenario dynamics

In a typical simulation year, annual leaf litter CPOM input leads to an increase in the brown fraction of the food web, which decreases over the year as CPOM is broken down into FPOM, dissolved OM, and remineralized into free nutrients. This nutrient feeds the green food web fraction over the year as the brown part declines (**Appendix C, Figure C1**). The biomass of the focal functional groups shredder, detritivore, invertebrate predator, and fish increased immediately after the CPOM input because of the increases in food sources. Once nutrients become available through the brown food web, the grazer biomass increases (**Appendix C, Figure C1**). Following the initial increase, the biomass of all focal groups decreased gradually over the year (**Appendix C, Figure C1**).

### 3.2. Stressor effects on functional community composition

#### 3.2.1. Single and multiple stressors

The chosen salinity levels resulted in smaller effects than the chosen temperature levels (**Figure 3a, b, c, Appendix C, Figure C2**). Extinction only occurred under high salinity stress (scenarios 2, 11). In scenarios 9 and 1 the effects only led to temporary changes in the relative abundance of functional groups that returned to baseline levels in less than 2 years for scenario 9 (200 mg/L salt) and after several years for scenario 1 (600 mg/L salt) (**Figure 3a, g, d, Appendix C, Figure C2**). According to the ARC classification scenario 9 showed a fully *symmetric* response, following outcome 1, rubber band. The small change in densities caused by the stressor rapidly reverted in less than 2 years (**Figure 3g**). In Scenario 1 recovery took longer. It showed an *asymmetric* response pattern that followed ARC outcome 2, broken leg, as recovery took several years but was nonetheless complete at the end of the 10-year period (**Figure 3d**). The higher salinity scenario 2 (1500 mg/L salt), on the other hand, led to loss of the shredder, resulting in permanent changes in abundance of the remaining functional groups (**Appendix C, Figure C2**). This response pattern was *asymmetric*, following ARC outcome 3, partial repair. The densities of the grazer and detritivore came close to their original baseline values, but the densities of the invertebrate predator and fish remained different (**Appendix C, Figure C2 s2**). Reintroduction of the shredder was successful and led to a return of all groups to their original densities. The recovery showed an *asymmetric* response pattern which, according to the ARC, followed outcome 2, broken leg, as recovery was complete but took several years (**Appendix C, Figure C3 s2**).

**Figure 3.**
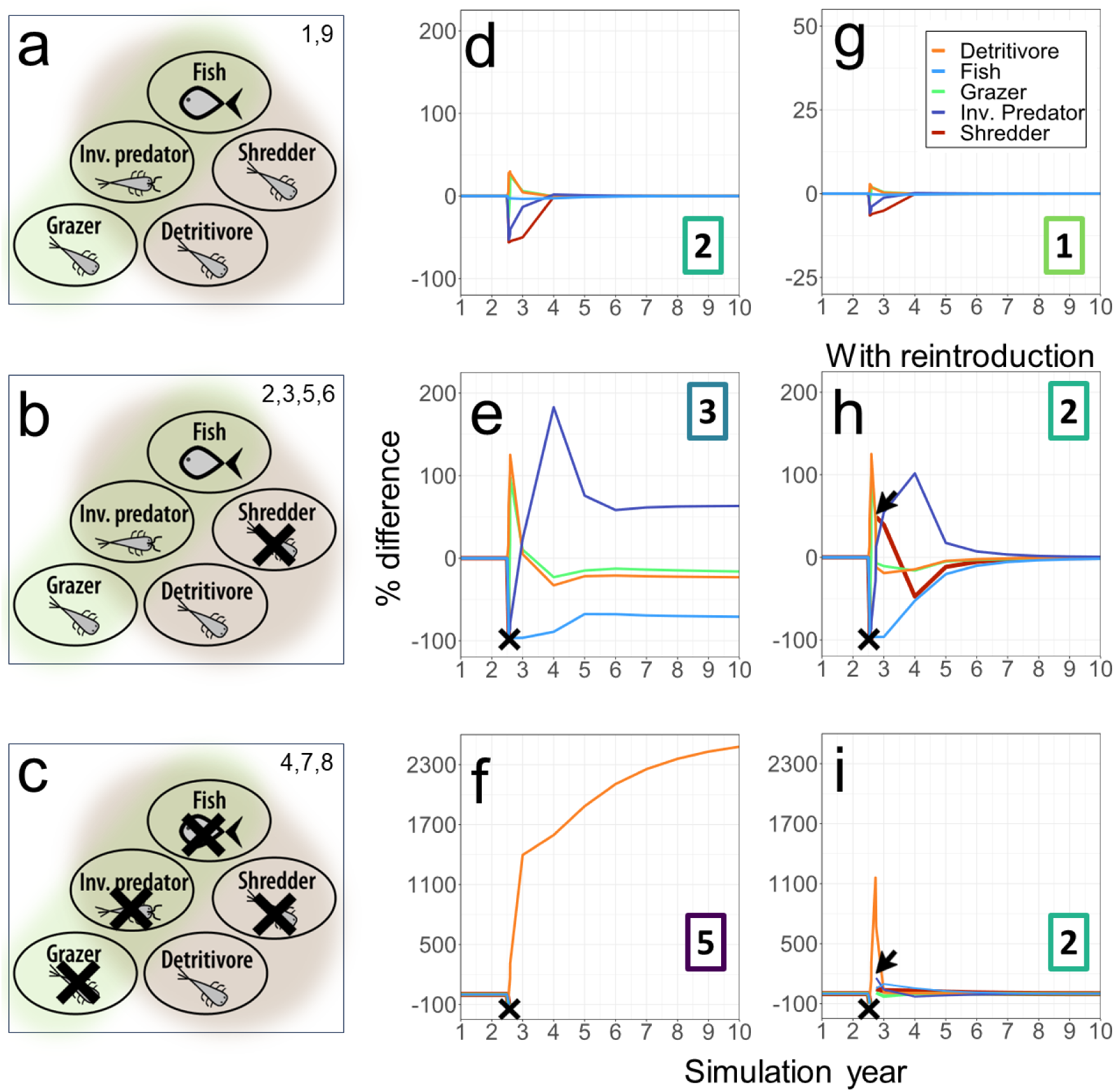
Single and multiple stressor scenarios resulted in either **a**) no extinctions, **b**) extinction of the shredder, or **c**) four extinctions (shredder, fish, grazer, invertebrate predator). Extinctions are marked with a black **x**. The numbers within **a**–**c** represent the corresponding scenario numbers in **Table 1**. Numbers in the top right of **d**–**i** represent the outcomes according to the ARC (Figure 1c), responses are exemplarily depicted. Changes in abundance occurred in absence of extinctions (**d, g**). When no extinctions occurred, all affected groups returned to the undisturbed baseline (**d, g**). The scenarios with four extinctions resulted in a large increase in the detritivore population, which was resilient across all scenarios (**f**). With reintroduction, marked with a black arrow (**h, i**), a return to baseline occurred for all affected groups (**h**, **i**). Depicted reintroductions were conducted at time step 1000 (towards the end of year three); the end of the stressor phase occurred at time step 925.

Temperature stress alone led to loss of the shredder at 18.5°C (scenario 3, **Appendix C, Figure C2 s3**) and extinctions of the shredder, grazer, invertebrate predator, and fish at 25°C (scenario 4, **Appendix C, Figure C2 s4**). These scenarios resulted in lasting changes in the abundance of the remaining functional groups when no reintroductions occurred (**Figure 3e, f**). The response pattern of scenario 3 (18.5°C) followed ARC outcome 3, partial repair. For scenario 4 (25°C), on the other hand, the response pattern followed outcome 5, new state. The difference between the recovery trajectory and non-stressed dynamics increased until the end of the simulated period (**Figure 3f**). When lost groups were reintroduced, densities returned to their original values (**Figure 3h, i, Appendix C, Figure C3**). In both scenarios the response pattern was *asymmetric* and followed ARC outcome 2, broken leg, as recovery took several years.

Salinity levels added to temperature levels did not lead to additional extinctions. Temperature dominated the overall effects, and single temperature stressor scenarios resulted in very similar effects to multiple stressor scenarios. Both scenarios 5 and 6 that combined the 18.5°C temperature stressor with salinity (600 and 1500 mg/L) led to the extinction of the shredder when implemented simultaneously for 14 days (**Figure 3e**). These multiple stressor scenarios showed a similar *asymmetric* response following ARC outcome 3, partial repair (**Appendix C, Figure C2 s5, s6**). Both scenarios 7 and 8 that combined the 25°C temperature stressor with salinity (600 and 1500 mg/L) led to loss of the shredder, grazer, invertebrate predator and fish, when implemented simultaneously for 14 days (**Figure 3c, f**). No differences in terms of functional group dynamics were observed between the two 25°C multiple stressor scenarios (**Appendix C, Figure C2 s5, s6**). In both cases, the biomass of the detritivore continued to increase over the years (**Figure 3f**). Both scenarios led to an *asymmetric* response pattern following ARC outcome 5, new state, as the difference between the recovery trajectory and non-stressed dynamics kept increasing until the end of the studied 10-year period. The system returned to its original dynamics and densities when all lost species were reintroduced. This was the case in all multiple stressor scenarios (5, 6, 7, 8, **Figure 3h, i**). All of these showed an *asymmetric* response following ARC outcome 2, broken leg.

### 3.2.2. Tolerance rankings for invertebrate groups and fish

Tolerance rankings differed with the stressor scenario and time; rankings for the 25°C temperature scenarios are depicted in **Figure 4**. Although none of the Spearman rank correlation coefficients were statistically significant due to the small sample size, the pattern in correlation strength confirmed expectations. Relative tolerances in the laboratory context were more similar to those during the stressor phase than after. All three tolerance rankings from the dynamic food web context were more similar than the laboratory context-based ranking. Notable differences between tolerance derived from the laboratory and food web context under stressor impact were exemplified by the detritivore, which was a close second in sensitivity under the individual tolerance ranking, yet resoundingly tolerant in the food web ranking (**Figure 4**). A similar but opposite pattern was observed for the shredder, which was average in terms of laboratory-based tolerance but sensitive in the food web context (**Figure 4**).

**Figure 4.**
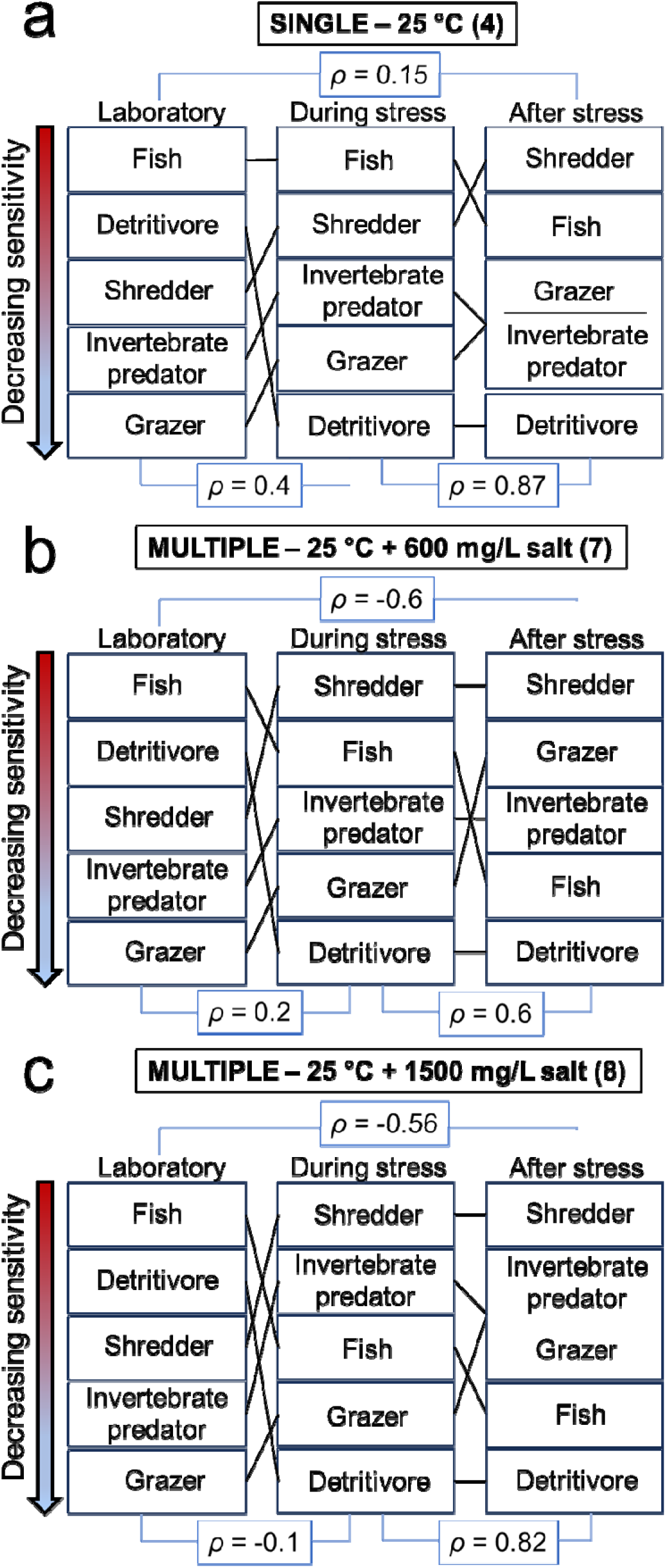
Temperature tolerance rankings of invertebrate groups and fish according to individual tolerance data from the laboratory context, contrasted with rankings derived from the food web context during and after the stressor phase, for high stress due to a single (**a**) or multiple (**b, c**) stressors; scenario number in parentheses. Laboratory data for a single species were averaged by functional group and then ranked. The ranking during stress was derived according to the relative density decrease rate during the first half of the stressor phase, and the ranking after stress was determined using the days to extinction (see Appendix A). Single stress (**a**) reflects the 25°C scenario, whereas multiple stress reflects lower and higher levels of the secondary stressor; 25°C and 600 mg/L salt (**b**) and 25°C and 1500 mg/L salt (**c**). Spearman’s *ρ* is marked for pairs of rankings. While none of the p values were significant for Spearman’s *ρ*, rankings within the food web context were more similar than to the laboratory rankings. As the secondary stressor level increased, correlation between laboratory and community metrics decreased.

## 3.3. Recovery

All groups that were simultaneously reintroduced after local extinction were successfully re-established at all tested reintroduction times and densities. Thus, full recovery in terms of community composition was possible for all stressor scenarios at all times after stressor release. The sequential reintroduction of several groups was also always successful in all scenarios, regardless of the order of reintroduction (**Appendix C**). The phenomenon of community closure (Lundberg et al., 2000), when the re-establishment of taxa is blocked by biotic interactions of the post-stressor community, was not observed. Note that community closure was not expected in these simulations because periphyton, bacteria, and fungi were not expected or modeled to decline under the stressor levels tested, and no mechanisms (such as priority effects) that would likely cause community closure were implemented in the parameterization.

## 3.4. Nutrient cycling

The 600 mg/L salinity stressor alone resulted in nutrient perturbations within 1% of the undisturbed system, whereas the 1500 mg/L salinity stressor resulted in greater changes that increased over time (**Figure 5**). Effects on nutrient amount were indistinguishable after the stressor phase between high temperature (25°C) single and multiple stressor scenarios and were moderate for mild (18.5°C) temperature stressor scenarios (**Figure 5**).

**Figure 5.**
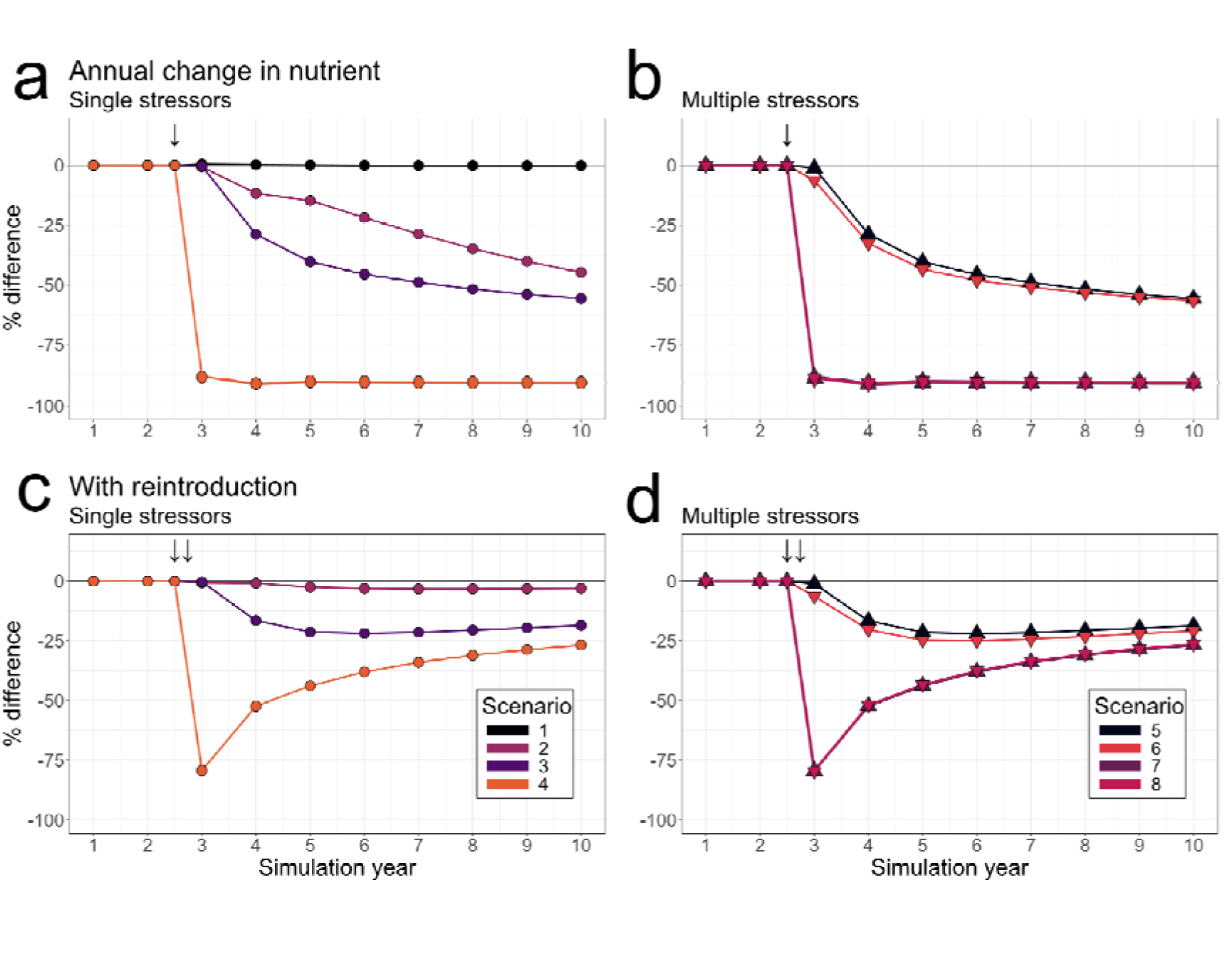
Fraction of free nutrients over time following single and multiple stressor scenarios with reference to the unstressed system for scenario 1-8. Upper row represent single stressor scenarios (left) and multiple stressor scenarios (right) with no reintroduction; lower row represent stressor scenarios followed by the reintroduction of extinct groups 75 days after the end of the stressor phase (not present when no extinctions occurred). Arrows mark the start of the stressor phase and the time point of reintroduction; reintroduction always occurred after the stressor phase.

Interestingly, changes in nutrient quantity relative to the reference community persisted for at least 7 years following stressor release, even when no extinctions occurred (**Figure 5**). The extinction of the shredder led to a pronounced difference in nutrient quantity compared with the reference community. The changes in nutrient quantity in scenarios 2, 3, 5 & 6 further increased in the following 7 years. While the changes in scenario 2 were constant, they slowed down in scenarios 3, 5 and 6 (**Figure 5**). The overall pattern is that scenarios 2, 3, 5 and 6 continued to move in the direction of a new state at the ecosystem level. The reintroduction of the shredder mainly resulted in a slow return toward baseline levels, reaching within 20% of baseline levels by year 10, thus showing partial repair. In scenario 2, the reintroduction only led to a reduced deviation, rather than to recovery. This kept the nutrient level within 3% of the baseline (Figure 5). With extinctions of several groups, on the other hand, the nutrient quantity decreased and stabilized in a new state much lower than the reference (scenarios 4, 7, 8, **Figure 5a, b**).

The reintroduction of extinct groups pushed the nutrient quantity toward pre-stressor community levels. However, the recovery in nutrient quantity did not reach reference levels within 7 years after the disturbance. The nutrient quantity was within 25-30% of the reference by year 10 of the simulation (scenarios 3-8, **Figure 5**) and hence showed only partial repair. Biotic interactions within the community dictate dynamics that inhibited a rapid return to reference conditions.

Patterns often differed in the response between communities and ecosystem functions (**Figure 6**). We for example found that some pairs of scenarios that had the same input effect strength were identical with respect to the number and identity of extinctions but showed quantitative and qualitative differences in their nutrient response pattern (scenarios 2, 3, **Figure 6b**). We also observed the opposite: some nutrient responses were indistinguishable between scenarios, even though these scenarios differed in the number of extinctions (scenarios 4, 11, **Figure 6d**).

**Figure 6.**
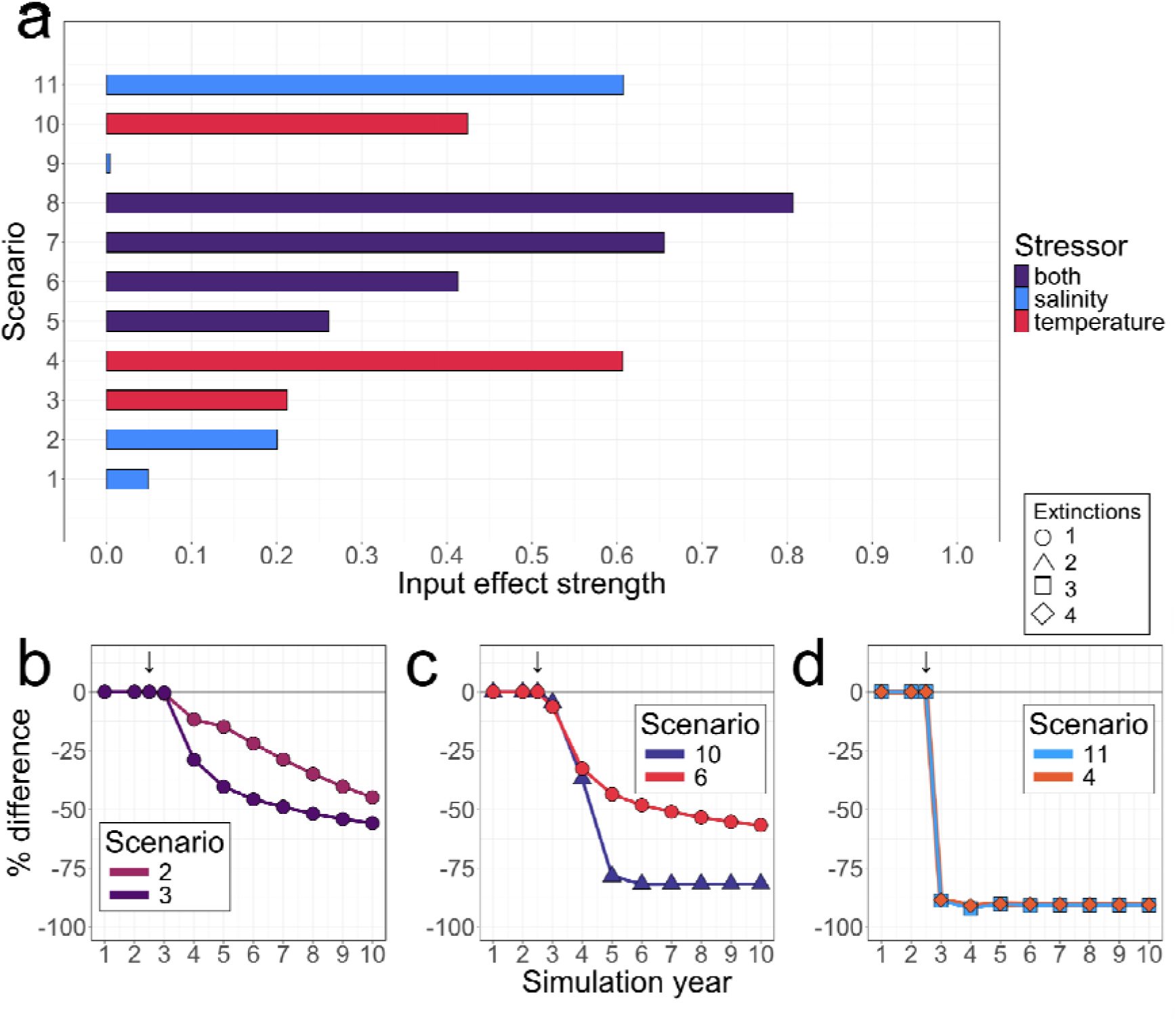
Comparison of the relative effects of all scenarios, with color indicating the stressors (**a**). Input effect strength consists of sum total of death rates for each scenario standardized to a scale between 0 and 1. Panels **b**, **c**, and **d** show the effects on nutrient in the system following the stressor phase, of pairs of scenarios with comparable input effect strength. The input effect strength of scenario pairs 2 and 3, 6 and 10, and 4 and 11 are the same (±0.01). Scenarios 2 and 3 have differing effects on the nutrient despite the same input effect strength and extinctions, but the nutrient approaches the same level over time (**b**). Scenarios 6 and 10 (**c**) have differing extinctions and effects on the nutrient. Scenarios 4 and 11 have indistinguishable effects on the nutrient, despite having different numbers of stressor-induced extinctions (**d**).

## 4. Discussion

The Asymmetric Response Concept provides a theoretical framework for the response of ecological communities to impacts and release from multiple stressors (Vos et al., 2023). The core of the ARC is a general and rather abstract idea: Variation in how the key mechanisms of tolerance, dispersal, and biotic interactions govern ecological responses over time determines which of the five common outcome types occurs in any particular scenario of stressor increase and release (see Figure 1 and Vos et al., 2023). Our analysis presents a practical test of the ARC by studying 11 scenarios of single or multiple stressor exposure and release using a stream food web model. Below, we discuss the results with respect to our hypotheses and in relation to the ARC.

### 4.1 Role of tolerance during the stressor phase in a food web context

First, we zoomed in on the mechanism of tolerance. We aggregated tolerance data from laboratory experiments with individual species into average tolerance values for each functional group. For example, in laboratory studies, fish species tend to be less tolerant of temperature increases than invertebrate predators (Appendix A). In the model, this is reflected in different increases in excess mortality for these functional groups in response to a high-temperature stressor phase. Our first hypothesis was that the strength of response to single or multiple stressors, measured by density decline during stressor exposure in the food web, should directly follow from the relative tolerance in the laboratory context. Given hypothesis 1, all animal groups should decline in the food web simulations in response to the stressors. In contrast, the detritivore functional group initially increased in density in *all* 11 stressor scenarios. This group benefited from the excess mortality experienced by other functional groups that produced additional animal detritus. This mechanism was particularly strong and caused a clear change in tolerance ranking in single and multiple stressor scenarios 4, 7 and 8 (**Table 1**), all of which included high (25°C) temperature stress. These stressor scenarios led to local extinction of the shredder, grazer, invertebrate predator and fish. The detritivore also experienced excess mortality, but this was compensated by a higher resource availability. Therefore, the detritivore moved rank to the least vulnerable position when stressed in a food web context (**Figure 3**, **Figure 4**). In our food web model, the detritivore did not compete with any other group for detritus and as it was no longer preyed upon due to the extinction of its two predators, its density continued to increase. In the absence of reintroductions, this led to a detritivore-dominated new state of the food web (**Figure 3f, Appendix C, Figure C2, s4, s7, s8**), whereas with reintroductions, the food web fully recovered after several years (**Figure 3i, Appendix C, Figure C3, s4, s7, s8**).

Furthermore, in 5 out of 11 scenarios (2, 3, 5, 6 and 10 **Table 1**, **Figure 3e, Appendix C, Figure C2**), the invertebrate predator increased in density. The underlying mechanism is that its fish predator decreased, which was caused by a substantial density decline in the grazer and shredder functional groups in response to the stressor. Large apex predators often suppress smaller mesopredators in a wide range of food webs (Strong and Frank, 2010). When the apex predator, in our case the fish, declined, this released predation pressure on the invertebrate predator to such a degree that its density increased, even though it was exposed to the same stressor as the other groups. That such interactions between species play a major role in community response to increased temperature was also shown by Zhang et al. (2017). In our study, stressor release led to a partial recovery of the system (**Figure 3e**). To test whether this outcome was really one of partial recovery rather than a very slowly recovering broken leg, we performed additional simulations (not shown). In these we swapped the densities of fish and the invertebrate predator at the end of the 10-year period (**Figure 3e**) and ran the simulation without stress for another 10 years. We found that the densities of both predators returned to the original state (i.e. as in **Figure 3e**). We also let the simulation run for 50 years without stress and found no further recovery towards the undisturbed baseline. Both tests show that the outcome of partial recovery following the extinction of the shredder is an alternative equilibrium, in which the relative densities of fish and the invertebrate predator differ from those in the undisturbed food web.

Many studies in real-world ecosystems confirm the general result that the food web influences the tolerance of species to stressors and the specific result that a stressor-driven release from predation and increase of available resources may compensate for stressor effects (Fleeger, 2020; Fleeger et al., 2003). For example, a comparison of species tolerances to chemical stressors between laboratory and mesocosm contexts found mismatches in both directions, i.e., higher and lower tolerance (Giddings et al., 2001). Similarly, many of the laboratory-derived metrics for thermal tolerance were correlated with the temperature range at which invertebrate taxa occur in real-world ecosystems, pointing to mechanisms of community assembly other than thermal tolerance (Sundermann et al., 2022). A constant increase in salinity in a mesocosm study led to changes in the food web that included an increase in functional groups in response to release from predation (Van Meter et al., 2011). Similarly, higher resource availability increased zooplankton taxa densities in microcosms, compensating for an initial decrease in response to chemical and solar radiation treatments, which was associated with changes in temperature, pH, salinity, and dissolved oxygen (Stampfli et al., 2011). Strong evidence that food webs change the responses of species to stressors has ignited a debate on the relevance of tolerance and biotic interactions in structuring communities (Cadotte and Tucker, 2017). Based on the ARC we hypothesized (Hypothesis 1), that during the stressor phase the relative tolerances in the laboratory and food web context should match. However, in all 11 tested scenarios, the rankings already differed during the stressor phase, suggesting that biotic interactions also play a major role in structuring communities during the stressor phase. Our results may not be directly transferable to real-world ecosystems because our food web model was comparatively simple and consisted of functional groups that were assigned a group-specific tolerance, whereas in real-world ecosystems functional groups can consist of many species with different tolerances (see Appendix A). In addition, our model relies on continuous growth of population biomass that does not account for differences between the growth of population size (i.e., number of individuals) and of individuals as part of their development, whereas in real world ecosystems, developmental cycles, for example, with resting egg stages and near-complete loss of adult biomass in winter, can lead to pronounced biomass cycles (Brüchner-Hüttemann et al., 2020). Thus, the extent to which biological interactions play a role will also depend on the relationship between stressor duration and the generation time of organisms, which can vary strongly from days to years between freshwater insect species (Huryn and Wallace, 2000). While we reject hypothesis 1 for our results, this may not generally be the case and suggests that the ARC needs to relate the stressor duration to the life histories of the organisms under scrutiny (Jackson et al., 2021).

### 4.2 Reintroduction and recovery of structure and function

Our second hypothesis focused on the effect of functional group reintroductions on nutrient cycling in food webs after the stressor phase. We hypothesized that successful reintroductions would lead to a rapid, nearly symmetric recovery of nutrient cycling that would, in contrast, be asymmetric in the absence of re-colonization. We did not observe such a fully symmetric pattern due to re- colonization in any of the studied stressor scenarios that resulted in extinctions. All attempted reintroductions were successful, but they never led to a full and very rapid ‘rubber band’ recovery of nutrient cycling (Outcome 1 in Fig. 1). Communities with reintroductions always require several years for recovery. Hence, our results suggest that hypothesis 2 should be rejected. As discussed above, our model comprised generalized functional groups with a uniform response, whereas functional groups in real communities are composed of many species and genotypes. If competition within functional groups plays an important role in the process of re-colonization of real stream food webs, this could impose stronger limits on recovery than those considered in our model. However, freshwater communities are classically thought to be less saturated than terrestrial communities, and a meta-analysis suggested that biotic resistance in freshwater ecosystems is driven by consumption rather than competition (Alofs and Jackson, 2014; Darwin, 1859). Furthermore, selection and evolutionary adaptive mechanisms certainly occur at the time scales simulated in this study, especially for groups with rapid life cycles, and are an important mechanism to consider for ongoing multiple stressor research (Hiltunen et al., 2018; Orr et al., 2022). Integrating mechanisms of evolutionary adaptation into food web models such as the one used in this study would offer a powerful tool to study shifts in community responses to multiple stressors over time. Local adaptation could create priority effects by monopolizing the local environment,and in turn, change the outcome of species reintroductions (De Meester et al., 2002; Meester et al., 2016; Tielke et al., 2020). Overall, a more complex community composition and community adaptation that occurs in real-world ecosystems may moderate our findings, but the asymmetric response pattern is likely robust. In a comparative review of recovery processes in rivers, lakes, estuarine, and coastal waters, Verdonschot et al. (2013) concluded that for recovery of the original biotic composition, diversity, and functioning, periods of 15 to 25 years may be needed in all four types of aquatic ecosystems. In some cases, recovery of rivers was even in the range of 30-40 years. The recovery periods for ‘broken leg’ scenarios in this model study were in the 5-10 years range. Such a relatively rapid recovery may have been driven by reintroductions very soon after the stressor release in our simulations. When there is dispersal limitation, for example because of physical barriers or because certain species have become very rare in the regional species pool, this can cause major time lags (Vos et al., 2023).

### 4.4 Recovery of structure vs. function exemplified by nutrient cycling

Reductions in the availability of free nutrient were only relatively small in stressor scenarios without the extinction of functional groups. Unless the lost group was reintroduced, the loss of a functional group resulted in a steady reduction in the available free nutrient (e.g., **Figure 5**). Thereby, the nutrient level reached a new state whereas the food web could partially recover. High temperature (25°C) led to a strong reduction without recovery as long as the lost functional groups remained absent. Interestingly, the pattern was a ‘new state’ outcome at the food web level (**Figure 3f**), but practically a ‘no recovery’ outcome at the ecosystem function level, in scenarios 4, 7 and 8 (without reintroduction) (**Figure 5**). In this case, the food web was temporarily locked in a degraded new state. The reintroduction of the lost functional groups caused a transition to a partial recovery outcome: recovery clearly occurred but was still incomplete at the end of the simulated 10-year period (**Figure 5**). Given that a gradual recovery was still visible, the outcome may follow the broken leg pattern at longer timescales. These results confirm the idea that the inclusion of both bottom-up control by nutrients and top-down control by consumer-resource interactions in food web models is essential if one aims to gain an understanding of ecosystem responses (DeAngelis, 1992). This is because consumers have strong effects on nutrients and detritus, which feeds back on consumers in functional groups throughout the food web (Vanni and de Ruiter, 1996). In real food webs, the recovery of structure and function will also be affected by biological stoichiometry (Elser et al., 2000). A meta-analysis of empirical studies showed that ecosystem functions recover more quickly than the original composition of communities and food webs (Hillebrand and Kunze, 2020). This pattern may be explained by the fact that only some species contribute most to any particular ecosystem function. For example, only 2–5 species contribute more than 90% of all carbon flow in a re-assembling macroinvertebrate community (Rossi et al., 2009). Processes of community adaptation might further lead to faster recovery of functions, as demonstrated in microcosm studies on leaf decomposition in different biogeographical regions (Schreiner et al., 2018). However, the recovery in our study lasted longer than the stressor degradation phase in 11 out of 11 cases, i.e., it was largely asymmetric.

### 4.5 Implications for the restoration of stream ecosystems

The food web model allowed us to simulate all five major outcomes defining community or ecosystem response to stressor exposure and release (Figure 1, Fig. 3 in Vos et al., 2023). The model can also be used to find stressor thresholds (such as for heat stress) for transitions between any of these outcomes. It can similarly be used to study which measures could speed up *the recovery rate* within the broken leg category or to increase the *degree of repair* in cases where recovery remains partial. In addition, alternative reintroduction scenarios can be tested for their restorative effect, especially in attempts to remedy degraded new states and no recovery cases. This model is a heuristic tool that provides general insight rather than simulate any particular stream in the field. However, our simulations show that the modeling approach we developed can be used to study the relationship between the intensity of single and multiple stressors, alternative restorative interventions, and the five possible outcomes addressed by the ARC. This study addresses a significantly higher level of food web complexity than the food web modules analyzed in Tielke et al. (2020) and Tielke and Vos (2024). The simultaneous reintroductions that were most successful in Tielke et al. (2020) and Tielke and Vos (2024) were also successful at this higher level of food web complexity (also see Dumeier et al., 2020).

## 5. Conclusions

One major conclusion is that biotic interactions can be more important during the stressor phase than is currently acknowledged by the ARC (Vos et al., 2023). Within the context of the food web model we studied, consumer–resource interactions were, with tolerance, already important *before* stressor release. Another major conclusion is that the observed response pattern was *asymmetric* in 10 of the 11 studied scenarios. Even in scenarios without loss of functional groups, densities tended to recover over several years. In these scenarios, internal recovery from the remaining organisms was sufficient for recovery to the initial density.

In contrast, reintroduction was indispensable for recovery when one or more functional groups had been lost in response to heat. This is not a trivial finding, as loss of the shredder could have been compensated for by increases in the bacterial and fungal functional groups. In real ecosystems, the loss of entire functional groups will only occur in response to extreme events. Recent decades have seen the emergence of a new category of extremely hot summers that were much less frequent before the 1980s. These are likely to occur more widely and frequently in the future (Hansen et al., 2012). In conclusion, our results suggest that the recovery of stream food webs will often be slow. Unassisted recovery can take decades. Our previous work (Tielke et al., 2020; Tielke and Vos, 2024) and our present results suggest that recovery will occur as long as all functional groups are present or are simultaneously reintroduced.

## Acknowledgments

Funded by the Deutsche Forschungsgemeinschaft (DFG, German Research Foundation) – SFB 1439/1 2021 – 426547801. We thank Armin Lorenz for his support with the field data. We are grateful to two anonymous expert Reviewers for their helpful and insightful comments and to Paola Verlicchi and Christoph Matthaei for all their editorial work.

## CRediT author statement

**Annabel Kuppels**: Conceptualization, Methodology, Software, Formal analysis, Visualization, Writing- Original Draft; **Helena Bayat**: Conceptualization, Methodology, Formal analysis, Visualization, Writing - Original Draft; **Svenja Gillmann**: Data Curation; **Ralf B. Schäfer**: Conceptualization, Writing - Original Draft, Writing - Review & Editing, Supervision; **Matthijs Vos**: Conceptualization, Methodology, Writing - Original Draft, Writing - Review & Editing, Supervision

## References

1. Alofs, K.M., Jackson, D.A., 2014. Meta-analysis suggests biotic resistance in freshwater environments is driven by consumption rather than competition. Ecology 95, 3259–3270. 10.1890/14-0060.1

2. Ashauer, R., O’Connor, I., Escher, B.I., 2017. Toxic Mixtures in Time—The Sequence Makes the Poison. Environ. Sci. Technol. 51, 3084–3092. 10.1021/acs.est.6b06163

3. Brose, U., Williams, R.J., Martinez, N.D., 2003. Comment on “Foraging Adaptation and the Relationship Between Food-Web Complexity and Stability.” Science 301, 918–918. 10.1126/science.1085902

4. Brüchner-Hüttemann, H., Ptatscheck, C., Traunspurger, W., 2020. Meiofauna in stream habitats: temporal dynamics of abundance, biomass and secondary production in different substrate microhabitats in a first-order stream. Aquat. Ecol. 54, 1079–1095. 10.1007/s10452-020-09795-5

5. Cadotte, M.W., Tucker, C.M., 2017. Should Environmental Filtering be Abandoned? Trends Ecol. Evol. 32, 429–437. 10.1016/j.tree.2017.03.004

6. Calvin, K., Dasgupta, D., Krinner, G., Mukherji, A., Thorne, P.W., Trisos, C., Romero, J., Aldunce, P., Barrett, K., Blanco, G., Cheung, W.W.L., Connors, S., Denton, F., Diongue-Niang, A., Dodman, D., Garschagen, M., Geden, O., Hayward, B., Jones, C., Jotzo, F., Krug, T., Lasco, R., Lee, Y.-Y., Masson-Delmotte, V., Meinshausen, M., Mintenbeck, K., Mokssit, A., Otto, F.E.L., Pathak, M., Pirani, A., Poloczanska, E., Pörtner, H.-O., Revi, A., Roberts, D.C., Roy, J., Ruane, A.C., Skea, J., Shukla, P.R., Slade, R., Slangen, A., Sokona, Y., Sörensson, A.A., Tignor, M., Van Vuuren, D., Wei, Y.-M., Winkler, H., Zhai, P., Zommers, Z., Hourcade, J.-C., Johnson, F.X., Pachauri, S., Simpson, N.P., Singh, C., Thomas, A., Totin, E., Arias, P., Bustamante, M., Elgizouli, I., Flato, G., Howden, M., Méndez-Vallejo, C., Pereira, J.J., Pichs-Madruga, R., Rose, S.K., Saheb, Y., Sánchez Rodríguez, R., Ürge-Vorsatz, D., Xiao, C., Yassaa, N., Alegría, A., Armour, K., Bednar- Friedl, B., Blok, K., Cissé, G., Dentener, F., Eriksen, S., Fischer, E., Garner, G., Guivarch, C., Haasnoot, M., Hansen, G., Hauser, M., Hawkins, E., Hermans, T., Kopp, R., Leprince-Ringuet, N., Lewis, J., Ley, D., Ludden, C., Niamir, L., Nicholls, Z., Some, S., Szopa, S., Trewin, B., Van Der Wijst, K.-I., Winter, G., Witting, M., Birt, A., Ha, M., Romero, J., Kim, J., Haites, E.F., Jung, Y., Stavins, R., Birt, A., Ha, M., Orendain, D.J.A., Ignon, L., Park, S., Park, Y., Reisinger, A., Cammaramo, D., Fischlin, A., Fuglestvedt, J.S., Hansen, G., Ludden, C., Masson-Delmotte, V., Matthews, J.B.R., Mintenbeck, K., Pirani, A., Poloczanska, E., Leprince-Ringuet, N., Péan, C., 2023. IPCC, 2023: Climate Change 2023: Synthesis Report. Contribution of Working Groups I, II and III to the Sixth Assessment Report of the Intergovernmental Panel on Climate Change [Core Writing Team, H. Lee and J. Romero (eds.)]. IPCC, Geneva, Switzerland. Intergovernmental Panel on Climate Change (IPCC). 10.59327/IPCC/AR6-9789291691647

7. Cañedo-Argüelles, M., Kefford, B.J., Piscart, C., Prat, N., Schäfer, R.B., Schulz, C.-J., 2013. Salinisation of rivers: An urgent ecological issue. Environ. Pollut. 173, 157–167. 10.1016/j.envpol.2012.10.011

8. Cordone, G., Salinas, V., Marina, T.I., Doyle, S.R., Pasotti, F., Saravia, L.A., Momo, F.R., 2020. Green vs brown food web: Effects of habitat type on multidimensional stability proxies for a highly- resolved Antarctic food web. Food Webs 25, e00166. 10.1016/j.fooweb.2020.e00166

9. Darwin, C.R., 1859. On the origin of species by means of natural selection, or the preservation of favoured races in the struggle for life., 1st ed. John Murray, London, UK.

10. De Laender, F., 2018. Community- and ecosystem-level effects of multiple environmental change drivers: Beyond null model testing. Glob. Change Biol. 24, 5021–5030. 10.1111/gcb.14382

11. De Meester, L., Gómez, A., Okamura, B., Schwenk, K., 2002. The Monopolization Hypothesis and the dispersal–gene flow paradox in aquatic organisms. Acta Oecologica 23, 121–135. 10.1016/S1146-609X(02)01145-1

12. de Ruiter, P.C., Neutel, A.-M., Moore, J.C., 1995. Energetics, Patterns of Interaction Strengths, and Stability in Real Ecosystems. Science 269, 1257–1260. 10.1126/science.269.5228.1257

13. DeAngelis, D.L., 1992. Dynamics of Nutrient Cycling and Food Webs. Springer Netherlands, Dordrecht. 10.1007/978-94-011-2342-6

14. Downs, P.W., Skinner, K.S., Kondolf, G.M., 2002. 12. Rivers and streams, in: Handbook of Ecological Restoration. Volume 2 Restoration in Practice. Cambridge University Press, pp. 267–296.

15. Dumeier, A.C., Lorenz, A.W., Kiel, E., 2020. Active reintroduction of benthic invertebrates to increase stream biodiversity. Limnologica 80, 125726. 10.1016/j.limno.2019.125726

16. Elser, J.J., Fagan, W.F., Denno, R.F., Dobberfuhl, D.R., Folarin, A., Huberty, A., Interlandi, S., Kilham, S.S., McCauley, E., Schulz, K.L., Siemann, E.H., Sterner, R.W., 2000. Nutritional constraints in terrestrial and freshwater food webs. Nature 408, 578–580. 10.1038/35046058

17. Evans-White, M.A., Halvorson, H.M., 2017. Comparing the Ecological Stoichiometry in Green and Brown Food Webs – A Review and Meta-analysis of Freshwater Food Webs. Front. Microbiol. 8.

18. Fleeger, J.W., 2020. How Do Indirect Effects of Contaminants Inform Ecotoxicology? A Review. Processes 8, 1659. 10.3390/pr8121659

19. Fleeger, J.W., Carman, K.R., Nisbet, R.M., 2003. Indirect effects of contaminants in aquatic ecosystems. Sci. Total Environ. 317, 207–233. 10.1016/S0048-9697(03)00141-4

20. Garnier, S., Ross, N., BoB Rudis, Filipovic-Pierucci, A., Galili, T., Timelyportfolio, O’Callaghan, A., Greenwell, B., Sievert, C., Harris, D.J., Sciaini, M., JJ Chen, 2023. viridis(Lite) - Colorblind-Friendly Color Maps for R. 10.5281/ZENODO.4679423

21. Giddings, J.M., Solomon, K.R., Maund, S.J., 2001. Probabilistic risk assessment of cotton pyrethroids: II. Aquatic mesocosm and field studies. Environ. Toxicol. Chem. 20, 660–668. 10.1002/etc.5620200327

22. Gillmann, S.M., Hering, D., Lorenz, A.W., 2023. Habitat development and species arrival drive succession of the benthic invertebrate community in restored urban streams. Environ. Sci. Eur. 35, 49. 10.1186/s12302-023-00756-x

23. Gunderson, A.R., Armstrong, E.J., Stillman, J.H., 2016. Multiple Stressors in a Changing World: The Need for an Improved Perspective on Physiological Responses to the Dynamic Marine Environment. Annu. Rev. Mar. Sci. 8, 357–378. 10.1146/annurev-marine-122414-033953

24. Hansen, J., Sato, M., Ruedy, R., 2012. Perception of climate change. Proc. Natl. Acad. Sci. 109, E2415–E2423. 10.1073/pnas.1205276109

25. Hillebrand, H., Kunze, C., 2020. Meta-analysis on pulse disturbances reveals differences in functional and compositional recovery across ecosystems. 10.1111/ele.13457

26. Hiltunen, T., Cairns, J., Frickel, J., Jalasvuori, M., Laakso, J., Kaitala, V., Künzel, S., Karakoc, E., Becks, L., 2018. Dual-stressor selection alters eco-evolutionary dynamics in experimental communities. Nat. Ecol. Evol. 2, 1974–1981. 10.1038/s41559-018-0701-5

27. Huang, W., Liu, X., Peng, W., Wu, L., Yano, S., Zhang, J., Zhao, F., 2018. Periphyton and ecosystem metabolism as indicators of river ecosystem response to environmental flow restoration in a flow-reduced river. Ecol. Indic., Multi-Scale Ecological Indicators for Supporting Sustainable Watershed Management 92, 394–401. 10.1016/j.ecolind.2017.11.025

28. Huryn, A.D., Wallace, J.B., 2000. Life History and Production of Stream Insects. Annu. Rev. Entomol. 45, 83–110. 10.1146/annurev.ento.45.1.83

29. Jackson, M.C., Loewen, C.J.G., Vinebrooke, R.D., Chimimba, C.T., 2016. Net effects of multiple stressors in freshwater ecosystems: a meta-analysis. Glob. Change Biol. 22, 180–189. 10.1111/GCB.13028

30. Jackson, M.C., Pawar, S., Woodward, G., 2021. The Temporal Dynamics of Multiple Stressor Effects: From Individuals to Ecosystems. Trends Ecol. Evol. 36, 402–410. 10.1016/j.tree.2021.01.005

31. Jacobs, G., 2017. Bericht zur Fischfauna der Fließgewässer im Einzugsgebiet der Emscher. Jourdan, J., Plath, M., Tonkin, J.D., Ceylan, M., Dumeier, A.C., Gellert, G., Graf, W., Hawkins, C.P., Kiel, E., Lorenz, A.W., Matthaei, C.D., Verdonschot, P.F.M., Verdonschot, R.C.M., Haase, P., 2019. Reintroduction of freshwater macroinvertebrates: challenges and opportunities. Biol. Rev. 94, 368–387. 10.1111/brv.12458

32. Kalinkat, G., Rall, B.C., Uiterwaal, S.F., Uszko, W., 2023. Empirical evidence of type III functional responses and why it remains rare. Front. Ecol. Evol. 11.

33. Kalinkat, G., Schneider, F.D., Digel, C., Guill, C., Rall, B.C., Brose, U., 2013. Body masses, functional responses and predator–prey stability. Ecol. Lett. 16, 1126–1134. 10.1111/ele.12147

34. Kaushal, S.S., Groffman, P.M., Likens, G.E., Belt, K.T., Stack, W.P., Kelly, V.R., Band, L.E., Fisher, G.T., 2005. Increased salinization of fresh water in the northeastern United States. Proc. Natl. Acad. Sci. 102, 13517–13520. 10.1073/pnas.0506414102

35. Kaushal, S.S., Likens, G.E., Pace, M.L., Reimer, J.E., Maas, C.M., Galella, J.G., Utz, R.M., Duan, S., Kryger, J.R., Yaculak, A.M., Boger, W.L., Bailey, N.W., Haq, S., Wood, K.L., Wessel, B.M., Park, C.E., Collison, D.C., Aisin, B.Y. ’aaqob I., Gedeon, T.M., Chaudhary, S.K., Widmer, J., Blackwood, C.R., Bolster, C.M., Devilbiss, M.L., Garrison, D.L., Halevi, S., Kese, G.Q., Quach, E.K., Rogelio, C.M.P., Tan, M.L., Wald, H.J.S., Woglo, S.A., 2021. Freshwater salinization syndrome: from emerging global problem to managing risks. Biogeochemistry 154, 255–292. 10.1007/s10533-021-00784-w

36. Lake, P.S., Bond, N., Reich, P., 2007. Linking ecological theory with stream restoration. Freshw. Biol. 52, 597–615. 10.1111/j.1365-2427.2006.01709.x

37. Leach, J.A., Hudson, D.T., Moore, R.D., 2022. Assessing stream temperature response and recovery for different harvesting systems in northern hardwood forests using 40 years of spot measurements. Hydrol. Process. 36, e14753. 10.1002/hyp.14753

38. Lundberg, P., Ranta, E., Kaitala, V., 2000. Species loss leads to community closure. Ecol. Lett. 3, 465– 468. 10.1111/j.1461-0248.2000.00170.x

39. Lynch, A.J., Cooke, S.J., Arthington, A.H., Baigun, C., Bossenbroek, L., Dickens, C., Harrison, I., Kimirei, I., Langhans, S.D., Murchie, K.J., Olden, J.D., Ormerod, S.J., Owuor, M., Raghavan, R., Samways, M.J., Schinegger, R., Sharma, S., Tachamo-Shah, R.-D., Tickner, D., Tweddle, D., Young, N., Jähnig, S.C., 2023. People need freshwater biodiversity. WIREs Water 10, e1633. 10.1002/wat2.1633

40. MacLennan, M.M., Vinebrooke, R.D., 2021. Exposure order effects of consecutive stressors on communities: the role of co-tolerance. Oikos 130, 2111–2121. 10.1111/oik.08884

41. Majumdar, P., Debnath, S., Sarkar, S., Ghosh, U., 2022. The Complex Dynamical Behavior of a Prey- Predator Model with Holling Type-III Functional Response and Non-Linear Predator Harvesting. Int. J. Model. Simul. 42, 287–304. 10.1080/02286203.2021.1882148

42. Masson-Delmotte, V., Zhai, P., Pirani, A., Connors, S.L., Péan, C., Berger, S., Caud, N., Chen, Y., Goldfarb, L., Gomis, M.I., Huang, M., Leitzell, K., Lonnoy, E., Matthews, J.B.R., Maycock, T.K., Waterfield, T., Yelekçi, Ö., Yu, R., Zhou, B. (Eds.), 2021. Climate Change 2021: The Physical Science Basis. Contribution of Working Group I to the Sixth Assessment Report of the Intergovernmental Panel on Climate Change. Cambridge University Press, Cambridge, United Kingdom and New York, NY, USA. 10.1017/9781009157896

43. Masson-Delmotte, V.P., Zhai, P., Pirani, S.L., Connors, C., Péan, S., Berger, N., Caud, Y., Chen, L., Goldfarb, M.I., Scheel Monteiro, P.M., 2021. IPCC, 2021: Summary for Policymakers. In: Climate Change 2021: The Physical Science Basis. Contribution of Working Group I to the Sixth Assessment Report of the Intergovernmental Panel on Climate Change (Report). Cambridge University Press, Cambridge, United Kingdom and New York, NY, USA.

44. May, R.M., 1972. Will a Large Complex System be Stable? Nature 238, 413–414. 10.1038/238413a0

45. McCann, K., Hastings, A., Huxel, G.R., 1998. Weak trophic interactions and the balance of nature. Nature 395, 794–798. 10.1038/27427

46. Meester, L.D., Vanoverbeke, J., Kilsdonk, L.J., Urban, M.C., 2016. Evolving Perspectives on Monopolization and Priority Effects. Trends Ecol. Evol. 31, 136–146. 10.1016/j.tree.2015.12.009

47. Meyling, N.V., Arthur, S., Pedersen, K.E., Dhakal, S., Cedergreen, N., Fredensborg, B.L., 2018. Implications of sequence and timing of exposure for synergy between the pyrethroid insecticide alpha-cypermethrin and the entomopathogenic fungus Beauveria bassiana. Pest Manag. Sci. 74, 2488–2495. 10.1002/ps.4926

48. Moog, O., 1995. Fauna Aquatica Austriaca. Bundesministerium für Land- und Forstwirtschaft, Umwelt und Wasserwirtschaft, Wien.

49. Moore, J.C., Berlow, E.L., Coleman, D.C., de Ruiter, P.C., Dong, Q., Hastings, A., Johnson, N.C., McCann, K.S., Melville, K., Morin, P.J., Nadelhoffer, K., Rosemond, A.D., Post, D.M., Sabo, J.L., Scow, K.M., Vanni, M.J., Wall, D.H., 2004. Detritus, trophic dynamics and biodiversity. Ecol. Lett. 7, 584–600. 10.1111/j.1461-0248.2004.00606.x

50. Mougi, A., 2020. Coupling of green and brown food webs and ecosystem stability. Ecol. Evol. 10, 9192–9199. 10.1002/ece3.6586

51. Muhar, S., Januschke, K., Kail, J., Poppe, M., Schmutz, S., Hering, D., Buijse, A.D., 2016. Evaluating good-practice cases for river restoration across Europe: context, methodological framework, selected results and recommendations. Hydrobiologia 769, 3–19. 10.1007/s10750-016-2652-7

52. Neutel, A.-M., Heesterbeek, J.A.P., de Ruiter, P.C., 2002. Stability in Real Food Webs: Weak Links in Long Loops. Science 296, 1120–1123. 10.1126/science.1068326

53. Neutel, A.-M., Heesterbeek, J.A.P., van de Koppel, J., Hoenderboom, G., Vos, A., Kaldeway, C., Berendse, F., de Ruiter, P.C., 2007. Reconciling complexity with stability in naturally assembling food webs. Nature 449, 599–602. 10.1038/nature06154

54. O’Brien, P.J., Wehr, J.D., 2010. Periphyton biomass and ecological stoichiometry in streams within an urban to rural land-use gradient, in: Stevenson, R.J., Sabater, S. (Eds.), Global Change and River Ecosystems—Implications for Structure, Function and EcosystemServices, Developments in Hydrobiology 215. Springer Netherlands, Dordrecht, pp. 89–105. 10.1007/978-94-007-0608-8_7

55. Orr, J.A., Luijckx, P., Arnoldi, J.-F., Jackson, A.L., Piggott, J.J., 2022. Rapid evolution generates synergism between multiple stressors: Linking theory and an evolution experiment. Glob. Change Biol. 28, 1740–1752. 10.1111/gcb.15633

56. Orr, J.A., Vinebrooke, R.D., Jackson, M.C., Kroeker, K.J., Kordas, R.L., Mantyka-Pringle, C., van den Brink, P.J., de Laender, F., Stoks, R., Holmstrup, M., Matthaei, C.D., Monk, W.A., Penk, M.R., Leuzinger, S., Schäfer, R.B., Piggott, J.J., 2020. Towards a unified study of multiple stressors: Divisions and common goals across research disciplines. Proc. R. Soc. B Biol. Sci. 287. 10.1098/rspb.2020.0421

57. Pedersen, TL., 2023. patchwork: The Composer of Plots.

58. Piggott, J.J., Townsend, C.R., Matthaei, C.D., 2015. Reconceptualizing synergism and antagonism among multiple stressors. Ecol. Evol. 5, 1538–1547. 10.1002/ece3.1465

59. R Core Team, 2022. R: A language and environment for statistical computing.

60. Reid, A.J., Carlson, A.K., Creed, I.F., Eliason, E.J., Gell, P.A., Johnson, P.T.J., Kidd, K.A., MacCormack, T.J., Olden, J.D., Ormerod, S.J., Smol, J.P., Taylor, W.W., Tockner, K., Vermaire, J.C., Dudgeon, D., Cooke, S.J., 2019. Emerging threats and persistent conservation challenges for freshwater biodiversity. Biol. Rev. 94, 849–873. 10.1111/brv.12480

61. Rillig, M.C., Ryo, M., Lehmann, A., Aguilar-Trigueros, C.A., Buchert, S., Wulf, A., Iwasaki, A., Roy, J., Yang, G., 2019. The role of multiple global change factors in driving soil functions and microbial biodiversity. Science 366, 886–890. 10.1126/science.aay2832

62. Rossi, F., Vos, M., Middelburg, J.J., 2009. Species identity, diversity and microbial carbon flow in reassembling macrobenthic communities. Oikos 118, 503–512. 10.1111/j.1600-0706.2008.17112.x

63. Sarr, D.A., 2002. Riparian Livestock Exclosure Research in the Western United States: A Critique and Some Recommendations. Environ. Manage. 30, 516–526. 10.1007/s00267-002-2608-8

64. Schäfer, R.B., Jackson, M., Juvigny-Khenafou, N., Osakpolor, S.E., Posthuma, L., Schneeweiss, A., Spaak, J., Vinebrooke, R., 2023. Chemical Mixtures and Multiple Stressors: Same but Different? Environ. Toxicol. Chem. 42, 1915–1936. 10.1002/etc.5629

65. Scharmüller, A., Schreiner, V.C., Schäfer, R.B., 2020. Standartox: Standardizing Toxicity Data. Data 5. 10.5281/zenodo.3785031

66. Schmidt-Kloiber, A., Hering, D., 2015. www.freshwaterecology.info – An online tool that unifies, standardises and codifies more than 20,000 European freshwater organisms and their ecological preferences. Ecol. Indic. 53, 271–282. 10.1016/j.ecolind.2015.02.007

67. Schneeweiss, A., Juvigny-Khenafou, N.P.D., Osakpolor, S., Scharmüller, A., Scheu, S., Schreiner, V.C., Ashauer, R., Escher, B.I., Leese, F., Schäfer, R.B., 2023. Three perspectives on the prediction of chemical effects in ecosystems. Glob. Change Biol. 29, 21–40. 10.1111/gcb.16438

68. Schreiner, V.C., Feckler, A., Fernández, D., Frisch, K., Muñoz, K., Szöcs, E., Zubrod, J.P., Bundschuh, M., Rasmussen, J.J., Kefford, B.J., Axelsen, J., Cedergreen, N., Schäfer, R.B., 2018. Similar recovery time of microbial functions from fungicide stress across biogeographical regions. Sci. Rep. 8, 17021. 10.1038/s41598-018-35397-1

69. Simmons, B.I., Blyth, P.S.A., Blanchard, J.L., Clegg, T., Delmas, E., Garnier, A., Griffiths, C.A., Jacob, U., Pennekamp, F., Petchey, O.L., Poisot, T., Webb, T.J., Beckerman, A.P., 2021. Refocusing multiple stressor research around the targets and scales of ecological impacts. Nat. Ecol. Evol. 10.1038/s41559-021-01547-4

70. Speißer, B., Wilschut, R.A., van Kleunen, M., 2022. Number of simultaneously acting global change factors affects composition, diversity and productivity of grassland plant communities. Nat. Commun. 13, 7811. 10.1038/s41467-022-35473-1

71. Stampfli, N.C., Knillmann, S., Liess, M., Beketov, M.A., 2011. Environmental context determines community sensitivity of freshwater zooplankton to a pesticide. Aquat. Toxicol. 104, 116– 124. 10.1016/j.aquatox.2011.04.004

72. Straalen, N.M.V., 2003. Ecotoxicology Becomes Stress Ecology. Environ. Sci. Technol. 37, 324A–330A. 10.1017/CBO9781107415324.004

73. Strong, D.R., Frank, K.T., 2010. Human Involvement in Food Webs. Annu. Rev. Environ. Resour. 35, 1– 23. 10.1146/annurev-environ-031809-133103

74. Sundermann, A., Müller, A., Halle, M., 2022. A new index of a water temperature equivalent for summer respiration conditions of benthic invertebrates in rivers as a bio-indicator of global climate change. Limnologica 95. 10.1016/j.limno.2022.125980

75. Tielke, A.-K., Karreman, J., Vos, M., 2020. Mild cycles open closed communities to ecological restoration. Restor. Ecol. 28, 841–849. 10.1111/rec.13136

76. Tielke, A.-K., Vos, M., 2024. Successful reintroduction of species: improving on windows of opportunity for biodiversity repair. Restor. Ecol. 32, e14091. 10.1111/rec.14091

77. Tomczyk, N.J., Rosemond, A.D., Rogers, P.A., Cummins, C.S., 2022. Thermal traits of freshwater macroinvertebrates vary with feeding group and phylogeny. Freshw. Biol. 67, 1994–2003. 10.1111/fwb.13992

78. Van Meter, R.J., Swan, C.M., Leips, J., Snodgrass, J.W., 2011. Road Salt Stress Induces Novel Food Web Structure and Interactions. Wetlands 31, 843–851. 10.1007/s13157-011-0199-y

79. Vanni, M.J., de Ruiter, P.C., 1996. Detritus and Nutrients in Food Webs, in: Polis, G.A., Winemiller, K.O. (Eds.), Food Webs: Integration of Patterns & Dynamics. Springer US, Boston, MA, pp. 25–29. 10.1007/978-1-4615-7007-3_2

80. Venâncio, C., Wijewardene, L., Ribeiro, R., Lopes, I., 2023. Combined effects of two abiotic stressors (salinity and temperature) on a laboratory-simulated population of *Daphnia longispina*. Hydrobiologia 850, 3197–3208. 10.1007/s10750-023-05249-9

81. Verdonschot, P.F.M., Spears, B.M., Feld, C.K., Brucet, S., Keizer-Vlek, H., Borja, A., Elliott, M., Kernan, M., Johnson, R.K., 2013. A comparative review of recovery processes in rivers, lakes, estuarine and coastal waters. Hydrobiologia 704, 453–474. 10.1007/s10750-012-1294-7

82. Vinebrooke, R.D., Cottingham, K.L., Norberg, J., Marten Scheffer, Dodson, S.I., Maberly, S.C., Sommer, U., 2004. Impacts of multiple stressors on biodiversity and ecosystem functioning: the role of species co-tolerance. Oikos 104, 451–457. 10.1111/j.0030-1299.2004.13255.x

83. Vos, M., Hering, D., Gessner, M.O., Leese, F., Schäfer, R.B., Tollrian, R., Boenigk, J., Haase, P., Meckenstock, R., Baikova, D., Bayat, H., Beermann, A., Beisser, D., Beszteri, B., Birk, S., Boden, L., Brauer, V., Brauns, M., Buchner, D., Burfeid-Castellanos, A., David, G., Deep, A., Doliwa, A., Dunthorn, M., Enß, J., Escobar-Sierra, C., Feld, C.K., Fohrer, N., Grabner, D., Hadziomerovic, U., Jähnig, S.C., Jochmann, M., Khaliq, S., Kiesel, J., Kuppels, A., Lampert, K.P., Le, T.T.Y., Lorenz, A.W., Madariaga, G.M., Meyer, B., Pantel, J.H., Pimentel, I.M., Mayombo, N.S., Nguyen, H.H., Peters, K., Pfeifer, S.M., Prati, S., Probst, A.J., Reiner, D., Rolauffs, P., Schlenker, A., Schmidt, T.C., Shah, M., Sieber, G., Stach, T.L., Tielke, A.-K., Vermiert, A.-M., Weiss, M., Weitere, M., Sures, B., 2023. The Asymmetric Response Concept explains ecological consequences of multiple stressor exposure and release. Sci. Total Environ. 872, 162196. 10.1016/j.scitotenv.2023.162196

84. Wallace, J.B., Webster, J.R., 1996. The Role of Macroinvertebrates in Stream Ecosystem Function. Annu. Rev. Entomol. 41, 115–139. 10.1146/annurev.en.41.010196.000555

85. Welcomme, R.L., Ryder, R.A., Sedell, J.A., 1989. Dynamics of fish assemblages in river systems-a synthesis. Can Spec Publ Fish Aquat Sci 106, 569–577.

86. Wickham, H., 2016. ggplot2: Elegant Graphics for Data Analysis. Springer-Verlag, New York. Wickham, H., Averick, M., Bryan, J., Chang, W., McGowan, L., François, R., Grolemund, G., Hayes, A., Henry, L., Hester, J., Kuhn, M., Pedersen, T., Miller, E., Bache, S., Müller, K., Ooms, J., Robinson, D., Seidel, D., Spinu, V., Takahashi, K., Vaughan, D., Wilke, C., Woo, K., Yutani, H., 2019. Welcome to the Tidyverse. J. Open Source Softw. 4, 1686. 10.21105/joss.01686

87. Wootton, J.T., 1994. The nature and consequences of indirect effects in ecological communities. Annu. Rev. Ecol. Syst. 25, 443–466. 10.1146/annurev.es.25.110194.002303

88. Zhang, L., Takahashi, D., Hartvig, M., Andersen, K.H., 2017. Food-web dynamics under climate change. Proc. R. Soc. B Biol. Sci. 284, 20171772. 10.1098/rspb.2017.1772

89. Zelnik, Y.R., Manzoni, S., Bommarco, R., 2022. The coordination of green–brown food webs and their disruption by anthropogenic nutrient inputs. Glob. Ecol. Biogeogr. 31, 2270–2280. 10.1111/geb.13576

90. Zou, K., Thébault, E., Lacroix, G., Barot, S., 2016. Interactions between the green and brown food web determine ecosystem functioning. Funct. Ecol. 30, 1454–1465. 10.1111/1365-2435.12626

