## Supplementary material for "Putting the Asymmetric Response Concept to the test: modeling multiple stressor exposure and release in a stream food web": Supp Appendix A

1. Background
   1.1. Metrics of tolerance
        1.1.1. Thermal tolerance

Biological thermal tolerance has garnered scientific interest for at least the last 200 years, with reviews comparing thermal tolerances of different organisms published since the 1870s (Cameron & Brownlee, 1915; Davenport & Castle, 1895; J. Davy, personal communication, 1855; Hoppe-Seyler, 1875; Spallanzani, 1777; Vernon, 1899). Though different methodologies have developed to compare thermal tolerances, the critical thermal methodology has gained popularity due to its ease of implementation and applicability to the maxima and minima of the thermal performance curve (TPC) (Beitinger et al., 2000; Cowles & Bogert, 1944; Sinclair et al., 2016). TPCs are characterized by a steep decline between the optimum (peak of the curve) and the critical thermal maximum (CTmax) which delineates the upper limit of thermal tolerance (**Figure A1**). This curve shape characterized by a steep decline is observed across performance metrics and organism groups (Rezende & Bozinovic, 2019). The CTmax marks the point at which an organism becomes disorganized and loses equilibrium, and thus would swiftly die in a natural setting. Some studies quantify the lethal thermal maximum (LTmax), which entails ramping the temperature beyond the CTmax until the organism is completely dead (Chatterjee et al., 2004).

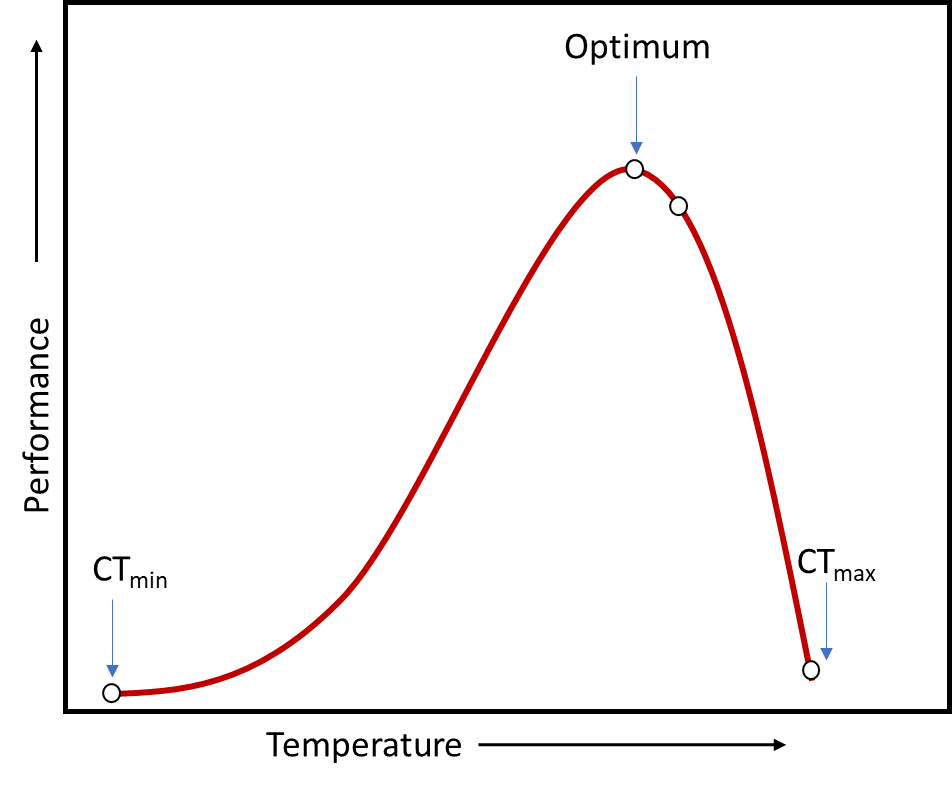

**Figure A1**. A generalized thermal performance curve (TPC), demonstrating the critical thermal minimum (CTmin) as the lower limit, the critical thermal maximum (CTmax) as the upper limit, and the optimum as the peak. The TPC is characteristically asymmetric in shape, with a steep decline from the optimum until the CTmax.

1.1.2. Salinity tolerance

Salinity tolerance in freshwater ecological research has been identified as a trait of organisms, in a fuzzy-coded system derived from occurrences under field conditions (Tachet et al., 2010). However, salt is also a chemical that has been tested in standard ecotoxicological concentration-response assays under the same methodology that pesticides (among other chemicals) are tested.

The standard concentration-response test consists of exposing groups of individual organisms to a series of increasing concentrations over a set time period, then monitoring an endpoint within each concentration group (OECD, 2002). The response of the chosen endpoint at different concentrations (for a defined time period) is analyzed using probit analysis (old-fashioned) or fitting a log-logistic function to obtain an effect concentration at which 50% of the population reaches the endpoint, commonly known as the EC50 (Ritz et al., 2015). For lethal endpoints, the lethal concentration at which 50% of the population dies is known as the LC50, and is one of the most commonly tested endpoints in ecotoxicological assays. The US Environmental Protection Agency has compiled a continually-updated database of ecotoxicological assays which includes results from government agencies, gray literature, and peer reviewed research studies for a variety of chemical classes, including salts (Olker et al., 2022).

1.2. Applications of individual tolerance data

Individual tolerance to stressors, most notably chemicals but also environmental factors such as desiccation, oxygen, and heat, can be measured in laboratory assays to compare sensitivity among organisms (Scharmüller et al., 2020; Suemoto et al., 2004; Walshe, 1948). These assays frequently run on timescales that fit conveniently into a working week for standard test organisms that are easily maintained in the laboratory (OECD, 2002). Public institutions, such as environmental protection agencies of various countries, run these tests according to strict international guidelines to set threshold values for stressors in drinking water (OECD, 2002; *Technical Guidance for Deriving Environmental Quality Standards*, 2018). While these tests and determinations of threshold values are primarily conducted for chemicals, thresholds may also be set for environmental factors such as temperature. For example, the advent of nuclear power after the 1950s spurred implementation of thresholds for thermal pollution of freshwater ecosystems resulting from cooling water outflows (Coutant & Brook, 1970; Coutant & Goodyear, 1972; Olsen et al., 2012; *Temperature: Water Quality Standards Criteria Summaries: A Compilation of State/Federal Criteria*, 1988). Due to the presence of standardized protocols and their use by government agencies, individual tolerance data from comparable laboratory assays is available for a multitude of freshwater species (Olker et al., 2022; Scharmüller et al., 2020). Crossover of methodology, such as applying acute toxicity assays, and ideas between the fields of ecotoxicology and ecology has increased within the burgeoning field of multiple stressor research (Orr et al., 2020; Schäfer et al., 2023; Straalen, 2003).

2. Methods.
   2.1. Individual tolerance in the laboratory context
        2.1.1. Thermal tolerance
               2.1.1.1. Data acquisition

To obtain upper thermal tolerance data for fish, known databases were searched for central European fish taxa (Bennett, 2018; Cereja, 2020; Leiva et al., 2019). When necessary, additional information was queried from source studies.

Freshwater invertebrates were not well represented in existing compilations of thermal tolerance. Thus, upper thermal tolerance for freshwater invertebrates was queried using a taxa list of freshwater invertebrates obtained from field sampling campaigns in the Boye catchment, implemented in combination with a search string in Google Scholar (**Table A1**) (Gillmann et al., 2023).

**Table A1**. Search strings implemented in Google Scholar to obtain studies on upper thermal tolerance for a taxa list of 475 central European freshwater invertebrates. All 475 species names were queried, as well as queries with just the genus and the families represented in the taxa list.

**Search string in Google Scholar**

"species name" AND "thermal tolerance" OR "heat tolerance" OR "ctmax" OR "incipient lethal temperature" OR "critical thermal maximum" OR "thermal limit"

"genus name" AND "thermal tolerance" OR "heat tolerance" OR "ctmax" OR "incipient lethal temperature" OR "critical thermal maximum" OR "thermal limit"

"family name" AND "thermal tolerance" OR "heat tolerance" OR "ctmax" OR "incipient lethal temperature" OR "critical thermal maximum" OR "thermal limit"

2.1.1.2. Aggregation of tolerance data

The data queried from existing databases and literature searches often contained multiple thermal tolerance measures for the same species or genus, under different test conditions or using a different test design.

To keep all data comparable, only data from temperate locations was included. When multiple tests were conducted, the result from the lowest acclimation temperature (usually 10 - 15°C) was selected. Tests measuring both LTmax and CTmax in the same organisms were compared to determine a relationship between these two metrics. All LTmax values were subsequently corrected to be comparable to the CTmax, using the following relationship, as depicted graphically in **Figure A2**:

$corrected thermal maximum = 0.6605 \times LTmax + 7.0561$	(1)

Relationships between thermal tolerance metrics determined from different methodologies have consistently been found in the literature and provide the basis for such a standardization (Dallas & Ketley, 2011; Jørgensen et al., 2021; Rezende et al., 2014, p. 201). After selecting the lowest acclimation temperature and correcting LTmax metrics, the thermal tolerance was averaged per species. When no species level data was available for relevant taxa, genus level data was used. Genus level data was averaged across species when multiple species of the same genus were included, but for many of the taxa only one species represented a genus within the dataset.

Invertebrate species derived from the aforementioned taxa list were classified into feeding groups according to Moog 1995, as obtained from the freshwater ecology database freshwaterecology.info (Moog, 1995; Schmidt-Kloiber & Hering, 2015). The feeding group classifications of Moog were modified slightly to fit our food web; all taxa classified as active filter feeders, passive filter feeders, and gatherers were combined into the detritivore functional group. The categories of grazer, shredder, and invertebrate predator were not modified. The thermal tolerance values of taxa within each category were averaged to obtain a thermal tolerance value for each functional group. For the fish group, all relevant fish thermal tolerances were averaged together.

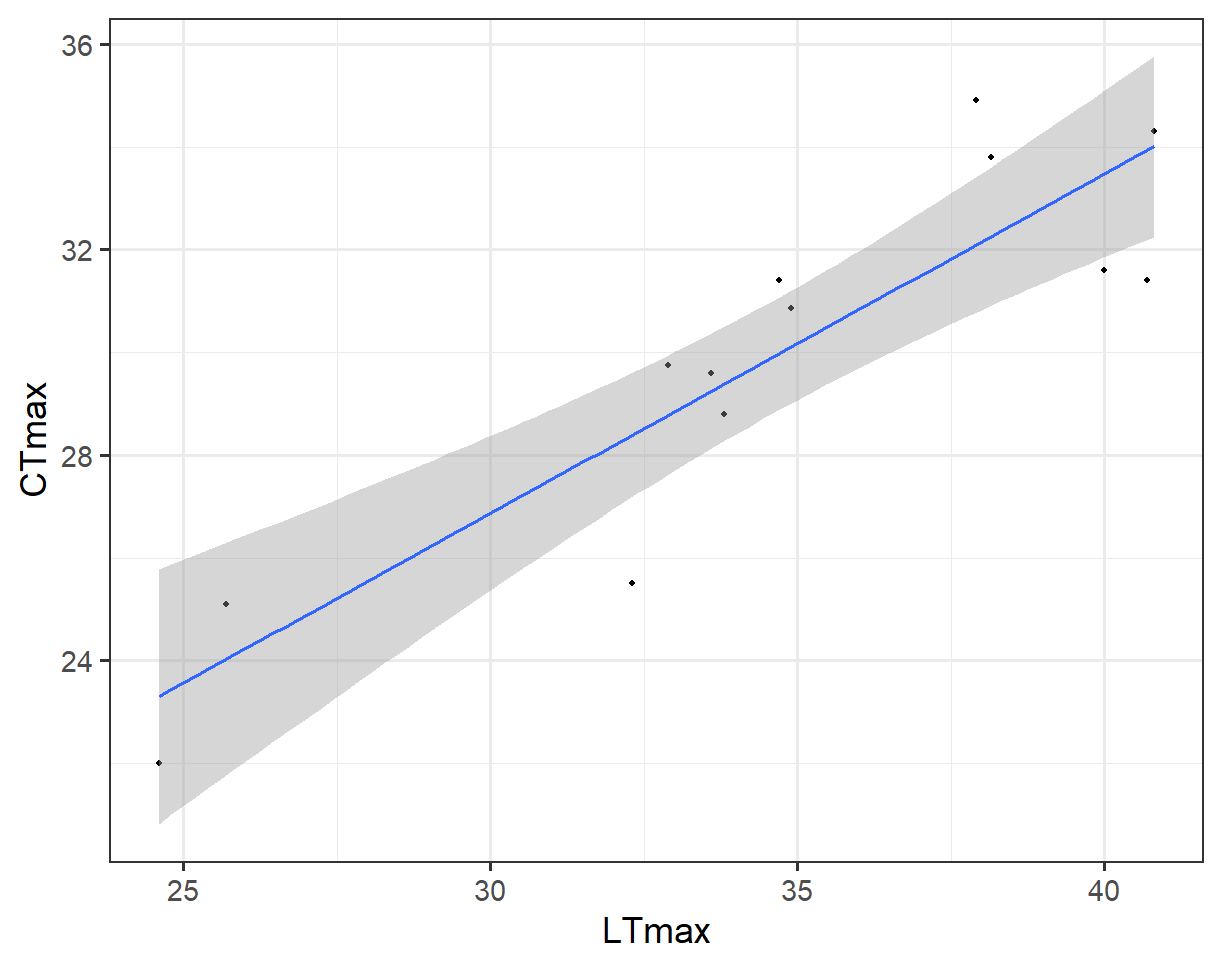
**Figure A2**. The fitted relationship (r = 0.8, p<<0.001) between LTmax and CTmax, which was used to correct the LTmax values to a comparable metric to CTmax, overlaid on the raw data points (n =13).

2.1.1.3. Derivation of excess mortality

To derive excess mortality from the tolerance values of each functional group, a linear relationship between the optimum and the maximum was assumed, based upon the known shape of thermal performance curves (**Figure A3**). Previous studies have observed a relationship between temperature approaching thermal maxima and mortality in selected organisms (Cicchino et al., 2023; Desforges et al., 2023).

This linear relationship between an optimum at which no excess mortality beyond the natural background mortality occurs and the thermal limit, at which complete mortality occurs, is represented in the following equation:

$excess mortality \% = \frac{100}{(thermal maximum - optimum)}\times(excess temperature)$	(2)

Where excess temperature equals the difference between the stress temperature and the optimum.

This relationship is depicted graphically in **Figure A4**. An optimum of 15 °C was assumed for all functional groups, in accordance with recent reviews of thermal optima for freshwater invertebrates (Sundermann et al., 2022; Tomczyk et al., 2022).

Since all temperature tolerance assays occurred over an acute time period (typically from 2 to 24 hours), the excess mortality (in percent) was translated into a daily excess death rate for implementation to the model. Thus the excess mortality was assumed to occur per day.

The excess mortalities for each functional group are reported in **Table A2**.

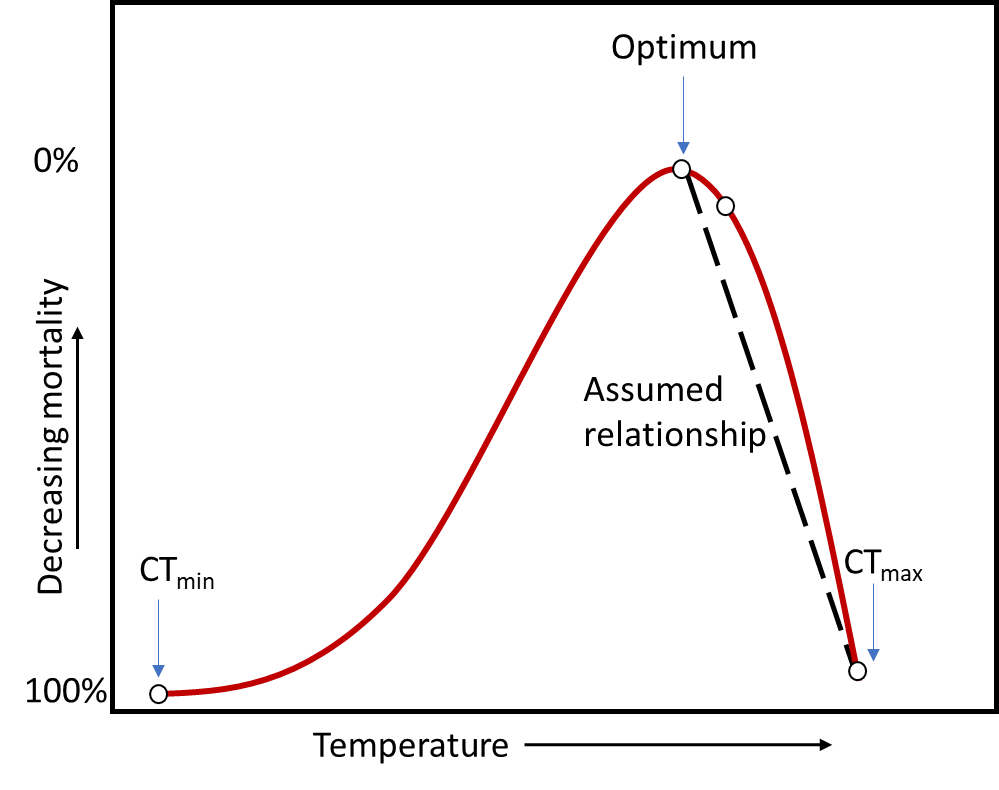

**Figure A3**. A generalized thermal performance curve including the assumed linear relationship of mortality and temperature between the temperature optimum and CTmax. The calculation is depicted graphically in **Figure A5**.

**Table A2**. Excess mortalities (%) computed for each functional group at chosen stressor temperatures of 18.5 °C (3.5 °C above the optimum of 15 °C) and 25 °C (10 °C above the optimum of 15 °C).

|  | **Low temperature stressor** | **High temperature stressor** |
| --- | --- | --- |
| **Functional group** | **18.5** °**C** | **25** °**C** |
| Fish | 24.57% | 70.19% |
| Detritivore | 22.26% | 63.59% |
| Shredder | 22.11% | 63.16% |
| Invertebrate predator | 18.69% | 53.4% |
| Grazer | 18.50% | 52.87% |

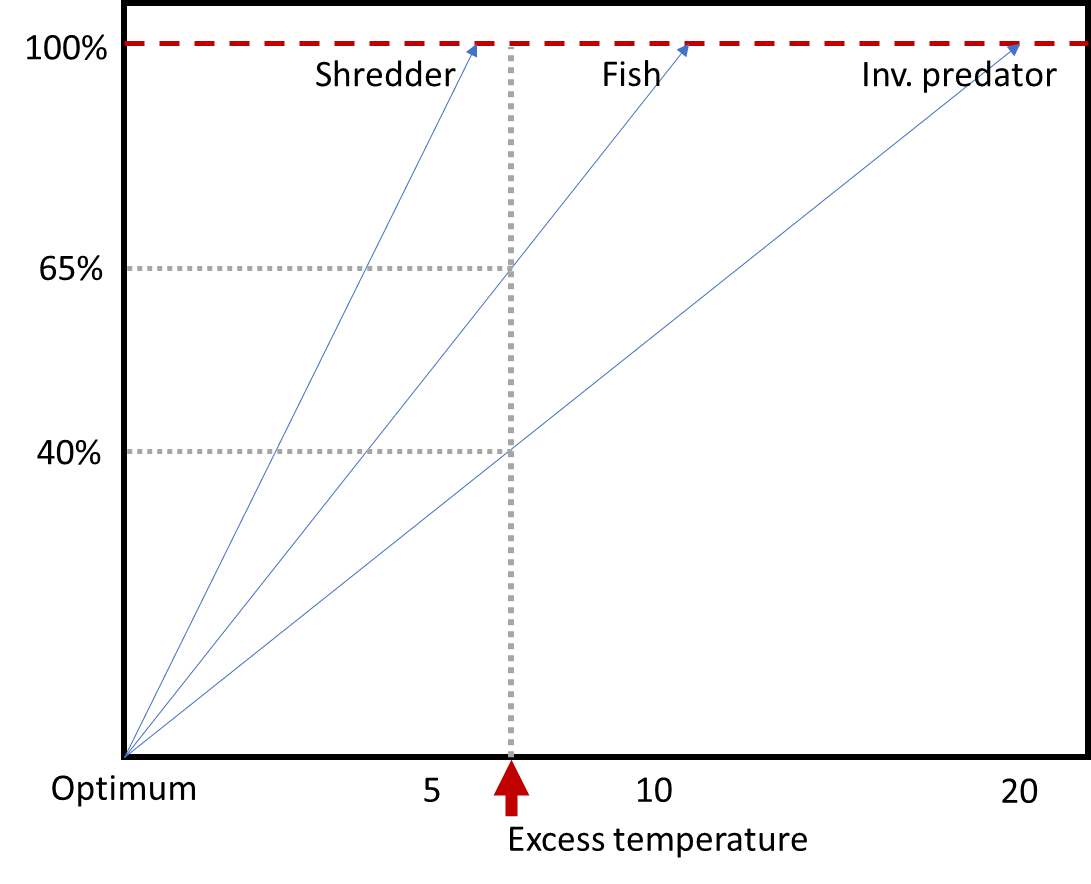

**Figure A4**. A graphical example of the calculation of excess mortality at a chosen excess temperature, marked by the red arrow, for groups with different thermal maximum values. The excess temperature is equal to the difference between the stress temperature and the optimum. In this example, the stress temperature is above the thermal limit of the shredder, resulting in 100% excess mortality due to the stress. Excess mortality is 65% for the fish and 40% for the invertebrate predator at this stress temperature. These values are an example of the calculation and do not reflect the real values used in the subsequent simulations; axes are not to scale.

2.1.2. Salinity tolerance
       2.1.2.1. Data acquisition and aggregation

Insufficient data was available for temperate taxa residing in central European freshwaters to assign data to each functional group. Instead, the general sensitivity of fish and invertebrates to salinity was derived based on freshwater taxa covered by the EPA ECOTOX database.

LC50s for acute tests of NaCl salt exposure with durations between 12 and 168 hours were queried using StandarTox, a cleaned and harmonized version of the EPA ECOTOX database that includes additional taxonomic and biological information (Olker et al., 2022; Scharmüller et al., 2020). Only tests for freshwater fish and invertebrates were included. The geometric mean of the LC50s for each taxon were calculated when multiple values were present, resulting in an average LC50 for each species. LC50s were available for 67 freshwater taxa, 53 invertebrate species and 14 fish species. Species sensitivity distributions were fit to all taxa together to visualize relative tolerance, as well as separately to each group to compute the excess mortality at chosen salinity levels for the stressor scenarios. Species sensitivity distributions were fitted and plotted in R version 4.2.1 (R Core Team, 2022).

The general differences in the salinity tolerances are visualized in **Figure A5**. The fish taxa are clustered towards the top, more tolerant, fraction of the distribution, while the invertebrates are clustered near the bottom and middle.

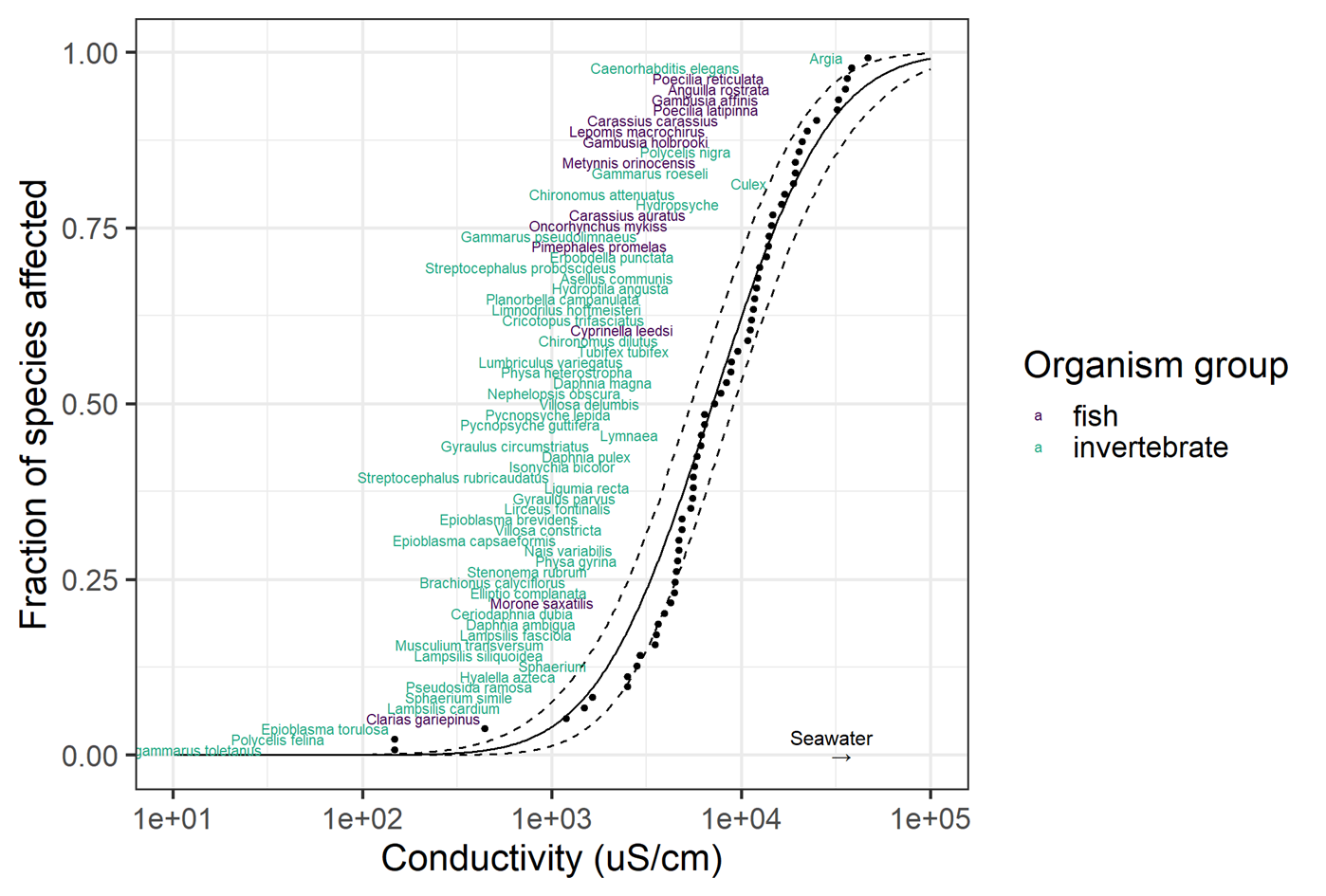

**Figure A5**. Species sensitivity for freshwater ectotherms (n = 67); fish (in purple) and invertebrates (in blue). The clustering of freshwater fish species near the top of the distribution indicate their higher tolerance to saline conditions compared to invertebrates.

2.1.2.2. Derivation of excess mortality

Excess mortality at the chosen salinity levels of 200 and 600 mg/L was derived by calculating the fraction affected at the respective salinity levels in each distribution, depicted in **Figures A6 & A7** with the red points. All invertebrate functional groups had the same excess mortality for salinity at each stressor level. Fish had negligible mortality at the chosen salinity stressor levels. The affected fraction was assumed to represent a daily excess death rate, as for temperature, as the data used came from acute assays.

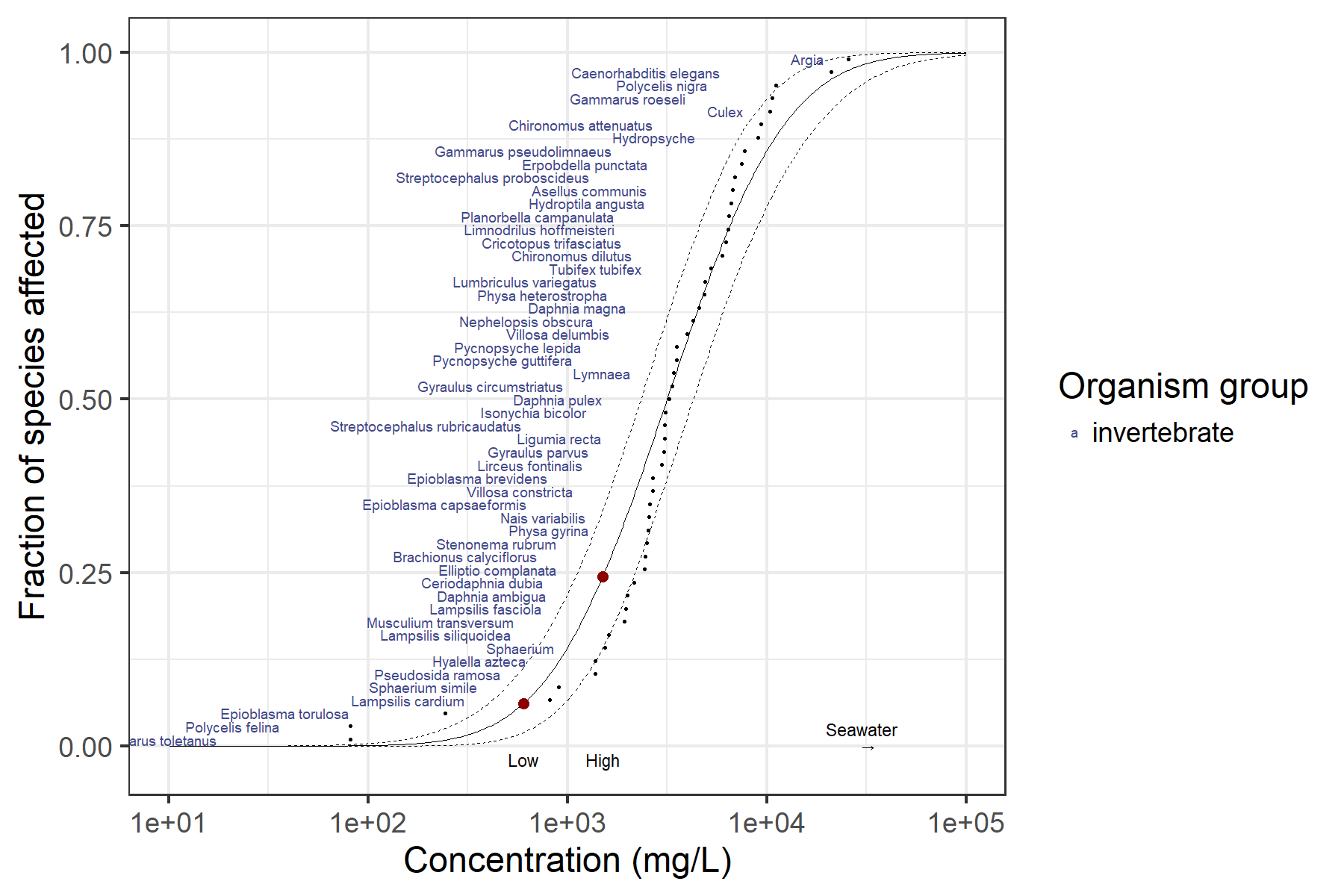

**Figure A6**. Species sensitivity distribution for acute LC50 of NaCl of freshwater invertebrates (n = 53). Data was queried from StandarTox and originated from the EPA ECOTOX database (Olker et al., 2022; Scharmüller et al., 2020). The chosen stressor levels for low (200 mg/L) and high (600 mg/L) salinity scenarios in this study are marked with red points.

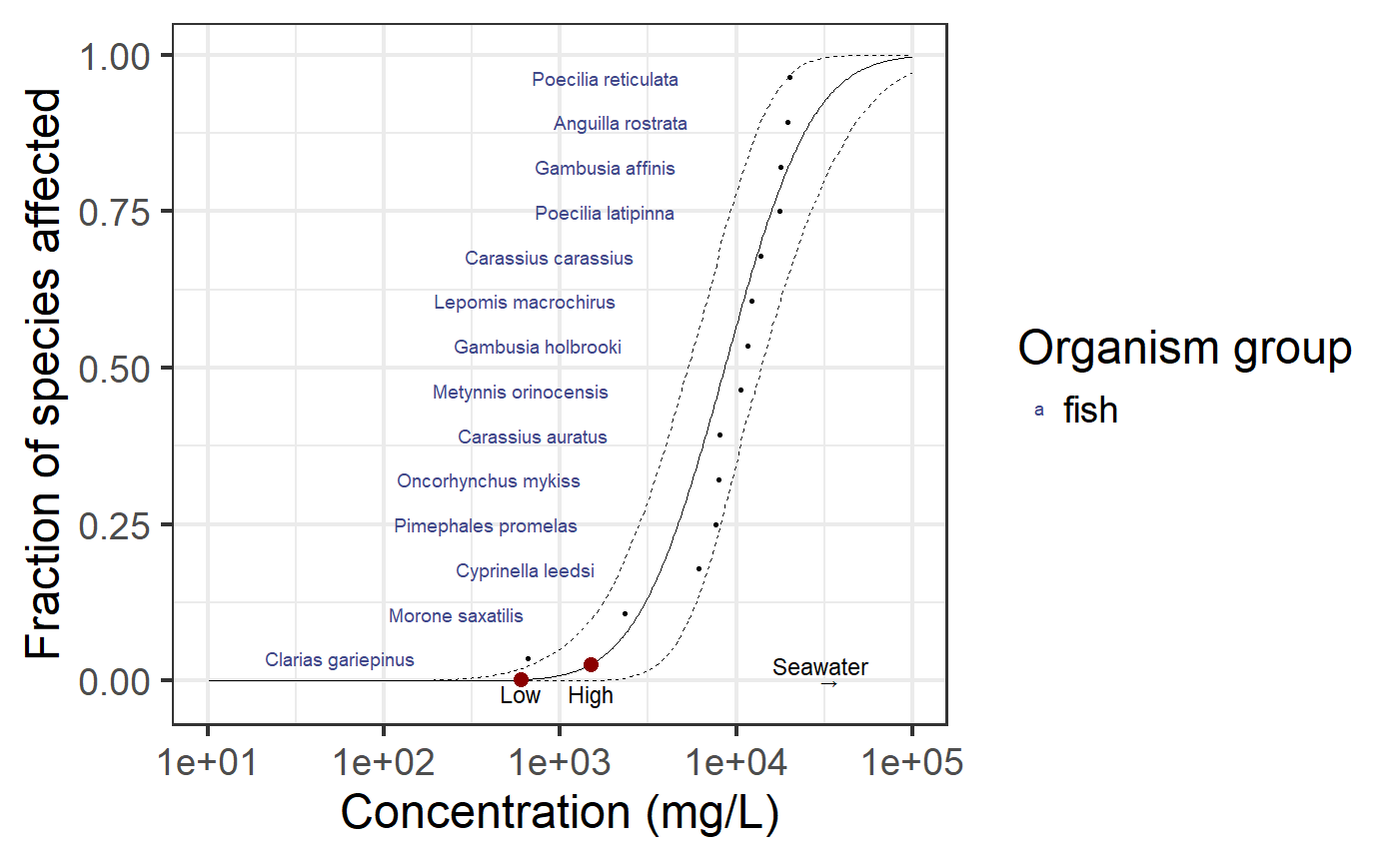

**Figure A7**. Species sensitivity distribution for acute LC50 of NaCl of freshwater fish (n = 14). Data was queried from StandarTox and originated from the EPA ECOTOX database (Olker et al., 2022; Scharmüller et al., 2020). The chosen stressor levels for low (600 mg/L) and high (1500 mg/L) salinity scenarios in this study are marked with red points.

**Table A3**. Excess mortality (rate per day) due to salinity for fish and invertebrate functional groups

| **Group** | **600 mg/L NaCl** | **1500 mg/L NaCl** |
| --- | --- | --- |
| **Fish** | 0.0016 | 0.2441 |
| **Shredder** | 0.061 | 0.0266 |
| **Invertebrate predator** | 0.061 | 0.2441 |
| **Grazer** | 0.061 | 0.2441 |
| **Detritivore** | 0.061 | 0.2441 |

2.2. Implementing death rates in scenarios

To implement death rates into the simulations, excess mortality was added to the natural mortality rate for each functional group. For multiple stressor scenarios, the single stressor mortalities were added together, on top of the natural mortality rate. Excess death rates for each group implemented in simulation scenarios, as well as the natural mortality for each group, are reported in **Table A4**.

The total effect was computed by adding all excess mortalities for each scenario; this was standardized to a value between 0 and 1 to obtain the effect strength for each scenario.

**Table A4**. Excess mortalities implemented in simulation scenarios. Groups represented by Sh = shredder, F = fish, IP = invertebrate predator, G = grazer, D = detritivore.

| **Scenario** | | | | | | | | | | | | |
| --- | --- | --- | --- | --- | --- | --- | --- | --- | --- | --- | --- | --- |
|  | **1** | **2** | **3** | **4** | **5** | **6** | **7** | **8** | **9** | **10** | **11** | **BG** |
| **Group** | 600 mgL | 1500mgL | 18.5C | 25C | 600 mgL + 18.5C | 1500mgL + 18.5C | 600 mgL + 25C | 1500 mgL + 25C | 200 mgL | 22C | 5300 mgL | No stress |
| **Sh** | 0.0610 | 0.2441 | 0.2211 | 0.6316 | 0.2821 | 0.4652 | 0.6926 | 0.8757 | 0.0051 | 0.4421 | 0.6853 | 0.0020 |
| **F** | 0.0016 | 0.0266 | 0.2457 | 0.7019 | 0.2473 | 0.2723 | 0.7035 | 0.7285 | 0.0000 | 0.4913 | 0.2966 | 0.0040 |
| **IP** | 0.0610 | 0.2441 | 0.1869 | 0.5340 | 0.2479 | 0.4310 | 0.5950 | 0.7781 | 0.0051 | 0.3738 | 0.6853 | 0.0025 |
| **G** | 0.0610 | 0.2441 | 0.1850 | 0.5287 | 0.2460 | 0.4291 | 0.5897 | 0.7728 | 0.0051 | 0.3701 | 0.6853 | 0.0020 |
| **D** | 0.0610 | 0.2441 | 0.2226 | 0.6359 | 0.2836 | 0.4667 | 0.6969 | 0.8800 | 0.0051 | 0.4451 | 0.6853 | 0.0035 |
| **total effect** | 0.2456 | 1.0030 | 1.0612 | 3.0321 | 1.3068 | 2.0642 | 3.2777 | 4.0351 | 0.0204 | 2.1224 | 3.0376 |  |
| **Avg effect** | 0.0491 | 0.2006 | 0.2122 | 0.6064 | 0.2614 | 0.4128 | 0.6555 | 0.8070 | 0.0041 | 0.4245 | 0.6075 |  |
| **Stand. effect** | 0.0491 | 0.2006 | 0.2122 | 0.6064 | 0.2614 | 0.4128 | 0.6555 | 0.8070 | 0.0041 | 0.4245 | 0.6075 |  |

2.3. Tolerance in community context

The relative tolerance in the community context was assessed during and after the stressor phase using two different metrics - the rate of change in the first stressor phase half and the days to extinction.

2.3.1. During the stressor phase

To assess the relative tolerance of different groups during the stressor phase, the rate of decrease within the first 7 days of the 14 day stressor phase was quantified, as referenced to the same time period in an unstressed system. The rate of decrease relative to the unstressed system was calculated as follows; first, the relative biomass was determined for each time step:

Relative change in biomass = biomass stressed - biomass unstressed/biomass unstressed

Then, the relative change was averaged across the first 7 days of the stressor phase:

$relative change = (\sum_{i = 1}^{7} {relative biomass}_{i})\div7$	(3)

This was done for each functional group affected by stressor impact. The first 7 days were chosen over the whole 14 day period because all impacted functional groups were still present within the first seven days. After 7 days, functional groups began to go extinct. Once a functional group went extinct, the relative decrease rate remained at -100%, even though no further biomass was lost, which would skew the results after extinction. Unusually, the detritivore showed a positive rate of change during the stressor phase, despite the high excess mortality. Relative changes in biomass during the first half of the stressor phase for each group under a selection of scenarios are reported in **Table A5**.

**Table A5**. Relative rates of change for each functional group during the first 7 days of the stressor phase in units of % per day. Groups represented by Sh = shredder, F = fish, IP = invertebrate predator, G = grazer, D = detritivore.

|  | **Scenario** | | | | | | | | | | |
| --- | --- | --- | --- | --- | --- | --- | --- | --- | --- | --- | --- |
| **Group** | **1** | **2** | **3** | **4** | **5** | **6** | **7** | **8** | **9** | **10** | **11** |
| **Sh** | -20.87 | -54.76 | -54.33 | -84.39 | -62.06 | -75.54 | -86.21 | -89.80 | -1.98 | -75.35 | -86.00 |
| **G** | -18.12 | -51.48 | -45.00 | -78.49 | -54.46 | -71.20 | -81.26 | -86.56 | -1.63 | -68.02 | -84.68 |
| **D** | -2.91 | -13.69 | 5.46 | -41.84 | -1.86 | -23.47 | -49.57 | -66.10 | -0.22 | -17.13 | -53.60 |
| **IP** | -20.97 | -54.92 | -48.46 | -80.19 | -57.71 | -73.40 | -82.68 | -87.48 | -1.99 | -70.56 | -85.63 |
| **F** | -0.76 | -10.27 | -57.96 | -86.09 | -58.19 | -61.26 | -86.14 | -86.79 | -0.02 | -78.25 | -63.96 |

2.3.2. Second half and post-stressor phase

To assess the relative tolerance of groups after the stressor phase, the days to extinction were quantified for each group under various scenarios. These are reported in **Table A6**. This metric was used for Spearman calculations only in cases where more than 2 extinctions occurred.

**Table A6**. Days to extinction for each group under stressor scenarios where more than 2 extinctions occurred.

|  | **Scenario** | | | |
| --- | --- | --- | --- | --- |
| **Group** | **4** | **7** | **8** | **11** |
| **Shredder** | 7 | 6 | 5 | 6 |
| **Fish** | 11 | 13 | 11 | Not extinct |
| **Invertebrate predator** | 14 | 12 | 10 | 11 |
| **Grazer** | 14 | 11 | 10 | 11 |
| **Detritivore** | Not extinct | Not extinct | Not extinct | Not extinct |

3. **References**

Beitinger, T. L., Bennett, W. A., & Mccauley, R. W. (2000). Temperature tolerances of North American freshwater fishes exposed to dynamic changes in temperature. *Environmental Biology of Fishes*, *58*, 237–275.

Bennett, J. M. (2018). *Data Descriptor: GlobTherm, a global database on thermal tolerances for aquatic and terrestrial organisms Background & Summary*. https://doi.org/10.1038/sdata.2018.22

Cameron, A. T., & Brownlee, T. I. (1915). THE UPPER LIMIT OF TEMPERATURE COMPATIBLE WITH LIFE IN THE FROG. *Quarterly Journal of Experimental Physiology*, *9*(3), 247–260. https://doi.org/10.1113/expphysiol.1915.sp000206

Cereja, R. (2020). Critical thermal maxima in aquatic ectotherms. *Ecological Indicators*, *119*. https://doi.org/10.1016/j.ecolind.2020.106856

Chatterjee, N., Pal, A. K., Manush, S. M., Das, T., & Mukherjee, S. C. (2004). Thermal tolerance and oxygen consumption of Labeo rohita and Cyprinus carpio early fingerlings acclimated to three different temperatures. *Journal of Thermal Biology*, *29*(6), 265–270. https://doi.org/10.1016/j.jtherbio.2004.05.001

Cicchino, A. S., Ghalambor, C. K., & Funk, W. C. (2023). Linking critical thermal maximum to mortality from thermal stress in a cold-water frog. *Biology Letters*, *19*. https://doi.org/10.1098/rsbl.2023.0106

Coutant, C. C., & Brook, A. J. (1970). Biological aspects of thermal pollution I. Entrainment and discharge canal effects∗. *C R C Critical Reviews in Environmental Control*, *1*(1–4), 341–381. https://doi.org/10.1080/10643387009381570

Coutant, C. C., & Goodyear, C. P. (1972). Thermal Effects. *Journal (Water Pollution Control Federation)*, *44*(6), 1250–1294.

Cowles, R. B., & Bogert, C. M. (1944). A preliminary study of the thermal requirements of desert reptiles. *Bulletin of the AMNH*, *83*.

Dallas, H. F., & Ketley, Z. A. (2011). Upper thermal limits of aquatic macroinvertebrates: Comparing critical thermal maxima with 96-LT50 values. *Journal of Thermal Biology*, *36*(6), 322–327. https://doi.org/10.1016/j.jtherbio.2011.06.001

Davenport, C. B., & Castle, W. E. (1895). *Acclimatization of Organisms to High Temperatures 1)*.

Davy, J. (1855). *Some Observations on the Ova of the Salmon, in relation to the distribution of Species* [Personal communication].

Desforges, J. E., Birnie-Gauvin, K., Jutfelt, F., Gilmour, K. M., Eliason, E. J., Dressler, T. L., McKenzie, D. J., Bates, A. E., Lawrence, M. J., Fangue, N., & Cooke, S. J. (2023). The ecological relevance of critical thermal maxima methodology for fishes. *Journal of Fish Biology*, *102*(5), 1000–1016. https://doi.org/10.1111/jfb.15368

Gillmann, S. M., Hering, D., & Lorenz, A. W. (2023). Habitat development and species arrival drive succession of the benthic invertebrate community in restored urban streams. *Environmental Sciences Europe*, *35*(1), 49. https://doi.org/10.1186/s12302-023-00756-x

Hoppe-Seyler, F. (1875). Ueber die obere Temperaturgrenze des Lebens. *Archiv für die Gesammte Physiologie des Menschen und der Thiere*, *11*(1), 113–121. https://doi.org/10.1007/BF01659294

Jørgensen, L. B., Malte, H., Ørsted, M., Klahn, N. A., & Overgaard, J. (2021). A unifying model to estimate thermal tolerance limits in ectotherms across static, dynamic and fluctuating exposures to thermal stress. *Scientific Reports 2021 11:1*, *11*(1), 1–14. https://doi.org/10.1038/s41598-021-92004-6

Leiva, F. P., Calosi, P., & Verberk, W. C. E. P. (2019). Scaling of thermal tolerance with body mass and genome size in ectotherms: A comparison between water-and air-breathers. *Philosophical Transactions of the Royal Society B*, *374*. https://doi.org/10.1098/rstb.2019.0035

Moog, O. (1995). *Fauna Aquatica Austriaca*. Bundesministerium für Land- und Forstwirtschaft, Umwelt und Wasserwirtschaft. https://www.zobodat.at/pdf/Fauna-Aquatica-Austriaca_2002_0001-0670.pdf

OECD. (2002). *Detailed Review Paper on Aquatic Testing Methods for Pesticides and Industrial Chemicals*. OECD. https://doi.org/10.1787/9789264078291-en

Olker, J. H., Elonen, C. M., Pilli, A., Anderson, A., Kinziger, B., Erickson, S., Skopinski, M., Pomplun, A., LaLone, C. A., Russom, C. L., & Hoff, D. (2022). The ECOTOXicology Knowledgebase: A Curated Database of Ecologically Relevant Toxicity Tests to Support Environmental Research and Risk Assessment. *Environmental Toxicology and Chemistry*, *41*(6), 1520–1539. https://doi.org/10.1002/etc.5324

Olsen, D. A., Tremblay, L., Clapcott, J., & Holmes, R. (2012). *Water temperature criteria for native aquatic biota* (2012/036). Auckland Council.

Orr, J. A., Vinebrooke, R. D., Jackson, M. C., Kroeker, K. J., Kordas, R. L., Mantyka-Pringle, C., van den Brink, P. J., de Laender, F., Stoks, R., Holmstrup, M., Matthaei, C. D., Monk, W. A., Penk, M. R., Leuzinger, S., Schäfer, R. B., & Piggott, J. J. (2020). Towards a unified study of multiple stressors: Divisions and common goals across research disciplines. *Proceedings of the Royal Society B: Biological Sciences*, *287*(1926). https://doi.org/10.1098/rspb.2020.0421

R Core Team. (2022). *R: A language and environment for  statistical computing* [R]. R Foundation for Statistical Computing. https://www.R-project.org/

Rezende, E. L., & Bozinovic, F. (2019). *Thermal performance across levels of biological organization*. https://doi.org/10.1098/rstb.2018.0549

Rezende, E. L., Castañeda, L. E., & Santos, M. (2014). Tolerance landscapes in thermal ecology. *Functional Ecology*, *28*(4), 799–809. https://doi.org/10.1111/1365-2435.12268

Ritz, C., Baty, F., Streibig, J. C., & Gerhard, D. (2015). Dose-Response Analysis Using R. *PLOS ONE*, *10*(12).

Schäfer, R. B., Jackson, M., Juvigny-Khenafou, N., Osakpolor, S. E., Posthuma, L., Schneeweiss, A., Spaak, J., & Vinebrooke, R. (2023). Chemical Mixtures and Multiple Stressors: Same but Different? *Environmental Toxicology and Chemistry*, *42*(9), 1915–1936. https://doi.org/10.1002/etc.5629

Scharmüller, A., Schreiner, V. C., & Schäfer, R. B. (2020). Standartox: Standardizing Toxicity Data. *Data*, *5*(2). https://doi.org/10.5281/zenodo.3785031

Schmidt-Kloiber, A., & Hering, D. (2015). Www.freshwaterecology.info – An online tool that unifies, standardises and codifies more than 20,000 European freshwater organisms and their ecological preferences. *Ecological Indicators*, *53*, 271–282. https://doi.org/10.1016/j.ecolind.2015.02.007

Sinclair, B. J., Marshall, K. E., Sewell, M. A., Levesque, D. L., Willett, C. S., Slotsbo, S., Dong, Y., Harley, C. D. G., Marshall, D. J., Helmuth, B. S., & Huey, R. B. (2016). Can we predict ectotherm responses to climate change using thermal performance curves and body temperatures? *Ecology Letters*, *19*(11), 1372–1385. https://doi.org/10.1111/ELE.12686

Spallanzani. (1777). *Opuscules de Physique, Animale et vegetale*. Chez Barthelemi Chirol.

Straalen, N. M. V. (2003). Ecotoxicology Becomes Stress Ecology. *Environmental Science & Technology*, *37*(17), 324A-330A. https://doi.org/10.1017/CBO9781107415324.004

Suemoto, T., Kawai, K., & Imabayashi, H. (2004). A comparison of desiccation tolerance among 12 species of chironomid larvae. *Hydrobiologia*, *515*(1), 107–114. https://doi.org/10.1023/B:HYDR.0000027322.11005.20

Sundermann, A., Müller, A., & Halle, M. (2022). A new index of a water temperature equivalent for summer respiration conditions of benthic invertebrates in rivers as a bio-indicator of global climate change. *Limnologica*, *95*. https://doi.org/10.1016/j.limno.2022.125980

Tachet, H., Richoux, P., Bournaud, M., & Usseglio-Polatera, P. (2010). *Invertébrés d’eau douce: Systématique, biologie, écologie* (Vol. 15). CNRS éditions Paris.

*Technical Guidance for Deriving Environmental Quality Standards* (Guidance Document No. 27). (2018). EU Commission.

*Temperature: Water Quality Standards Criteria Summaries: A Compilation of State/Federal Criteria* (EPA/440/5-88-023). (1988). USEPA. https://nepis.epa.gov/Exe/ZyNET.exe/00001NPM.txt?ZyActionD=ZyDocument&Client=EPA&Index=1986%20Thru%201990&Docs=&Query=&Time=&EndTime=&SearchMethod=1&TocRestrict=n&Toc=&TocEntry=&QField=&QFieldYear=&QFieldMonth=&QFieldDay=&UseQField=&IntQFieldOp=0&ExtQFieldOp=0&XmlQuery=&File=D%3A%5CZYFILES%5CINDEX%20DATA%5C86THRU90%5CTXT%5C00000001%5C00001NPM.txt&User=ANONYMOUS&Password=anonymous&SortMethod=h%7C-&MaximumDocuments=1&FuzzyDegree=0&ImageQuality=r75g8/r75g8/x150y150g16/i425&Display=hpfr&DefSeekPage=x&SearchBack=ZyActionL&Back=ZyActionS&BackDesc=Results%20page&MaximumPages=1&ZyEntry=5

Tomczyk, N. J., Rosemond, A. D., Rogers, P. A., & Cummins, C. S. (2022). Thermal traits of freshwater macroinvertebrates vary with feeding group and phylogeny. *Freshwater Biology*, *67*(11), 1994–2003. https://doi.org/10.1111/fwb.13992

Vernon, H. M. (1899). *The death temperature of certain marine organisms*.

Walshe, B. M. (1948). The Oxygen Requirements and Thermal Resistance of Chironomid Larvae from Flowing and from Still Waters. *Journal of Experimental Biology*, *25*(1), 35–44. https://doi.org/10.1242/jeb.25.1.35
