## Supplementary material for "Putting the Asymmetric Response Concept to the test: modeling multiple stressor exposure and release in a stream food web": Supp Appendix B

The food web model builds on a basic food web model of DeAngelis (**Figure B1**), which has been extended with several functional groups.

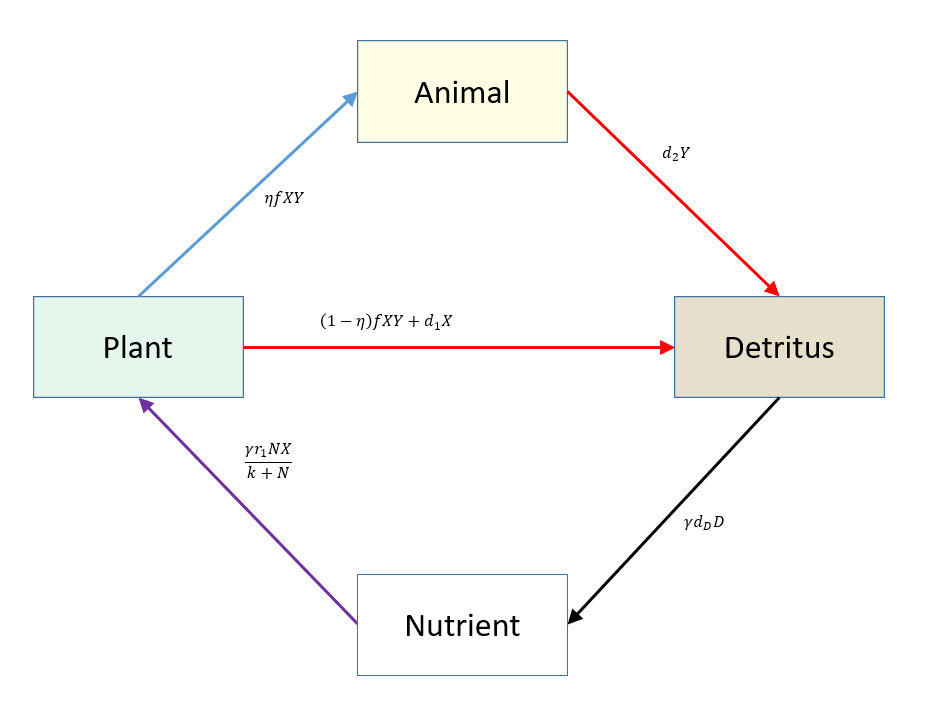

**Figure B1**. Basic food web model from DeAngelis (1992). The plant compartment takes up a limiting nutrient (purple arrow). Animals assimilate a portion η of the consumed plant material (blue arrow, fXY). The non-assimilated portion (1-η)fXY becomes detritus, along with dead plant biomass d_1_X (horizontal red arrow). In addition, dead animal material d_2_Y becomes detritus. Detritus is remineralized (black arrow), replenishing the nutrient, thereby closing the cycle.

All simulations were performed using Matlab Version R2023b (Mathworks) using the following equations (E.1)-(E.16). The parameter values used with this model were fitted to yield a model in equilibrium. All parameters and Matlab settings are presented in Table B1 and Table B2.

Equations:

(E.1) Periphyton:

$$\frac{dP}{dt}=\frac{gamma*r_{1}*P*N}{k_{1}+N}$$

$$-n_{1}*\frac{a_{1}*P^{q_{1}\left( 1 \right)}*G}{1+a_{1}*h_{1}*P^{q_{1}\left( 1 \right)}}-\left( 1-n_{1} \right)*\frac{a_{1}*P^{q_{1}\left( 1 \right)}*G}{1+a_{1}*h_{1}*P^{q_{1}\left( 1 \right)}}$$

$$-s_{1}+f_{1}\left( 1 \right)*{ds}_{1}\left( 1 \right)+f_{2}\left( 1 \right)*{ds}_{2}\left( 1 \right)*P$$

(E.2) Grazer:

$$\frac{dG}{dt}=n_{1}*\frac{a_{1}*P^{q_{1}\left( 1 \right)}*G}{1+a_{1}*h_{1}*P^{q_{1}\left( 1 \right)}}$$

$$-n_{5}*\frac{a_{5}*G^{q_{4}\left( 1 \right)}*IP}{1+a_{5}*h_{5}*G^{q_{4}\left( 1 \right)}+a_{6}*h_{6}*D^{q_{4}\left( 2 \right)}+a_{7}*h_{7}*{Sh}^{q_{4}\left( 3 \right)}}$$

$$-\left( 1-n_{5} \right)*\frac{a_{5}*G^{q_{4}\left( 1 \right)}*IP}{1+a_{5}*h_{5}*G^{q_{4}\left( 1 \right)}+a_{6}*h_{6}*D^{q_{4}\left( 2 \right)}+a_{7}*h_{7}*{Sh}^{q_{4}\left( 3 \right)}}$$

$$-n_{8}*\frac{a_{8}*G^{q_{5}\left( 1 \right)}*FP}{1+a_{8}*h_{8}*G^{q_{5}\left( 1 \right)}+a_{9}*h_{9}*D^{q_{5}\left( 2 \right)}+a_{10}*h_{10}*{Sh}^{q_{5}\left( 3 \right)}+a_{11}*h_{11}*{IP}^{q_{5}\left( 4 \right)}}$$

$$-\left( 1-n_{8} \right)*\frac{a_{8}*G^{q_{5}\left( 1 \right)}*FP}{1+a_{8}*h_{8}*G^{q_{5}\left( 1 \right)}+a_{9}*h_{9}*D^{q_{5}\left( 2 \right)}+a_{10}*h_{10}*{Sh}^{q_{5}\left( 3 \right)}+a_{11}*h_{11}*{IP}^{q_{5}\left( 4 \right)}}$$

$$-s_{2}+f_{1}\left( 2 \right)*{ds}_{1}\left( 2 \right)+f_{2}\left( 2 \right)*{ds}_{2}\left( 2 \right)*G$$

(E.3) Detritivore:

$$\frac{dD}{dt}=n_{2}*\frac{a_{2}*{PD}^{q_{2}\left( 1 \right)}*D}{1+a_{2}*h_{2}*{PD}^{q_{2}\left( 1 \right)}+a_{3}*h_{3}*{AD}^{q_{2}\left( 2 \right)}}+n_{3}*\frac{a_{3}*{AD}^{q_{2}\left( 2 \right)}*D}{1+a_{2}*h_{2}*{PD}^{q_{2}\left( 1 \right)}+a_{3}*h_{3}*{AD}^{q_{2}\left( 2 \right)}}$$

$$-n_{6}*\frac{a_{6}*D^{q_{4}\left( 2 \right)}*IP}{1+a_{5}*h_{5}*G^{q_{4}\left( 1 \right)}+a_{6}*h_{6}*D^{q_{4}\left( 2 \right)}+a_{7}*h_{7}*{Sh}^{q_{4}\left( 3 \right)}}$$

$$-\left( 1-n_{6} \right)*\frac{a_{6}*D^{q_{4}\left( 2 \right)}*IP}{1+a_{5}*h_{5}*G^{q_{4}\left( 1 \right)}+a_{6}*h_{6}*D^{q_{4}\left( 2 \right)}+a_{7}*h_{7}*{Sh}^{q_{4}\left( 3 \right)}}$$

$$-n_{9}*\frac{a_{9}*D^{q_{5}\left( 2 \right)}*FP}{1+a_{8}*h_{8}*G^{q_{5}\left( 1 \right)}+a_{9}*h_{9}*D^{q_{5}\left( 2 \right)}+a_{10}*h_{10}*{Sh}^{q_{5}\left( 3 \right)}+a_{11}*h_{11}*{IP}^{q_{5}\left( 4 \right)}}$$

$$-\left( 1-n_{9} \right)*\frac{a_{9}*D^{q_{5}\left( 2 \right)}*FP}{1+a_{8}*h_{8}*G^{q_{5}\left( 1 \right)}+a_{9}*h_{9}*D^{q_{5}\left( 2 \right)}+a_{10}*h_{10}*{Sh}^{q_{5}\left( 3 \right)}+a_{11}*h_{11}*{IP}^{q_{5}\left( 4 \right)}}$$

$$-s_{3}+f_{1}\left( 3 \right)*{ds}_{1}\left( 3 \right)+f_{2}\left( 3 \right)*{ds}_{2}\left( 3 \right)*D$$

(E.4) Shredder:

$$\frac{dSh}{dt}=n_{4}*\frac{a_{4}*{CPOM}^{q_{3}\left( 1 \right)}*Sh}{1+a_{4}*h_{4}*{CPOM}^{q_{3}\left( 1 \right)}}$$

$$-n_{7}*\frac{a_{7}*{Sh}^{q_{4}\left( 3 \right)}*IP}{1+a_{5}*h_{5}*G^{q_{4}\left( 1 \right)}+a_{6}*h_{6}*D^{q_{4}\left( 2 \right)}+a_{7}*h_{7}*{Sh}^{q_{4}\left( 3 \right)}}$$

$$-\left( 1-n_{7} \right)*\frac{a_{7}*{Sh}^{q_{4}\left( 3 \right)}*IP}{1+a_{5}*h_{5}*G^{q_{4}\left( 1 \right)}+a_{6}*h_{6}*D^{q_{4}\left( 2 \right)}+a_{7}*h_{7}*{Sh}^{q_{4}\left( 3 \right)}}$$

$$-n_{10}*\frac{a_{10}*{Sh}^{q_{5}\left( 3 \right)}*FP}{1+a_{8}*h_{8}*G^{q_{5}\left( 1 \right)}+a_{9}*h_{9}*D^{q_{5}\left( 2 \right)}+a_{10}*h_{10}*{Sh}^{q_{5}\left( 3 \right)}+a_{11}*h_{11}*{IP}^{q_{5}\left( 4 \right)}}$$

$$-\left( 1-n_{10} \right)*\frac{a_{10}*{Sh}^{q_{5}\left( 3 \right)}*FP}{1+a_{8}*h_{8}*G^{q_{5}\left( 1 \right)}+a_{9}*h_{9}*D^{q_{5}\left( 2 \right)}+a_{10}*h_{10}*{Sh}^{q_{5}\left( 3 \right)}+a_{11}*h_{11}*{IP}^{q_{5}\left( 4 \right)}}$$

$$-s_{4}+f_{1}\left( 4 \right)*{ds}_{1}\left( 4 \right)+f_{2}\left( 4 \right)*{ds}_{2}\left( 4 \right)*Sh$$

(E.5) invertebrate Predator:

$$\frac{dIP}{dt}=n_{5}*\frac{a_{5}*G^{q_{4}\left( 1 \right)}*IP}{1+a_{5}*h_{5}*G^{q_{4}\left( 1 \right)}+a_{6}*h_{6}*D^{q_{4}\left( 2 \right)}+a_{7}*h_{7}*{Sh}^{q_{4}\left( 3 \right)}}$$

$$+n_{6}\frac{a_{6}*D^{q_{4}\left( 2 \right)}*IP}{1+a_{5}*h_{5}*G^{q_{4}\left( 1 \right)}+a_{6}*h_{6}*D^{q_{4}\left( 2 \right)}+a_{7}*h_{7}*{Sh}^{q_{4}\left( 3 \right)}}$$

$$+n_{7}*\frac{a_{7}*{Sh}^{q_{4}\left( 3 \right)}*IP}{1+a_{5}*h_{5}*G^{q_{4}\left( 1 \right)}+a_{6}*h_{6}*D^{q_{4}\left( 2 \right)}+a_{7}*h_{7}*{Sh}^{q_{4}\left( 3 \right)}}$$

$$-n_{11}*\frac{a_{11}*{IP}^{q_{5}\left( 4 \right)}*FP}{1+a_{8}*h_{8}*G^{q_{5}\left( 1 \right)}+a_{9}*h_{9}*D^{q_{5}\left( 2 \right)}+a_{10}*h_{10}*{Sh}^{q_{5}\left( 3 \right)}+a_{11}*h_{11}*{IP}^{q_{5}\left( 4 \right)}}$$

$$-\left( 1-n_{11} \right)*\frac{a_{11}*{IP}^{q_{5}\left( 4 \right)}*FP}{1+a_{8}*h_{8}*G^{q_{5}\left( 1 \right)}+a_{9}*h_{9}*D^{q_{5}\left( 2 \right)}+a_{10}*h_{10}*{Sh}^{q_{5}\left( 3 \right)}+a_{11}*h_{11}*{IP}^{q_{5}\left( 4 \right)}}$$

$$-s_{5}+f_{1}\left( 5 \right)*{ds}_{1}\left( 5 \right)+f_{2}\left( 5 \right)*{ds}_{2}\left( 5 \right)*IP$$

(E.6) Fish:

$$\frac{dFP}{dt}=n_{8}*\frac{a_{8}*G^{q_{5}\left( 1 \right)}*FP}{1+a_{8}*h_{8}*G^{q_{5}\left( 1 \right)}+a_{9}*h_{9}*D^{q_{5}\left( 2 \right)}+a_{10}*h_{10}*{Sh}^{q_{5}\left( 3 \right)}+a_{11}*h_{11}*{IP}^{q_{5}\left( 4 \right)}}$$

$$+n_{9}*\frac{a_{9}*D^{q_{5}\left( 2 \right)}*FP}{1+a_{8}*h_{8}*G^{q_{5}\left( 1 \right)}+a_{9}*h_{9}*D^{q_{5}\left( 2 \right)}+a_{10}*h_{10}*{Sh}^{q_{5}\left( 3 \right)}+a_{11}*h_{11}*{IP}^{q_{5}\left( 4 \right)}}$$

$$+n_{10}*\frac{a_{10}*{Sh}^{q_{5}\left( 3 \right)}*FP}{1+a_{8}*h_{8}*G^{q_{5}\left( 1 \right)}+a_{9}*h_{9}*D^{q_{5}\left( 2 \right)}+a_{10}*h_{10}*{Sh}^{q_{5}\left( 3 \right)}+a_{11}*h_{11}*{IP}^{q_{5}\left( 4 \right)}}$$

$$+n_{11}*\frac{a_{11}*{IP}^{q_{5}\left( 4 \right)}*FP}{1+a_{8}*h_{8}*G^{q_{5}\left( 1 \right)}+a_{9}*h_{9}*D^{q_{5}\left( 2 \right)}+a_{10}*h_{10}*{Sh}^{q_{5}\left( 3 \right)}+a_{11}*h_{11}*{IP}^{q_{5}\left( 4 \right)}}$$

$$-s_{6}+f_{1}\left( 6 \right)*{ds}_{1}\left( 6 \right)+f_{2}\left( 6 \right)*{ds}_{2}\left( 6 \right)*FP$$

(E.7) Fungi:

$$\frac{dF}{dt} = n_{14}*\frac{a_{14}*{CPOM}^{q_{6}\left( 3 \right)}*F}{1+a_{14}*h_{14}*{CPOM}^{q_{6}\left( 3 \right)}+a_{15}*h_{15}*{FPOM}^{q_{6}\left( 4 \right)}+a_{16}*h_{16}*{OM}^{q_{6}\left( 5 \right)}+a_{17}*h_{17}*{DF}^{q_{6}\left( 6 \right)}}$$

$$+n_{15}*\frac{a_{15}*{FPOM}^{q_{6}\left( 4 \right)}*F}{1+a_{14}*h_{14}*{CPOM}^{q_{6}\left( 3 \right)}+a_{15}*h_{15}*{FPOM}^{q_{6}\left( 4 \right)}+a_{16}*h_{16}*{OM}^{q_{6}\left( 5 \right)}+a_{17}*h_{17}*{DF}^{q_{6}\left( 6 \right)}}$$

$$+n_{16}*\frac{c_{16}*{OM}^{q_{6}\left( 5 \right)}*F}{1+a_{14}*h_{14}*{CPOM}^{q_{6}\left( 3 \right)}+a_{15}*h_{15}*{FPOM}^{q_{6}\left( 4 \right)}+a_{16}*h_{16}*{OM}^{q_{6}\left( 5 \right)}+a_{17}*h_{17}*{DF}^{q_{6}\left( 6 \right)}}$$

$$+n_{17}*\frac{a_{17}*{DF}^{q_{6}\left( 6 \right)}*F}{1+a_{14}*h_{14}*{CPOM}^{q_{6}\left( 3 \right)}+a_{15}*h_{15}*{FPOM}^{q_{6}\left( 4 \right)}+a_{16}*h_{16}*{OM}^{q_{6}\left( 5 \right)}+a_{17}*h_{17}*{DF}^{q_{6}\left( 6 \right)}}$$

$$-s_{7}+f_{1}\left( 7 \right)*{ds}_{1}\left( 7 \right)+f_{2}\left( 7 \right)*{ds}_{2}\left( 7 \right)*F$$

(E.8) Bacteria:

$$\frac{dB}{dt}= n_{20}*\frac{a_{20}*{CPOM}^{q_{7}\left( 3 \right)}*B}{1+a_{20}*h_{20}*{CPOM}^{q_{7}\left( 3 \right)}+a_{21}*h_{21}*{FPOM}^{q_{7}\left( 4 \right)}+a_{22}*h_{22}*{OM}^{q_{7}\left( 5 \right)}+a_{23}*h_{23}*{DF}^{q_{7}\left( 6 \right)}}$$

$$+n_{21}*\frac{a_{21}*{FPOM}^{q_{7}\left( 4 \right)}*B}{1+a_{20}*h_{20}*{CPOM}^{q_{7}\left( 3 \right)}+a_{21}*h_{21}*{FPOM}^{q_{7}\left( 4 \right)}+a_{22}*h_{22}*{OM}^{q_{7}\left( 5 \right)}+a_{23}*h_{23}*{DF}^{q_{7}\left( 6 \right)}}$$

$$+n_{22}*\frac{a_{22}*{OM}^{q_{7}\left( 5 \right)}*B}{1+a_{20}*h_{20}*{CPOM}^{q_{7}\left( 3 \right)}+a_{21}*h_{21}*{FPOM}^{q_{7}\left( 4 \right)}+a_{22}*h_{22}*{OM}^{q_{7}\left( 5 \right)}+a_{23}*h_{23}*{DF}^{q_{7}\left( 6 \right)}}$$

$$+n_{23}*\frac{a_{23}*{DF}^{q_{7}\left( 6 \right)}*B}{1+a_{20}*h_{20}*{CPOM}^{q_{7}\left( 3 \right)}+a_{21}*h_{21}*{FPOM}^{q_{7}\left( 4 \right)}+a_{22}*h_{22}*{OM}^{q_{7}\left( 5 \right)}+a_{23}*h_{23}*{DF}^{q_{7}\left( 6 \right)}}$$

$$-s_{8}+f_{1}\left( 8 \right)*{ds}_{1}\left( 8 \right)+f_{2}\left( 8 \right)*{ds}_{2}\left( 8 \right)*B$$

(E.9) Plant Detritus :

$$\frac{dPD}{dt}=s_{1}+f_{1}\left( 1 \right)*{ds}_{1}\left( 1 \right)+f_{2}\left( 1 \right)*{ds}_{2}\left( 1 \right)*P+\left( 1-n_{1} \right)*\frac{a_{1}*P^{q_{1}\left( 1 \right)}*G}{1+a_{1}*h_{1}*P^{q_{1}\left( 1 \right)}}+z_{1}$$

$$-n_{2}*\frac{a_{2}*{PD}^{q_{2}\left( 1 \right)}*D}{1+a_{2}*h_{2}*{PD}^{q_{2}\left( 1 \right)}+a_{3}*h_{3}*{AD}^{q_{2}\left( 2 \right)}}{-(1-n}_{2})*\frac{a_{2}*{PD}^{q_{2}\left( 1 \right)}*D}{1+a_{2}*h_{2}*{PD}^{q_{2}\left( 1 \right)}+a_{3}*h_{3}*{AD}^{q_{2}\left( 2 \right)}}$$

(E.10) Animal Detritus:

$$\frac{dAD}{dt}=s_{2}+f_{1}\left( 2 \right)*{ds}_{1}\left( 2 \right)+f_{2}\left( 2 \right)*{ds}_{2}\left( 2 \right)*G+s_{3}+f_{1}\left( 3 \right)*{ds}_{1}\left( 3 \right)+f_{2}\left( 3 \right)*{ds}_{2}\left( 3 \right)*D+s_{4}+f_{1}\left( 4 \right)*{ds}_{1}\left( 4 \right)+f_{2}\left( 4 \right)*{ds}_{2}\left( 4 \right)*Sh$$

$$+s_{5}+f_{1}\left( 5 \right)*{ds}_{1}\left( 5 \right)+f_{2}\left( 5 \right)*{ds}_{2}\left( 5 \right)*IP+s_{6}+f_{1}\left( 6 \right)*{ds}_{1}\left( 6 \right)+f_{2}\left( 6 \right)*{ds}_{2}\left( 6 \right)*FP$$

$$+\left( 1-n_{5} \right)*\frac{a_{5}*G^{q_{4}\left( 1 \right)}*IP}{1+a_{5}*h_{5}*G^{q_{4}\left( 1 \right)}+a_{6}*h_{6}*D^{q_{4}\left( 2 \right)}+a_{7}*h_{7}*{Sh}^{q_{4}\left( 3 \right)}}+\left( 1-n_{6} \right)*\frac{a_{6}*D^{q_{4}\left( 2 \right)}*IP}{1+a_{5}*h_{5}*G^{q_{4}\left( 1 \right)}+a_{6}*h_{6}*D^{q_{4}\left( 2 \right)}+a_{7}*h_{7}*{Sh}^{q_{4}\left( 3 \right)}}$$

$$+\left( 1-n_{7} \right)*\frac{a_{7}*{Sh}^{q_{4}\left( 3 \right)}*IP}{1+a_{5}*h_{5}*G^{q_{4}\left( 1 \right)}+a_{6}*h_{6}*D^{q_{4}\left( 2 \right)}+a_{7}*h_{7}*{Sh}^{q_{4}\left( 3 \right)}}$$

$$+\left( 1-n_{8} \right)*\frac{a_{8}*G^{q_{5}\left( 1 \right)}*FP}{1+a_{8}*h_{8}*G^{q_{5}\left( 1 \right)}+a_{9}*h_{9}*D^{q_{5}\left( 2 \right)}+a_{10}*h_{10}*{Sh}^{q_{5}\left( 3 \right)}+a_{11}*h_{11}*{IP}^{q_{5}\left( 4 \right)}}$$

$$+\left( 1-n_{9} \right)*\frac{a_{9}*D^{q_{5}\left( 2 \right)}*FP}{1+a_{8}*h_{8}*G^{q_{5}\left( 1 \right)}+a_{9}*h_{9}*D^{q_{5}\left( 2 \right)}+a_{10}*h_{10}*{Sh}^{q_{5}\left( 3 \right)}+a_{11}*h_{11}*{IP}^{q_{5}\left( 4 \right)}}$$

$$+\left( 1-n_{10} \right)*\frac{a_{10}*{Sh}^{q_{5}\left( 3 \right)}*FP}{1+a_{8}*h_{8}*G^{q_{5}\left( 1 \right)}+a_{9}*h_{9}*D^{q_{5}\left( 2 \right)}+a_{10}*h_{10}*{Sh}^{q_{5}\left( 3 \right)}+a_{11}*h_{11}*{IP}^{q_{5}\left( 4 \right)}}$$

$$+\left( 1-n_{11} \right)*\frac{a_{11}*{IP}^{q_{5}\left( 4 \right)}*FP}{1+a_{8}*h_{8}*G^{q_{5}\left( 1 \right)}+a_{9}*h_{9}*D^{q_{5}\left( 2 \right)}+a_{10}*h_{10}*{Sh}^{q_{5}\left( 3 \right)}+a_{11}*h_{11}*{IP}^{q_{5}\left( 4 \right)}}+z_{2}+z_{3}+z_{4}+z_{5}+z_{6}$$

$$-n_{3}*\frac{a_{3}*{AD}^{q_{2}\left( 2 \right)}*D}{1+a_{2}*h_{2}*{PD}^{q_{2}\left( 1 \right)}+a_{3}*h_{3}*{AD}^{q_{2}\left( 2 \right)}}{-(1-n}_{3})*\frac{a_{3}*{AD}^{q_{2}\left( 2 \right)}*D}{1+a_{2}*h_{2}*{PD}^{q_{2}\left( 1 \right)}+a_{3}*h_{3}*{AD}^{q_{2}\left( 2 \right)}}$$

(E.11) Coarse particulate organic matter:

$$\frac{dCPOM}{dt}=-n_{4}*\frac{a_{4}*{CPOM}^{q_{3}\left( 1 \right)}*Sh}{1+a_{4}*h_{4}*{CPOM}^{q_{3}\left( 1 \right)}}-\left( 1-n_{4} \right)*\frac{a_{4}*{CPOM}^{q_{3}\left( 1 \right)}*Sh}{1+a_{4}*h_{4}*{CPOM}^{q_{3}\left( 1 \right)}}$$

$$-n_{14}*\frac{a_{14}*{CPOM}^{q_{6}\left( 3 \right)}*F}{1+a_{14}*h_{14}*{CPOM}^{q_{6}\left( 3 \right)}+a_{15}*h_{15}*{FPOM}^{q_{6}\left( 4 \right)}+a_{16}*h_{16}*{OM}^{q_{6}\left( 5 \right)}+a_{17}*h_{17}*{DF}^{q_{6}\left( 6 \right)}}$$

$$-\left( 1-n_{14} \right)*\frac{a_{14}*{CPOM}^{q_{6}\left( 3 \right)}*F}{1+a_{14}*h_{14}*{CPOM}^{q_{6}\left( 3 \right)}+a_{15}*h_{15}*{FPOM}^{q_{6}\left( 4 \right)}+a_{16}*h_{16}*{OM}^{q_{6}\left( 5 \right)}+a_{17}*h_{17}*{DF}^{q_{6}\left( 6 \right)}}$$

$$-n_{20}*\frac{a_{20}*{CPOM}^{q_{7}\left( 3 \right)}*B}{1+a_{20}*h_{20}*{CPOM}^{q_{7}\left( 3 \right)}+a_{21}*h_{21}*{FPOM}^{q_{7}\left( 4 \right)}+a_{22}*h_{22}*{OM}^{q_{7}\left( 5 \right)}+a_{23}*h_{23}*{DF}^{q_{7}\left( 6 \right)}}$$

$$-\left( 1-n_{20} \right)*\frac{a_{20}*{CPOM}^{q_{7}\left( 3 \right)}*B}{1+a_{20}*h_{20}*{CPOM}^{q_{7}\left( 3 \right)}+a_{21}*h_{21}*{FPOM}^{q_{7}\left( 4 \right)}+a_{22}*h_{22}*{OM}^{q_{7}\left( 5 \right)}+a_{23}*h_{23}*{DF}^{q_{7}\left( 6 \right)}}$$

(E.12) Fine particulate organic matter:

$$\frac{dFPOM}{dt}=\left( 1-n_{4} \right)*\frac{a_{4}*{CPOM}^{q_{3}\left( 1 \right)}*Sh}{1+a_{4}*h_{4}*{CPOM}^{q_{3}\left( 1 \right)}}$$

$$+\left( 1-n_{14} \right)*\frac{a_{14}*{CPOM}^{q_{6}\left( 3 \right)}*F}{1+a_{14}*h_{14}*{CPOM}^{q_{6}\left( 3 \right)}+a_{15}*h_{15}*{FPOM}^{q_{6}\left( 4 \right)}+a_{16}*h_{16}*{OM}^{q_{6}\left( 5 \right)}+a_{17}*h_{17}*{DF}^{q_{6}\left( 6 \right)}}$$

$$+\left( 1-n_{20} \right)*\frac{a_{20}*{CPOM}^{q_{7}\left( 3 \right)}*B}{1+a_{20}*h_{20}*{CPOM}^{q_{7}\left( 3 \right)}+a_{21}*h_{21}*{FPOM}^{q_{7}\left( 4 \right)}+a_{22}*h_{22}*{OM}^{q_{7}\left( 5 \right)}+a_{23}*h_{23}*{DF}^{q_{7}\left( 6 \right)}}$$

$$-n_{15}*\frac{a_{15}*{FPOM}^{q_{6}\left( 4 \right)}*F}{1+a_{14}*h_{14}*{CPOM}^{q_{6}\left( 3 \right)}+a_{15}*h_{15}*{FPOM}^{q_{6}\left( 4 \right)}+a_{16}*h_{16}*{OM}^{q_{6}\left( 5 \right)}+a_{17}*h_{17}*{DF}^{q_{6}\left( 6 \right)}}$$

$$-\left( 1-n_{15} \right)*\frac{a_{15}*{FPOM}^{q_{6}\left( 4 \right)}*F}{1+a_{14}*h_{14}*{CPOM}^{q_{6}\left( 3 \right)}+a_{15}*h_{15}*{FPOM}^{q_{6}\left( 4 \right)}+a_{16}*h_{16}*{OM}^{q_{6}\left( 5 \right)}+a_{17}*h_{17}*{DF}^{q_{6}\left( 6 \right)}}$$

$$-n_{21}*\frac{a_{21}*{FPOM}^{q_{7}\left( 4 \right)}*B}{1+a_{20}*h_{20}*{CPOM}^{q_{7}\left( 3 \right)}+a_{21}*h_{21}*{FPOM}^{q_{7}\left( 4 \right)}+a_{22}*h_{22}*{OM}^{q_{7}\left( 5 \right)}+a_{23}*h_{23}*{DF}^{q_{7}\left( 6 \right)}}$$

$$-\left( 1-n_{21} \right)*\frac{a_{21}*{FPOM}^{q_{7}\left( 4 \right)}*B}{1+a_{20}*h_{20}*{CPOM}^{q_{7}\left( 3 \right)}+a_{21}*h_{21}*{FPOM}^{q_{7}\left( 4 \right)}+a_{22}*h_{22}*{OM}^{q_{7}\left( 5 \right)}+a_{23}*h_{23}*{DF}^{q_{7}\left( 6 \right)}}$$

(E.13) Organic matter:

$$\frac{dOM}{dt}={(1-n}_{2})*\frac{a_{2}*{PD}^{q_{2}\left( 1 \right)}*D}{1+a_{2}*h_{2}*{PD}^{q_{2}\left( 1 \right)}+a_{3}*h_{3}*{AD}^{q_{2}\left( 2 \right)}}$$

$$+{(1-n}_{3})*\frac{a_{3}*{AD}^{q_{2}\left( 2 \right)}*D}{1+a_{2}*h_{2}*{PD}^{q_{2}\left( 1 \right)}+a_{3}*h_{3}*{AD}^{q_{2}\left( 2 \right)}}$$

$$+\left( 1-n_{15} \right)*\frac{a_{15}*{FPOM}^{q_{6}\left( 4 \right)}*F}{1+a_{14}*h_{14}*{CPOM}^{q_{6}\left( 3 \right)}+a_{15}*h_{15}*{FPOM}^{q_{6}\left( 4 \right)}+a_{16}*h_{16}*{OM}^{q_{6}\left( 5 \right)}+a_{17}*h_{17}*{DF}^{q_{6}\left( 6 \right)}}$$

$$+\left( 1-n_{17} \right)*\frac{a_{17}*{DF}^{q_{6}\left( 6 \right)}*F}{1+a_{14}*h_{14}*{CPOM}^{q_{6}\left( 3 \right)}+a_{15}*h_{15}*{FPOM}^{q_{6}\left( 4 \right)}+a_{16}*h_{16}*{OM}^{q_{6}\left( 5 \right)}+a_{17}*h_{17}*{DF}^{q_{6}\left( 6 \right)}}$$

$$+\left( 1-n_{21} \right)*\frac{a_{21}*{FPOM}^{q_{7}\left( 4 \right)}*B}{1+a_{20}*h_{20}*{CPOM}^{q_{7}\left( 3 \right)}+a_{21}*h_{21}*{FPOM}^{q_{7}\left( 4 \right)}+a_{22}*h_{22}*{OM}^{q_{7}\left( 5 \right)}+a_{23}*h_{23}*{DB}^{q_{7}\left( 6 \right)}}$$

$$+\left( 1-n_{23} \right)*\frac{a_{23}*{DB}^{q_{7}\left( 6 \right)}*B}{1+a_{20}*h_{20}*{CPOM}^{q_{7}\left( 3 \right)}+a_{21}*h_{21}*{FPOM}^{q_{7}\left( 4 \right)}+a_{22}*h_{22}*{OM}^{q_{7}\left( 5 \right)}+a_{23}*h_{23}*{DB}^{q_{7}\left( 6 \right)}}$$

$$-n_{16}*\frac{a_{16}*{OM}^{q_{6}\left( 5 \right)}*F}{1+a_{14}*h_{14}*{CPOM}^{q_{6}\left( 3 \right)}+a_{15}*h_{15}*{FPOM}^{q_{6}\left( 4 \right)}+a_{16}*h_{16}*{OM}^{q_{6}\left( 5 \right)}+a_{17}*h_{17}*{DF}^{q_{6}\left( 6 \right)}}$$

$$-\left( 1-n_{16} \right)*\frac{a_{16}*{OM}^{q_{6}\left( 5 \right)}*F}{1+a_{14}*h_{14}*{CPOM}^{q_{6}\left( 3 \right)}+a_{15}*h_{15}*{FPOM}^{q_{6}\left( 4 \right)}+a_{16}*h_{16}*{OM}^{q_{6}\left( 5 \right)}+a_{17}*h_{17}*{DF}^{q_{6}\left( 6 \right)}}$$

$$-n_{22}*\frac{a_{22}*{OM}^{q_{7}\left( 5 \right)}*B}{1+a_{20}*h_{20}*{CPOM}^{q_{7}\left( 3 \right)}+a_{21}*h_{21}*{FPOM}^{q_{7}\left( 4 \right)}+a_{22}*h_{22}*{OM}^{q_{7}\left( 5 \right)}+a_{23}*h_{23}*{DB}^{q_{7}\left( 6 \right)}}$$

$$-(1-n_{22})*\frac{a_{22}*{OM}^{q_{7}\left( 5 \right)}*B}{1+a_{20}*h_{20}*{CPOM}^{q_{7}\left( 3 \right)}+a_{21}*h_{21}*{FPOM}^{q_{7}\left( 4 \right)}+a_{22}*h_{22}*{OM}^{q_{7}\left( 5 \right)}+a_{23}*h_{23}*{DB}^{q_{7}\left( 6 \right)}}$$

(E.14) Dead Fungi:

$$\frac{dDF}{dt}=s_{7}+f_{1}\left( 7 \right)*{ds}_{1}\left( 7 \right)+f_{2}\left( 7 \right)*{ds}_{2}\left( 7 \right)*F+z_{7}$$

$$-n_{17}*\frac{a_{17}*{DF}^{q_{6}\left( 6 \right)}*F}{1+a_{14}*h_{14}*{CPOM}^{q_{6}\left( 3 \right)}+a_{15}*h_{15}*{FPOM}^{q_{6}\left( 4 \right)}+a_{16}*h_{16}*{OM}^{q_{6}\left( 5 \right)}+a_{17}*h_{17}*{DF}^{q_{6}\left( 6 \right)}}$$

$$-(1-n_{17})*\frac{a_{17}*{DF}^{q_{6}\left( 6 \right)}*F}{1+a_{14}*h_{14}*{CPOM}^{q_{6}\left( 3 \right)}+a_{15}*h_{15}*{FPOM}^{q_{6}\left( 4 \right)}+a_{16}*h_{16}*{OM}^{q_{6}\left( 5 \right)}+a_{17}*h_{17}*{DF}^{q_{6}\left( 6 \right)}}$$

(E.15) Dead Bacteria:

$$\frac{dDB}{dt}=s_{8}+f_{1}\left( 8 \right)*{ds}_{1}\left( 8 \right)+f_{2}\left( 8 \right)*{ds}_{2}\left( 8 \right)*B+z_{8}$$

$$-n_{23}*\frac{a_{23}*{DB}^{q_{7}\left( 6 \right)}*B}{1+a_{20}*h_{20}*{CPOM}^{q_{7}\left( 3 \right)}+a_{21}*h_{21}*{FPOM}^{q_{7}\left( 4 \right)}+a_{22}*h_{22}*{OM}^{q_{7}\left( 5 \right)}+a_{23}*h_{23}*{DB}^{q_{7}\left( 6 \right)}}$$

$$-\left( 1-n_{23} \right)*\frac{a_{23}*{DB}^{q_{7}\left( 6 \right)}*B}{1+a_{20}*h_{20}*{CPOM}^{q_{7}\left( 3 \right)}+a_{21}*h_{21}*{FPOM}^{q_{7}\left( 4 \right)}+a_{22}*h_{22}*{OM}^{q_{7}\left( 5 \right)}+a_{23}*h_{23}*{DB}^{q_{7}\left( 6 \right)}}$$

(E.16) Nutrient:

$$\frac{dN}{dt}=gamma*\left( 1-n_{16} \right)*\frac{a_{16}*{OM}^{q_{6}\left( 5 \right)}*F}{1+a_{14}*h_{14}*{CPOM}^{q_{6}\left( 3 \right)}+a_{15}*h_{15}*{FPOM}^{q_{6}\left( 4 \right)}+a_{16}*h_{16}*{OM}^{q_{6}\left( 5 \right)}+a_{17}*h_{17}*{DF}^{q_{6}\left( 6 \right)}}$$

$$+gamma*(1-n_{22})*\frac{a_{22}*{OM}^{q_{7}\left( 5 \right)}*B}{1+a_{20}*h_{20}*{CPOM}^{q_{7}\left( 3 \right)}+a_{21}*h_{21}*{FPOM}^{q_{7}\left( 4 \right)}+a_{22}*h_{22}*{OM}^{q_{7}\left( 5 \right)}+a_{23}*h_{23}*{DB}^{q_{7}\left( 6 \right)}}$$

$$-\frac{gamma*r_{1}*P*N}{k_{1}+N}$$

**Table B1**: Parameters

| **Parameter** | **Interpretation** | **Unit** | **Value** |
| --- | --- | --- | --- |
| r1 | growth rate | d^-1^ | 0.45**^a^** |
| gamma |  |  | 0.4 |
| k1 | half saturation |  | 0.5**^b^** |
| n1 | conversion efficiency from Periphyton to Grazer |  | 0.6**^a^** |
| n2 | conversion efficiency from plant detritus to Detritivore |  | 0.6**^a^** |
| n3 | conversion efficiency from animal detritus to Detritivore |  | 0.6**^a^** |
| n4 | conversion efficiency from CPOM to Shredder |  | 0.6**^a^** |
| n5 | conversion efficiency from Grazer to invert. Predator |  | 0.15 |
| n6 | conversion efficiency from Detritivore to invert. Predator |  | 0.15 |
| n7 | conversion efficiency from Shredder to invert. Predator |  | 0.15 |
| n8 | conversion efficiency from Grazer to Fish |  | 0.1**^a^** |
| n9 | conversion efficiency from Detritivore to Fish |  | 0.1**^a^** |
| n10 | conversion efficiency from Shredder to Fish |  | 0.1**^a^** |
| n11 | conversion efficiency from invert. Predator to Fish |  | 0.1**^a^** |
| n14 | conversion efficiency from CPOM to Fungi |  | 0.5 |
| n15 | conversion efficiency from FPOM to Fungi |  | 0.5 |
| n16 | conversion efficiency from OM to Fungi |  | 0.4 |
| n17 | conversion efficiency from dead Fungi to Fungi |  | 0.4 |
| n20 | conversion efficiency from CPOM to Bacteria |  | 0.5 |
| n21 | conversion efficiency from FPOM to Bacteria |  | 0.5 |
| n22 | conversion efficiency from OM to Bacteria |  | 0.4 |
| n23 | conversion efficiency from dead Bacteria to Bacteria |  | 0.4 |
| h1 | handling time Grazer on Periphyton | d^-1^*mg^-1^*mg^-1^ | 0.87**^c^** |
| h2 | handling time Detritivore on plant detritus | d^-1^*mg^-1^*mg^-1^ | 0.6 |
| h3 | handling time Detritivore on animal detritus | d^-1^*mg^-1^*mg^-1^ | 0.6 |
| h4 | handling time Shredder on CPOM | d^-1^*mg^-1^*mg^-1^ | 0.7 |
| h5 | handling time invert. Predator on Grazer | d^-1^*mg^-1^*mg^-1^ | 0.2**^d^** |
| h6 | handling time invert. Predator on Detritivore | d^-1^*mg^-1^*mg^-1^ | 0.2**^d^** |
| h7 | handling time invert. Predator on Shredder | d^-1^*mg^-1^*mg^-1^ | 0.2**^d^** |
| h8 | handling time Fish on Grazer | d^-1^*mg^-1^*mg^-1^ | 0.01**^d^** |
| h9 | handling time Fish on Detritivore | d^-1^*mg^-1^*mg^-1^ | 0.01**^d^** |
| h10 | handling time Fish on Shredder | d^-1^*mg^-1^*mg^-1^ | 0.01**^d^** |
| h11 | handling time Fish on invert. Predator | d^-1^*mg^-1^*mg^-1^ | 0.01**^d^** |
| h14 | handling time Fungi on CPOM | d^-1^*mg^-1^*mg^-1^ | 9 |
| h15 | handling time Fungi on FPOM | d^-1^*mg^-1^*mg^-1^ | 6 |
| h16 | handling time Fungi on OM | d^-1^*mg^-1^*mg^-1^ | 3 |
| h17 | handling time Fungi on dead Fungi | d^-1^*mg^-1^*mg^-1^ | 3 |
| h20 | handling time Bacteria on CPOM | d^-1^*mg^-1^*mg^-1^ | 9 |
| h21 | handling time Bacteria on FPOM | d^-1^*mg^-1^*mg^-1^ | 6 |
| h22 | handling time Bacteria on OM | d^-1^*mg^-1^*mg^-1^ | 3 |
| h23 | handling time Bacteria on dead Bacteria | d^-1^*mg^-1^*mg^-1^ | 3 |
| a1 | attack rate Grazer on Periphyton | L*mg^-1^ *mg^-1^ | 1.2 |
| a2 | attack rate Detritivore on plant detritus | L*mg^-1^ *mg^-1^ | 0.7 |
| a3 | attack rate Detritivore on animal detritus | L*mg^-1^ *mg^-1^ | 0.7 |
| a4 | attack rate Shredder on CPOM | L*mg^-1^ *mg^-1^ | 0.7 |
| a5 | attack rate invert. Predator on Grazer | L*mg^-1^ *mg^-1^ | 0.17**^d^** |
| a6 | attack rate invert. Predator on Detritivore | L*mg^-1^ *mg^-1^ | 0.17**^d^** |
| a7 | attack rate invert. Predator on Shredder | L*mg^-1^ *mg^-1^ | 0.17**^d^** |
| a8 | attack rate Fish on Grazer | L*mg^-1^ *mg^-1^ | 0.02**^d^** |
| a9 | attack rate Fish on Detritivore | L*mg^-1^ *mg^-1^ | 0.02**^d^** |
| a10 | attack rate Fish on Shredder | L*mg^-1^ *mg^-1^ | 0.02**^d^** |
| a11 | attack rate Fish on invert. Predator | L*mg^-1^ *mg^-1^ | 0.02**^d^** |
| a14 | attack rate Fungi on CPOM | L*mg^-1^ *mg^-1^ | 0.6 |
| a15 | attack rate Fungi on FPOM | L*mg^-1^ *mg^-1^ | 0.4 |
| a16 | attack rate Fungi on OM | L*mg^-1^ *mg^-1^ | 0.4 |
| a17 | attack rate Fungi on dead Fungi | L*mg^-1^ *mg^-1^ | 0.7 |
| a20 | attack rate Bacteria on CPOM | L*mg^-1^ *mg^-1^ | 0.4 |
| a21 | attack rate Bacteria onFPOM | L*mg^-1^ *mg^-1^ | 0.6 |
| a22 | attack rate Bacteria on OM | L*mg^-1^ *mg^-1^ | 0.6 |
| a23 | attack rate Bacteria on dead Bacteria | L*mg^-1^ *mg^-1^ | 0.7 |
| s1 | death rate Periphyton | d^-1^ | 0.09^a^ |
| s2 | death rate Grazer | d^-1^ | 0.002^a^ |
| s3 | death rate Detritivore | d^-1^ | 0.0035^a^ |
| s4 | death rate Shredder | d^-1^ | 0.002^a^ |
| s5 | death rate invert. Predator | d^-1^ | 0.0025^a^ |
| s6 | death rate Fish | d^-1^ | 0.004 |
| s7 | death rate Fungi | d^-1^ | 0.04 |
| s8 | death rate Bacteria | d^-1^ | 0.05**^e^** |
| x_0_ | initial density Periphyton | mg/L | 0.24858424 |
| x_0_ | initial density Grazer | mg/L | 0.29225558 |
| x_0_ | initial density Detritivore | mg/L | 0.67901786 |
| x_0_ | initial density Shredder | mg/L | 0.01111790 |
| x_0_ | initial density invert. Predator | mg/L | 0.73986864 |
| x_0_ | initial density Fish | mg/L | 0.69325469 |
| x_0_ | initial density Fungi | mg/L | 1.32170667 |
| x_0_ | initial density Bacteria | mg/L | 0.00542806 |
| x_0_ | initial density Plant detritus | mg/L | 0.26628438 |
| x_0_ | initial density Animal detritus | mg/L | 0.42796874 |
| x_0_ | initial density CPOM | mg/L | 25.00170936 |
| x_0_ | initial density FPOM | mg/L | 0.00326880 |
| x_0_ | initial density OM | mg/L | 0.44665074 |
| x_0_ | initial density dead Fungi | mg/L | 0.28361102 |
| x_0_ | initial density dead Bacteria | mg/L | 0.33634156 |
| x_0_ | initial density Nutrient | mg/L | 10.81806568 |
| Threshold | Minimum viable population density | mg/L | 0.001 |
| rd | reintroduction densities Grazer | mg/L | 0.03028799;  0.27259187;  0.3287986;  0.33316784 |
| rd | reintroduction densities Shredder | mg/L | 0.00238637;  0.02147737;  0.0238674;  0.02625011 |
| rd | reintroduction densities invert. Predator | mg/L | 0.06596450;  0.59368049;  0.65964499;  0.72560949 |
| rd | reintroduction densities Fish | mg/L | 0.09847580;  0.88628219:  0.98475799;  1.08323379 |

**^a^** Jorgensen et al., 1991; **^b^** DeAngelis, 2003; **^c^** Ciric et al., 2012; **^d^** Uiterwaal et al., 2018; **^e^** Servais et al., 1985

**Table B2**: Matlab settings

| **Settings** | **Value** | **Interpretation** |
| --- | --- | --- |
| ODE45 |  | Solver for ordinary differential equations (ODE) |
| Non-negative |  |  |
| Rel-Tol | 1.00E-03 | Relative tolerance |
| Abs-Tol | 1.00E-06 | Absolute tolerance |

**References**

Ciric, C., Ciffroy, P., Charles, S., 2012. Use of sensitivity analysis to identify influential and non-influential parameters within an aquatic ecosystem model. Ecol. Model. 246, 119–130. https://doi.org/10.1016/j.ecolmodel.2012.06.024

DeAngelis, D.L., 2003. Mathematical modeling relevant to closed artificial ecosystems. Adv. Space Res. 31, 1657–1665. https://doi.org/10.1016/S0273-1177(03)80012-1

Jorgensen, S.E., Nielsen, S.N., Jorgensen, L.A., 1991. Handbook of ecological parameters and ecotoxicology. Elsevier Editora.

Servais, P., Billen, G., Rego, J.V., 1985. Rate of Bacterial Mortality in Aquatic Environments. Appl. Environ. Microbiol. 49, 1448–1454. https://doi.org/10.1128/aem.49.6.1448-1454.1985

Uiterwaal, S.F., Lagerstrom, I.T., Lyon, S.R., DeLong, J.P., 2018. Data paper: FoRAGE (Functional Responses from Around the Globe in all Ecosystems) database: a compilation of functional responses for consumers and parasitoids (preprint). Ecology. https://doi.org/10.1101/503334
