## Supplementary material for "Putting the Asymmetric Response Concept to the test: modeling multiple stressor exposure and release in a stream food web": Supp Appendix C

**Table C1**. Results of each single and multiple stressor scenario in terms of extinctions. Groups were listed in order of extinction along with the time step (ts) at which the extinction happened.

| **Scenario no.** | **Salinity level** | **Temperature** | **Sequence** | **Extinctions** |
| --- | --- | --- | --- | --- |
| 1 | 600 mg/L | - | single | - |
| 2 | 1500 mg/L | - | single | Shredder ts 927 |
| 3 | - | 18.5°C | single | Shredder ts 946 |
| 4 | - | 25°C | single | Shredder ts 919  Fish ts 923  Grazer ts 926  invert. Predator ts 926 |
| 5 | 600 mg/L | 18.5°C | multiple | Shredder ts 925 |
| 6 | 1500 mg/L | 18.5°C | multiple | Shredder ts 921 |
| 7 | 600 mg/L | 25°C | multiple | Shredder ts 918  Grazer ts 923  invert. Predator ts 924  Fish ts 925 |
| 8 | 1500 mg/L | 25°C | multiple | Shredder ts 917  Grazer ts 922  invert. Predator ts 922  Fish ts 923 |
| 9 | 200 mg/L | - | single | - |
| 10 | - | 22°C | single | Shredder ts 921  Fish ts 927 |
| 11 | 5300 mg/L | - | single | Shredder ts 918  Grazer ts 923  invert. Predator ts 923 |

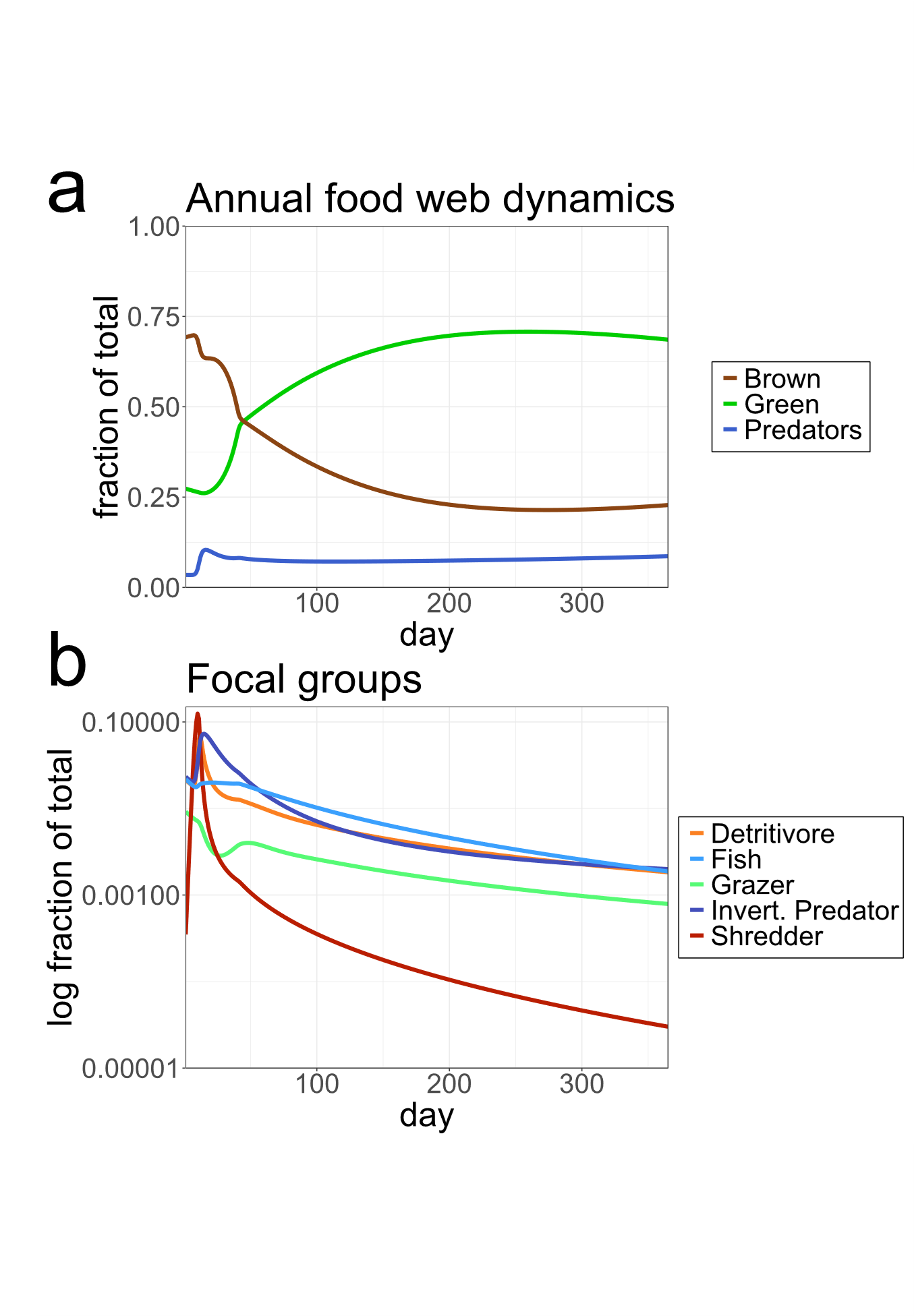

**Figure C1**. Annual dynamics of the fractions of the green, brown, and predator parts of the food web (**a**), as well as the annual dynamics of the densities of the focal functional groups detritivore, fish, grazer, invertebrate predator, and shredder (**b**), within an unstressed system. Depicted are the dynamics for one year, which repeat cyclically in subsequent years in an unstressed system. The entire food web is driven by an annual CPOM input, which results in peaks of the brown food web at the beginning of each simulated year. Green and brown parts alternate dominating the system. The green part consists of the nutrient, grazer, and periphyton; the brown part contains the CPOM, FPOM, OM, shredder, detritivore, bacteria, fungi, plant detritus, animal detritus, dead bacteria, and dead fungi; the predators that feed on both parts are fish and invertebrate predators.

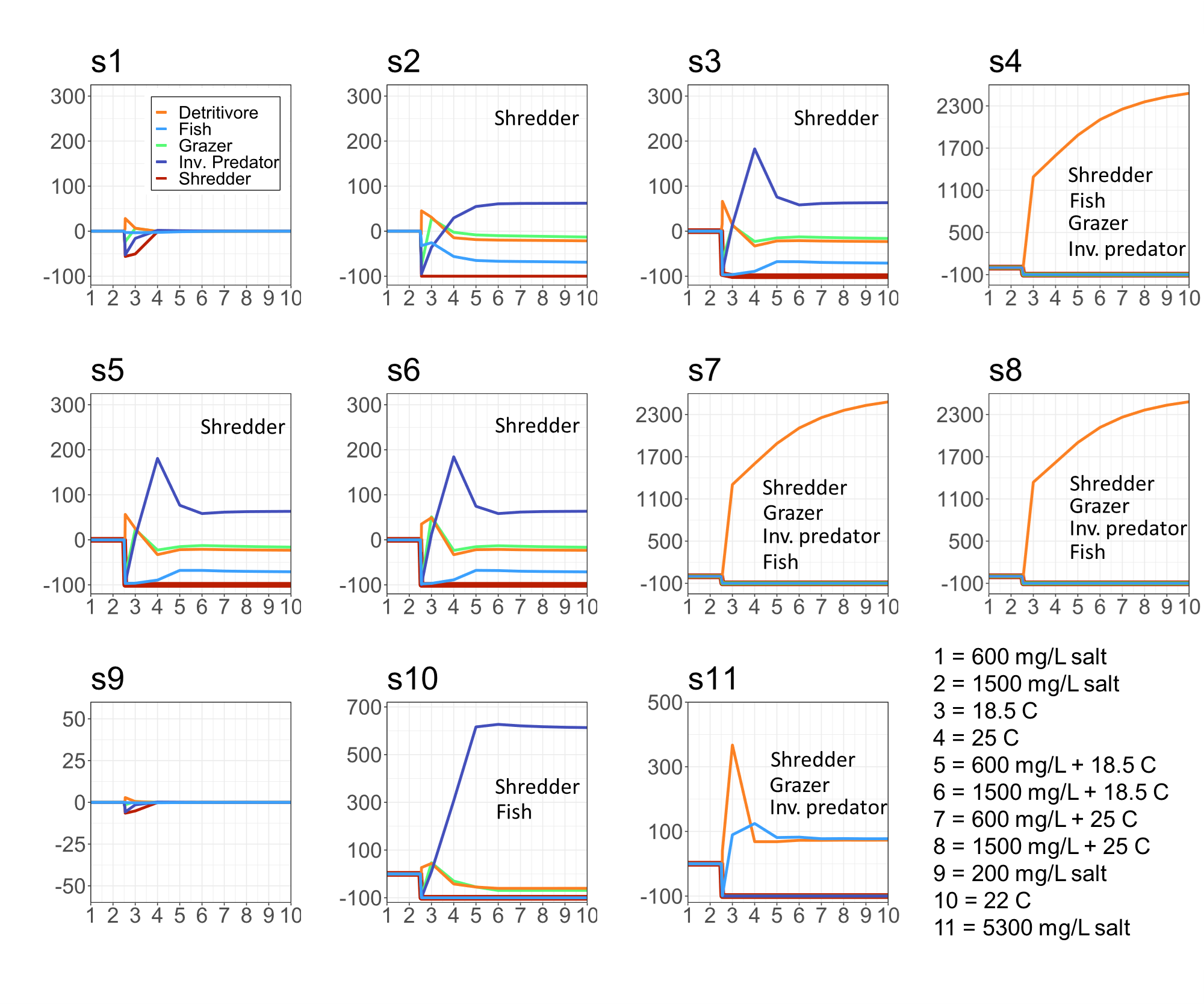

**Figure C2.** Results of all tested scenarios. Simulation year on the x-axis and % difference from baseline on the y-axis. Where applicable, extinct groups are labelled in order of extinction.

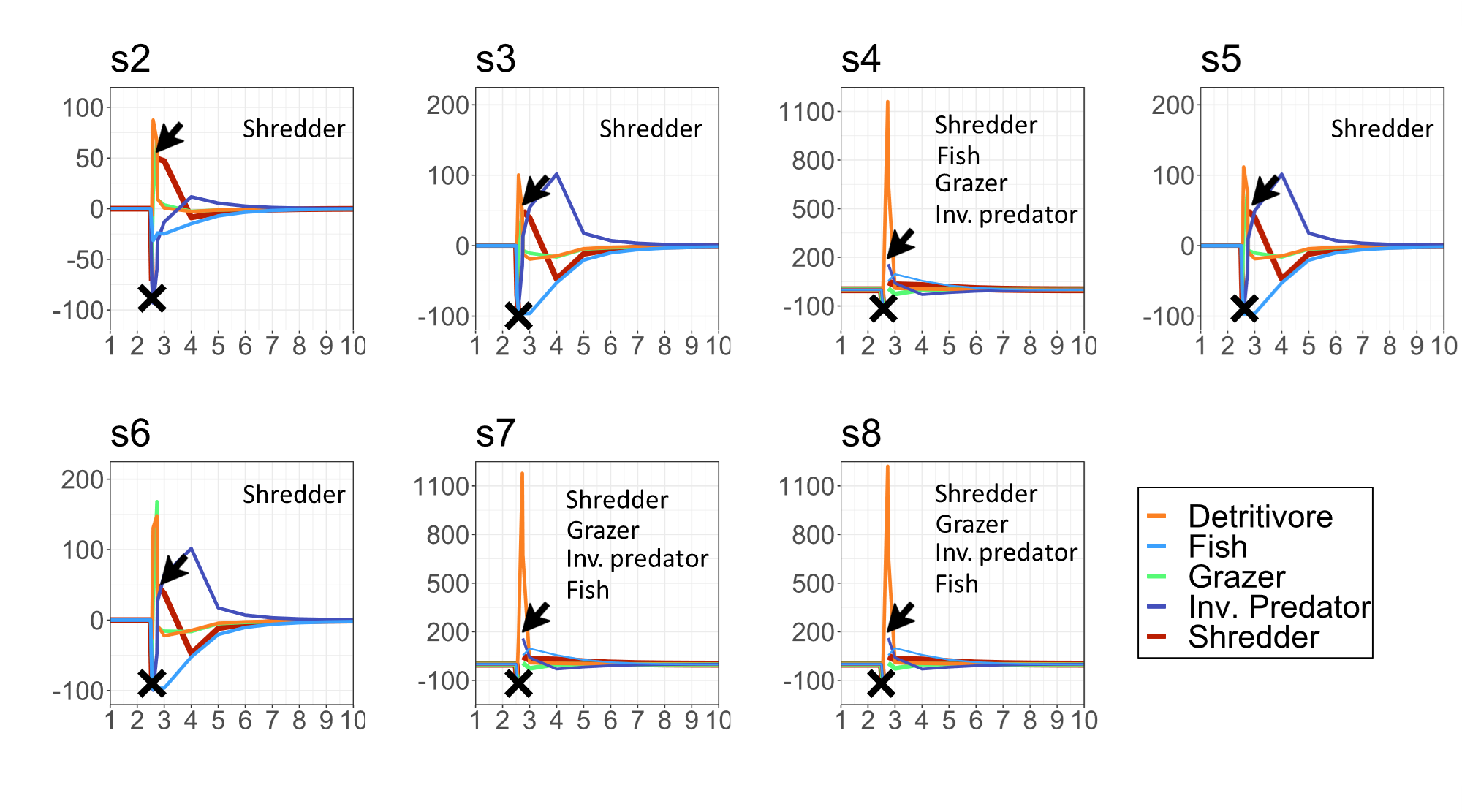

**Figure C3**. Results of reintroductions of all extinct functional groups after the stressor phase at time step 1000. All groups recolonize successfully.

**Table C2**. Spearman’s 𝜌 for the correlation comparing laboratory thermal tolerance ranking with decrease rate during stressor phase, laboratory thermal tolerance ranking with effect after the stressor phase (days to extinction), and decrease rate during the stressor phase with effect after the stressor phase (days to extinction). Correlations for days to extinction were only calculated when 3 or more extinctions occurred.

|  | Spearman’s 𝜌 between rankings | | |
| --- | --- | --- | --- |
| Scenario | Laboratory and stressor phase | Laboratory and after stressor phase | During and after stressor phase |
| 1 | -0.7 | NA | NA |
| 2 | -0.7 | NA | NA |
| 3 | 0.4 | NA | NA |
| 4 | 0.4 | 0.15 | 0.87 |
| 5 | 0.2 | NA | NA |
| 6 | -0.5 | NA | NA |
| 7 | 0.2 | -0.6 | 0.6 |
| 8 | -0.1 | -0.56 | 0.82 |
| 9 | -0.7 | NA | NA |
| 10 | 0.4 | NA | NA |
| 11 | -0.5 | -0.56 | 0.97 |

All results of the reintroduction scenarios which were implemented following the stressor phase can be seen in Figures C4 - C7.

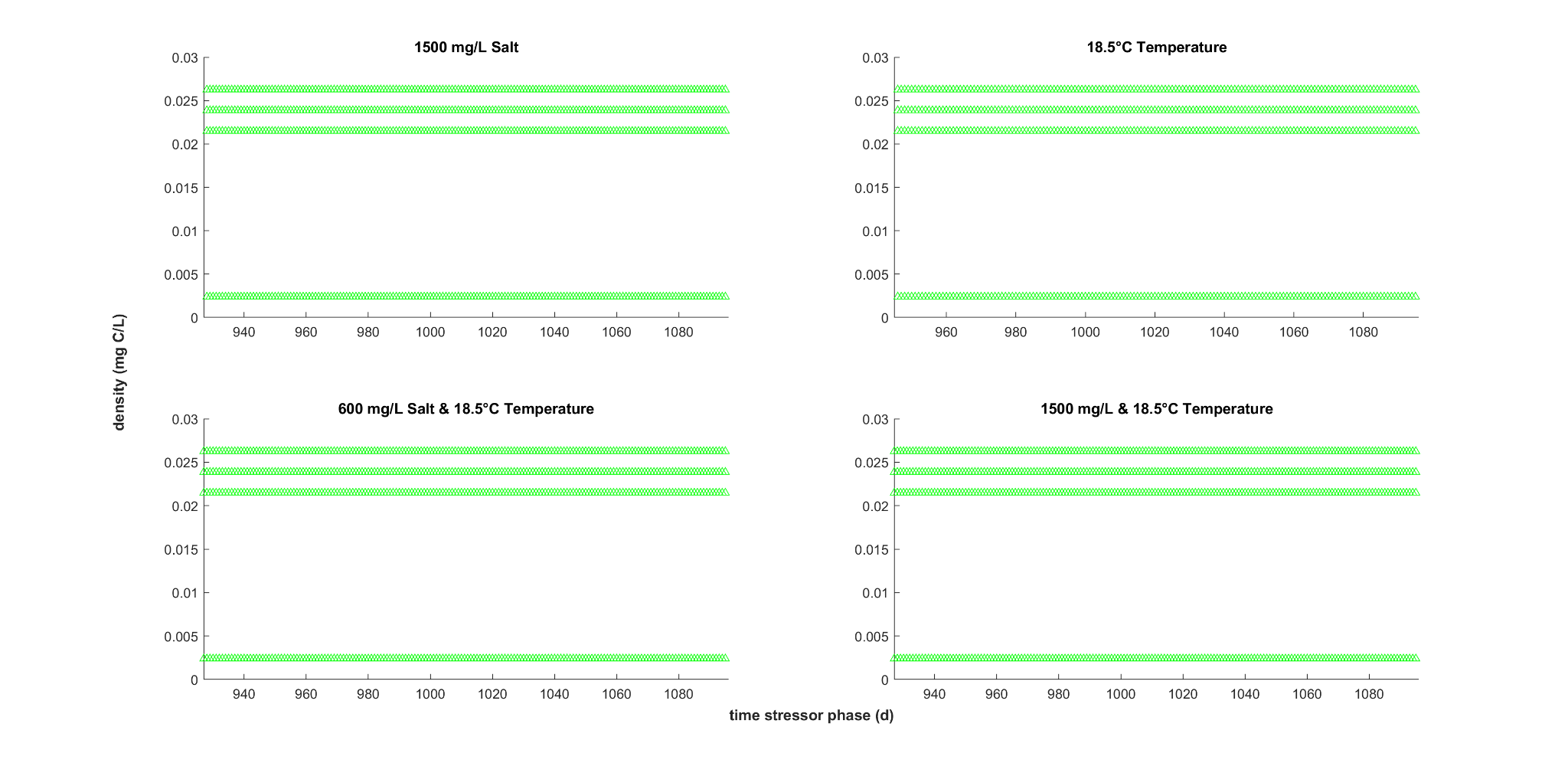

**Figure C4**. Reintroduction of the shredder after single and multiple stressor scenarios for scenario 2: 1500 mg/L salt between time steps 928 and 1095; scenario 3: 18.5°C temperature between time steps 947 and 1095 scenario type; and scenario 5: 600 mg/L salt & 18.5°C temperature and scenario 6: 1500 mg/L salt & 18.5°C temperature between time steps 927 and 1095. Green marks the successful reintroduction of the shredder.

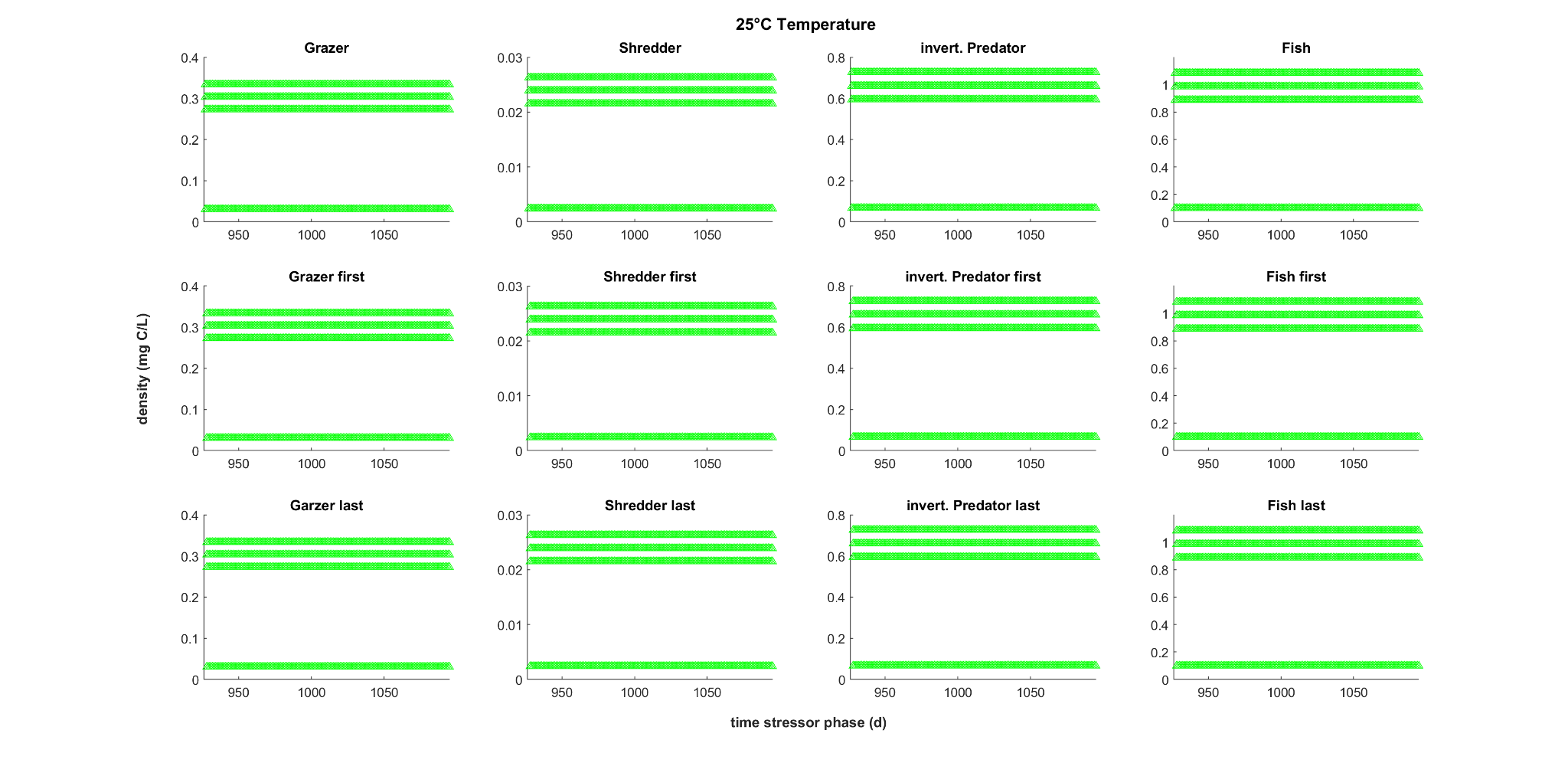

**Figure C5**. Reintroduction of all extinct groups either simultaneously (upper row) or sequentially, after the single stressor scenario 4: 25°C temperature between time steps 927 and 1095. In each column the focus was on a different group which was reintroduced with four different densities while the other groups were reintroduced with only the pre-stressor density. In the upper row all groups were reintroduced simultaneously. Sequential reintroductions with either the focal species reintroduced first at time step 927 and all others between time steps 928 and 1095 (middle row) or last when the other groups were reintroduced at time step 927 and the focal group between time steps 928 and 1095 (lower row). Green marks the successful reintroduction of all reintroduced groups.

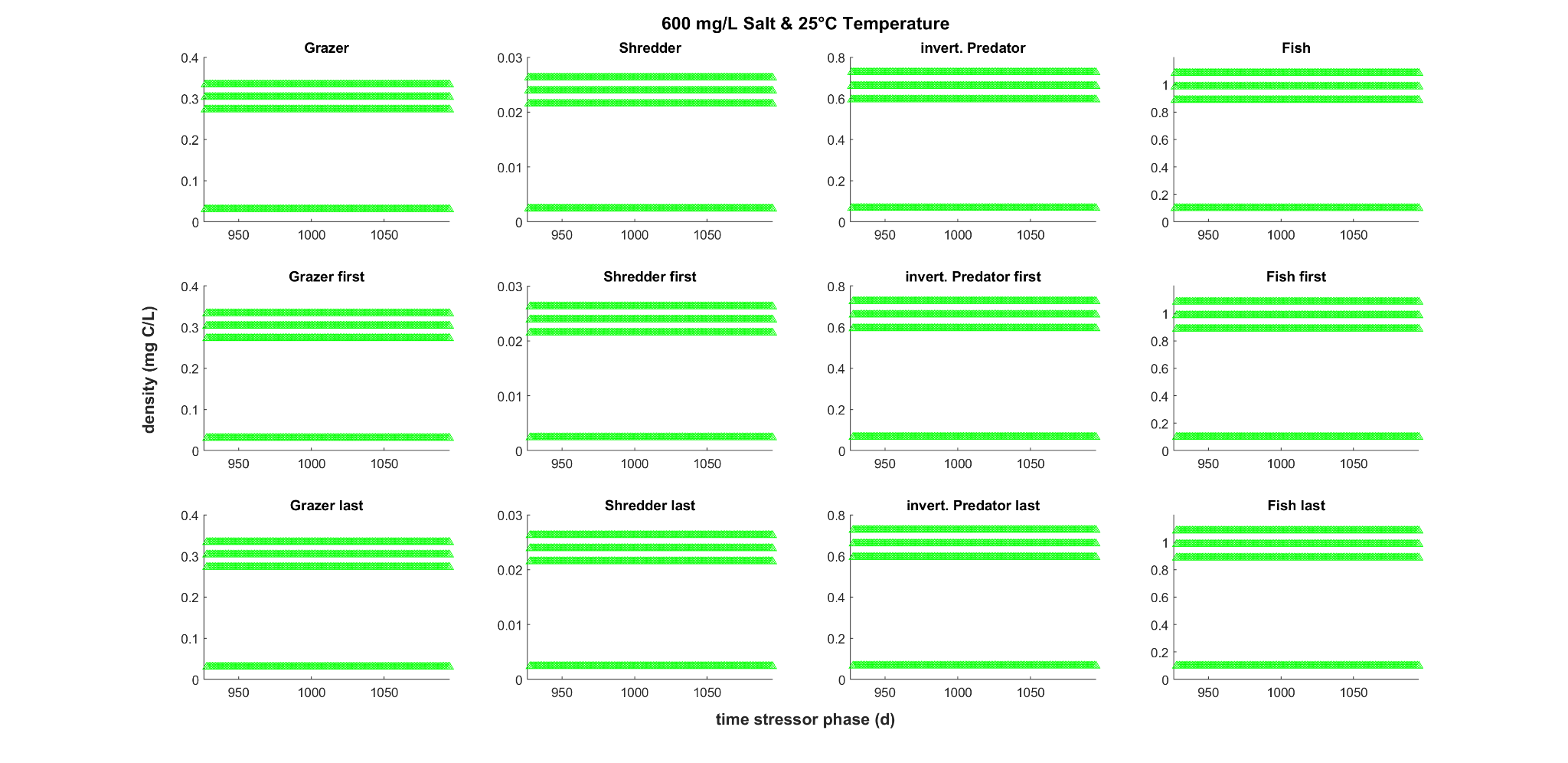

**Figure C6**. Reintroduction of all extinct groups either simultaneously (upper row) or sequentially, after multiple stressor scenario 7: 600 mg/L salt & 25°C temperature between time steps 927 and 1095. In each column the focus was on a different group that was reintroduced with four different densities while the other groups were reintroduced with only the pre-stressor density. In the upper row all groups were reintroduced simultaneously. Sequential reintroductions with either the focal group reintroduced first at time step 927 and all others between time steps 928 and 1095 (middle row) or last when the other groups were reintroduced at time step 297 and the focal group between time steps 928 and 1095 (lower row). Green marks the successful reintroduction of all reintroduced groups.

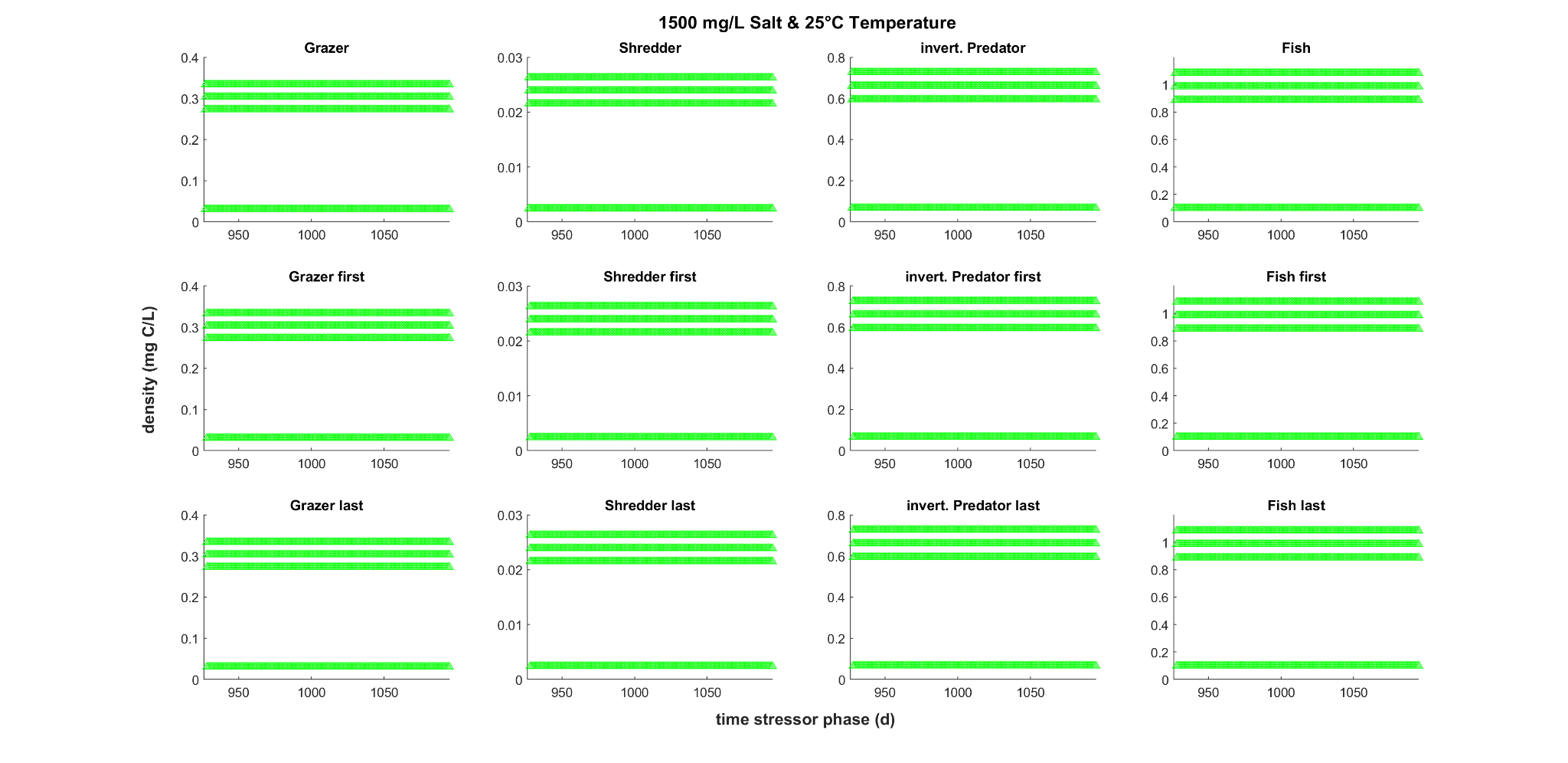

**Figure C7**. Reintroduction of all extinct groups either simultaneously (upper row) or sequentially, after multiple stressor scenario 8: 1500 mg/L salt & 25°C temperature between time steps 927 and 1095. In each column the focus was on a different group that was reintroduced with four different densities while the other groups were reintroduced with only the pre-stressor density. In the upper row all groups were reintroduced simultaneously. Sequential reintroductions with either the focal group reintroduced first at time step 927 and all others between time steps 928 and 1095 (middle row) or last when the other groups were reintroduced at time step 297 and the focal group between time steps 928 and 1095 (lower row). Green marks the successful reintroduction of all reintroduced groups.
